# Bacterial Swarmers exhibit a Protective Response to Intestinal Stress

**DOI:** 10.1101/759886

**Authors:** Weijie Chen, Arpan De, Hao Li, Justin R. Wright, Regina Lamendella, Dana J. Lukin, Wendy Szymczak, Katherine Sun, Libusha Kelly, Subho Ghosh, Daniel B. Kearns, Zhen He, Christian Jobin, Xiaoping Luo, Arjun Byju, Shirshendu Chatterjee, Beng San Yeoh, Matam Vijay-Kumar, Jay X. Tang, Sridhar Mani

## Abstract

Bacterial swarming, a collective movement on a surface, has rarely been associated with human pathophysiology. Here, we report for the first time that bacterial swarmers are associated with protection against intestinal inflammation. We show that bacterial swarmers are highly predictive of intestinal stress in mice and humans. We isolated a novel *Enterobacter* swarming strain, SM3, from mouse feces. SM3 and other known commensal swarmers contrast to their respective swarming-deficient, but swimming-competent isogenic strains abrogated intestinal inflammation in mice. Treatment of colitic mice with SM3, but not its mutants, enriched beneficial fecal anaerobes belonging to the family, Bacteroidales S24-7. We observed SM3 swarming associated pathways in the *in vivo* fecal metatranscriptomes. *In vitro* growth of S24-7 was enriched in presence of SM3 or its mutants conjecturing that bacterial swarming *in vivo* might influence SM3’s access to S24-7 in the intestines. Overall, our work identifies a new paradigm in which intestinal stress allows for the emergence of swarming bacteria, which can counterintuitively heal intestinal inflammation.

## Introduction

Bacterial motility is essential in mucosal colonization and has long been associated with virulence and pathogenesis (Stanton and Savage, 1984; Wiles et al., 2020). Intestinal inflammation, such as inflammatory bowel disease (IBD), is attributed to dysbiosis and the mucosal immune system (Rooks et al., 2014). The disease is characterized by enrichment of flagellated bacteria resident in the microbiome and its encroachment into the inner mucus layer and the intestinal epithelial cells (IEC) (Erben et al., 2014; Okumura et al., 2016; Tran et al., 2019). LY6/PLAUR Domain Containing 8 (LYPD8) (Okada et al., 2020)) is a secreted protein that binds polymerized flagella and prevents bacterial motility on the epithelial surface (Hsu et al., 2017)). Notably, flagellin, ound at high levels during intestinal inflammation (Lodes et al., 2004), is recognized by Toll-like Receptor 5 (TLR5) on IEC. An adaptive immune response is thought to ensue to maintain immune homeostasis (Chassaing et al., 2014a; Chassaing et al., 2014b; Cullender et al., 2013). However, despite these cues, during intestinal health and disease, the functional importance and consequence of bacterial motility in a microbial consortium is unknown.

Swimming and swarming are the two primary and common forms of bacterial motility (Kearns, 2010). Swarming, driven by flagella, is a fundamental process in certain groups of bacteria characterized by collective and rapid movement across a surface (Be’er and Ariel, 2019; Kearns, 2010). This process, in contrast with swimming, offers bacteria a competitive advantage in occupying specific niches (e.g., seeding colonization) (Barak et al., 2009); however, the cost-benefits to bacteria (Butler et al., 2010; Finkelshtein et al., 2015) and consequences to its host or the environment remain primarily unknown (Allison et al., 1994).

In the context of intestinal inflammation, we show that bacterial swarmers are a feature of a stressed intestine in mammals. In a mouse model of intestinal stress, bacterial swarmers, when dosed in sufficient abundance, suppressed intestinal inflammation compared to its swarming deficient mutants. We focused on a novel *Enterobacter* swarming strain SM3, which did not show protection in germ-free mice’s acute colitis model. However, we maintained its activity in a TLR5 knockout IL-10R antibody-induced mouse model of colitis harboring conventional microflora. SM3 supplementation in mice with acute intestinal inflammation exhibited enrichment of beneficial anaerobes, specifically the S24-7 group of bacteria, in the feces. SM3 also promoted a bacterial strain belonging to the family S24-7, only when in close interaction, in an *in vitro* co-culture assay. Swarming deficient strains of SM3 had reduced S24-7 levels in feces. We posit that it is likely the act of swarming *in vivo*, which facilitates close interaction between SM3 and S24-7, enriches the latter taxa. S24-7 is strongly associated with intestinal healing in mice (Borton et al., 2017; Osaka et al., 2017).

## Results

### Presence of Bacterial Swarmers is a feature of a stressed intestine

To test whether bacterial swarmers are associated with human and rodent gut health, we developed a modified swarming assay using feces based on an established soft-agar plate assay utilized for single species (Morales-Soto et al., 2015). Since prototypical swarming bacteria (e.g., *Proteus mirabilis*, *Pseudomonas aeruginosa*) are associated with virulence (Allison et al., 1994; Overhage et al., 2008), we surmised that bacterial swarming might be well represented in colonoscopy samples and feces from humans with bacterial virulence-associated pathologies (e.g., intestinal inflammation)(Yang and Jobin, 2014)). We obtained colonoscopy aspirates from individuals with a progressive illness (inflammatory bowel disease - Crohn’s and ulcerative colitis and other common forms of intestinal stress like intestinal polyps (Crespo-Sanjuán et al., 2015; Jass, 2003) as well as age and gender-matched controls (those without a clinically active illness). Within our sampling pool, bacterial collective spreading on soft agar was over-represented in cases with overt or clinically active intestinal stress (Fig. 1a-b). As a preliminary assessment, we judged bacterial swarmers’ presence in feces by the bacterial spread with a surfactant layer on soft-agar followed by isolation, identification by MALDI-TOF, and validation of its swarming motility (STAR Method Table 1).

**Figure 1.**
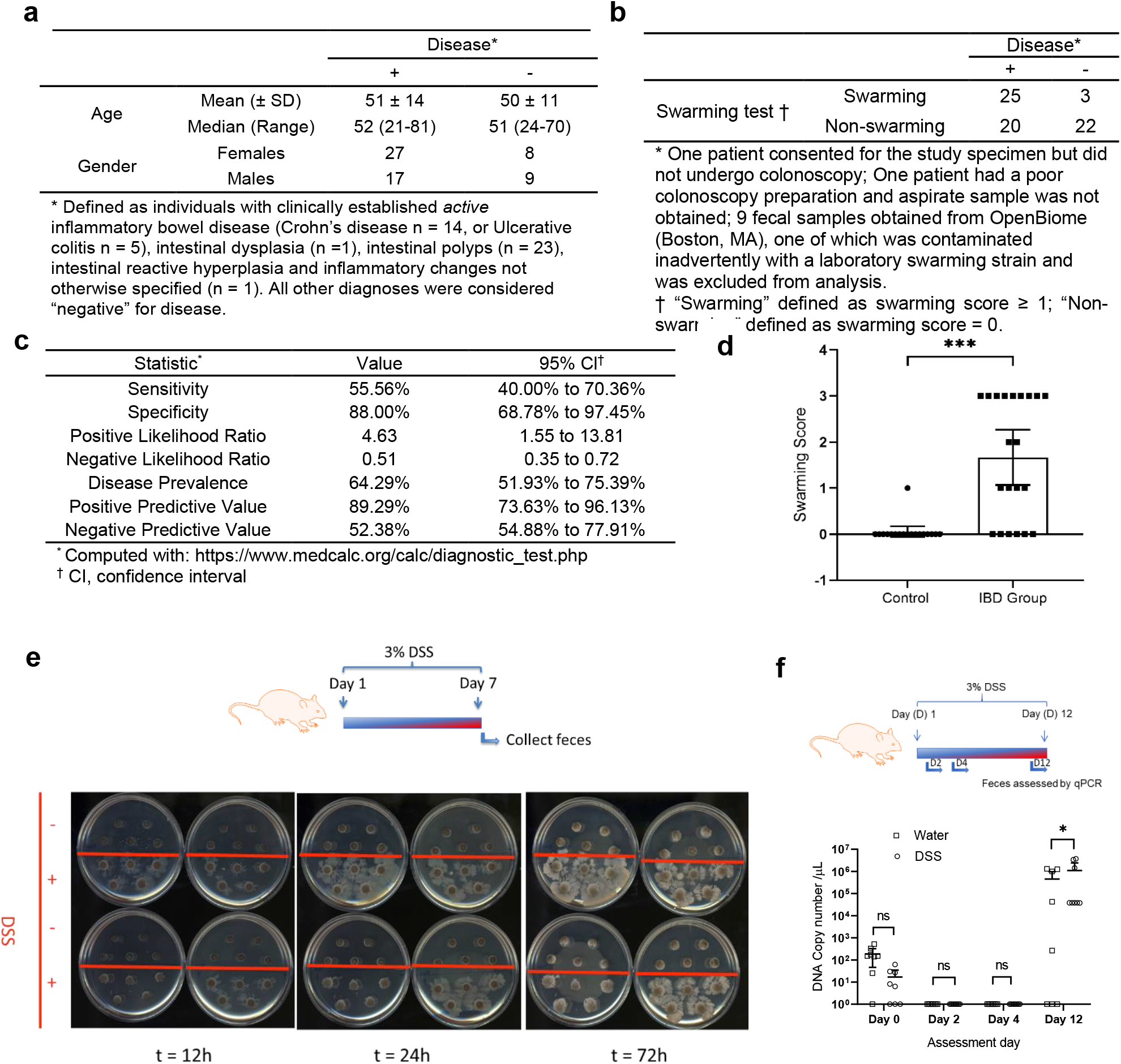
Effect of intestinal inflammation on bacterial swarming. **a-c**, Human colonoscopy aspirates (n = 45 intestinal disease; n = 25 non-disease) were spotted on 0.5% agar plates and the swarming assay performed. **a,** Colonoscopic washes were obtained from individuals with active intestinal disease and matched controls. Swarming assays performed using aspirates were binned by disease as defined both clinically and by intestinal histopathology, where available. **b**, Clinical demographics are described for the disease and non-disease population. **c**, Swarming assays’ clinical test characteristics. **d**, Swarming assays (72h) of fecal samples collected from pigs with and without IBD. Swarming scores - 0: no swarming, 1: swarming within 72h, 2: swarming within 48h, 3: swarming within 24h or less (Control: n = 6; IBD: n = 7, each in triplicate, sampled from distinct regions of the semi-solid feces). **e**, C57BL/6 mice (8-week old) were exposed to water or DSS water for 7 days (n = 4 per group). Fecal samples of control group (above red line) and DSS group (below red line) were collected for swarming assay. Swarming plates were scanned at 12, 24 and 48 hours. **f**, In a separate experiment, C57BL/6 mice (8-week old) were exposed with water or DSS water for 12 days (n = 8 per group). Fecal samples were collected for DNA extraction and SM1/SM3-specific PCR analysis was performed, and DNA copy number ascertained. Unless otherwise noted, data represented as mean and 95% CI, significance tested using Fisher’s Exact test.

In this pilot evaluation, the specificity and positive predictive value of the test for disease as defined was approximately 88 and 89%, respectively. In comparison, the test’s sensitivity and the negative predictive value was only approximately 56 and 52%, respectively (Fig. 1c). Similarly, feces collected from pigs with active inflammatory bowel disease also showed an increased prevalence of collective spreading and swarming compared to control pigs (Fig. 1d). Despite the caveat that our approach might preclude the selection of swarmers that do not produce surfactant (Kearns, 2010), these pilot data indicate that collective spreading and swarming is a specific feature, and potentially a biomarker of an intestinal pathology as defined by harboring active intestinal inflammation or polyps.

### Novel *Enterobacter* swarming strains were isolated from mouse feces

To identify the relevance of swarmers on host health, we focused on isolating endogenous swarming bacteria residing in rodents and humans. An initial approach was to determine if a single dominant swarming species could always be isolated from a polymicrobial culture (such as mammalian feces). In a competitive swarming assay, a mix of different pure bacterial cultures gave rise to a single bacterial species populating the leading edge of the swarm colony on agar (Fig. S1a-b). In congruence with our observation, a recent study has shown species dependence on motility during niche dominance, and stable coexistence when present in low abundance in a mixed population (Gude et al., 2020). Similarly, swarming assays using the pooled mouse or individual human feces yielded single species of a dominant swarmer as identified by MALDI-TOF (STAR Method Table 1; Fig. S1b). To test whether swarming bacteria are also present in preclinical models, we screened feces of mice exposed to dextran sulfate sodium (DSS), a chemical colitogen causing acute colonic inflammation (Chassaing et al., 2014a; Perse and Cerar, 2012). In a single experiment, we found three identical isolates from two different mouse fecal specimens - Strain 1 from mice exposed to water and Strain 2 and 3 from mice exposed to dextran sulfate sodium (DSS), respectively (Fig. S2a). Swarmers (in feces) were uniformly absent in water exposed mice, while present in DSS exposed mice (Fig. 1e). We picked the edge of the swarm colonies (as marked on Fig. S2a), then serially passaged twice on 1% agar from a single colony, and subsequently re-tested for swarm behavior on 0.5% agar plates (Fig. S2b). Strain 3 swarmed significantly faster compared to Strain 1 and 2. Interestingly, 16S rRNA gene analysis and Multi Locus Sequence Typing (Fig. S2c) identified the isolated strains to be closest to *Enterobacter asburiae*. Whole-genome sequence comparison of these *Enterobacter* strains (Fig. S2d) with related taxa *Enterobacter asburiae* and *Enterobacter cloacae*, revealed that all the three strains isolated here were “nearly identical” [>99% identical, one contig of 5,107,194 bp (NCBI BioProject PRJNA558971)] and phylogenetically distinct from the reference strains. Taken together, using an agar-based assay to isolate dominant swarmers from a heterogenous culture, we were able to isolate nearly identical strains with striking differences in their swarming potential. Strain 1 (*Enterobacter* sp. SM1) originated from feces of the vehicle (water) treated mice, while strain 2 (*Enterobacter* sp. SM2) and strain 3 (*Enterobacter* sp. SM3) originated from feces of DSS-induced colitis mice. Average Nucleotide Identity (ANI) analysis using OrthoANI (Yoon et al., 2017) found ~96% identity ANI between SM1/SM3 and *E. asburiae*. Interestingly, a quantitative PCR sequencing-based approach to accurately identify SM1 or SM3 like bacteria in feces showed an increase in its abundance during the evolution of DSS-induced colitis. The proportion of mice with high copy number values (>10,000 DNA copy number /μL) was significantly higher in the DSS group than water only group (Fig. 1f).

### Swarming *Enterobacter sp.* SM3 abrogates intestinal inflammation in a mouse model of colitis

To determine the functional consequence of bacterial swarmers in the host, we administered the “near-identical” swarming competent SM1 or SM3 strains to mice with DSS-induced colitis. In comparison with SM1, SM3 is a hyperswarmer (Fig. S3a; Video S1), but both strains possess the same swim speed (Fig. S3b, c), surfactant production (Fig. S3d) or growth rate (Fig. S3e). In contrast to that observed with SM1, SM3 significantly protected mice from intestinal inflammation (Fig. 2a-f). Comparison of clinical parameters showed that SM3 significantly protected from body weight loss (Fig. 2a), increased colon length (Fig. 2b), reduced the colonic inflammation score (Fig. 2d), and had reduced expression of pro-inflammatory mediators compared to vehicle-treated colitic mice (Fig. 2e-f). To test the mucosal healing capacity of swarming bacteria, we administered strains SM1 and SM3 to mice during the recovery phase of DSS exposure (Suzuki et al., 2016). When compared to the vehicle, SM3 significantly improved weight gain and colon length with reduced total inflammation and fibrosis at the microscopic level (Fig. 3). To identify if the effect is flagella mediated, we used a TLR5KO IL-10R neutralization-induced colitis model of mice. SM3 also significantly protected from body weight loss (Fig. 4a), reduced spleen and colon weight (Fig. 4b-c), increased the cecum weight (Fig. 4d), reduced serum KC level and lipocalin level (Fig. 4e-f), reduced levels of fecal lipocalin (Fig. 4g), reduced myeloperoxidase activity (Fig. 4h), and had reduced the colonic inflammation score (Fig. 4i), when compared to the SM1. We did not find differential regulation of any virulence associated genes between SM1 and SM3 strains when collected from swarming plates (Fig. S3f-g). SM3 and its isogenic transposon mutants (SM3_18 and SM3_24) only differed in swarming potential (Fig. S3h), but not swimming speed (Fig. S3i, j), surfactant production (Fig. S3k), or growth rate (Fig. S3l). In mice exposed to DSS, SM3, but not the swarming deficient mutants (SM3_18 and SM3_24), showed significant protection against weight loss (Fig. 2g), colon length (Fig. 2h), and inflammation (Fig. 2i). During the course of experiment, on day 4, the levels of SM1, SM3 and its mutants present in feces were not significantly different (Fig. S4a-b). We chose to enumerate bacterial levels in feces on day 4 due to the equivalent pathological conditions of mice, as defined by weight change, when treated with different strains. To identify, if the loss of protection by SM3_18 could be related to slightly higher levels of its presence compared to SM3, although not significant, we performed a dose attenuation study. Even at low dose, the levels of inflammation as represented by lipocalin concentration and the weight loss was not significantly different from the vehicle group, negating the possibility of any associated virulence that may attribute to loss of protection by SM3_18 (Fig. S4e-f). Furthermore, we did not find significant difference of any virulence associated gene between SM3, and SM3_18, and SM3_24 strains when collected from swarming plates. Thus, in our mutants, pleiotropic effects of gene mutations on virulence is not a cause for lack of protection from inflammation. Together, however, these data indicated that SM3 with swarming properties, as opposed to swarming-deficient strains, is associated with anti-inflammatory activity. In accordance with these results, a diverse set of commensal swarmers (*Bacillus subtilis* 3610 and *Serratia marcescens* Db10) and a clinical strain of *S. marcescens* (isolated from the surface washing of a human dysplastic polyp) exhibited protection against DSS induced inflammation in mice (Supplementary Text and Fig. S5 & S6).

**Figure 2.**
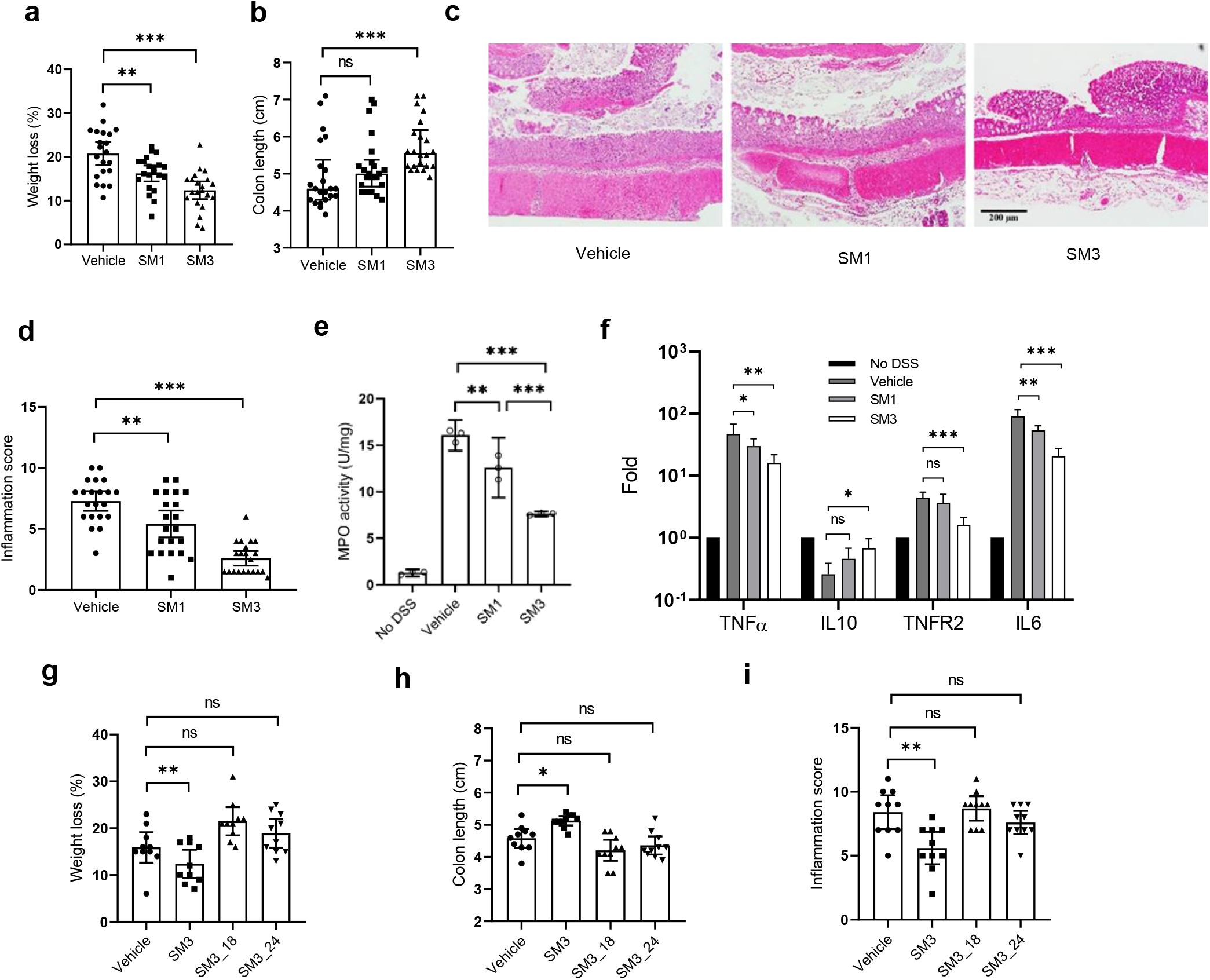
Effects of *Enterobacter* sp. SM strains on DSS induced colitis in C57BL/6 mice. **a-f**, 8-week old mice were exposed to DSS water and treated with vehicle (LB), SM1 or SM3 by oral gavage for 10 days. **a-b** indicates weight loss (**a**) and colon length (**b**) (n = 21 per treatment group). **c**, Representative images (100x magnification) of H&E stained colonic section treated with vehicle (left), SM1 (middle) and SM3 (Borton et al.). **d**, Inflammation score (n = 21 per treatment group). **e-f**, In a separate experiment, myeloperoxidase (MPO) enzyme activity was determined (n= 3, each in duplicate) (**e**). Colon total RNA (n = 4) was isolated and reverse transcribed to cDNA. RT-qPCR data show fold induction of mRNA (TNFα, IL10, TNFR2, IL6). PCR was repeated in quadruplicate. The expression was normalized to internal control, TBP. The entire experiment was repeated n = 2 for reproducibility (**f**). **g-i**, In a separate experiment, C57BL/6 mice (8-week old) were exposed to DSS water and administered vehicle (LB), SM3, or its mutants (SM3_18 or SM3_24) for 10 days. **g-i** indicates weight loss (**g**), colon length (**h**) and inflammation score (**i**) (n = 10 per treatment group). Unless otherwise noted, data are represented as mean and 95% CI, and significance tested using one-way ANOVA followed by Tukey’s post hoc test. **c**, data represented as median and interquartile range, and significance tested using Kruskal-Wallis followed by Dunn’s multiple comparisons test. * *P* < 0.05; ** *P* < 0.01; *** *P* < 0.001; ns, not significant. H&E, Hematoxylin and Eosin; TBP, TATA-Box Binding Protein; CI, Confidence Interval.

**Figure 3.**
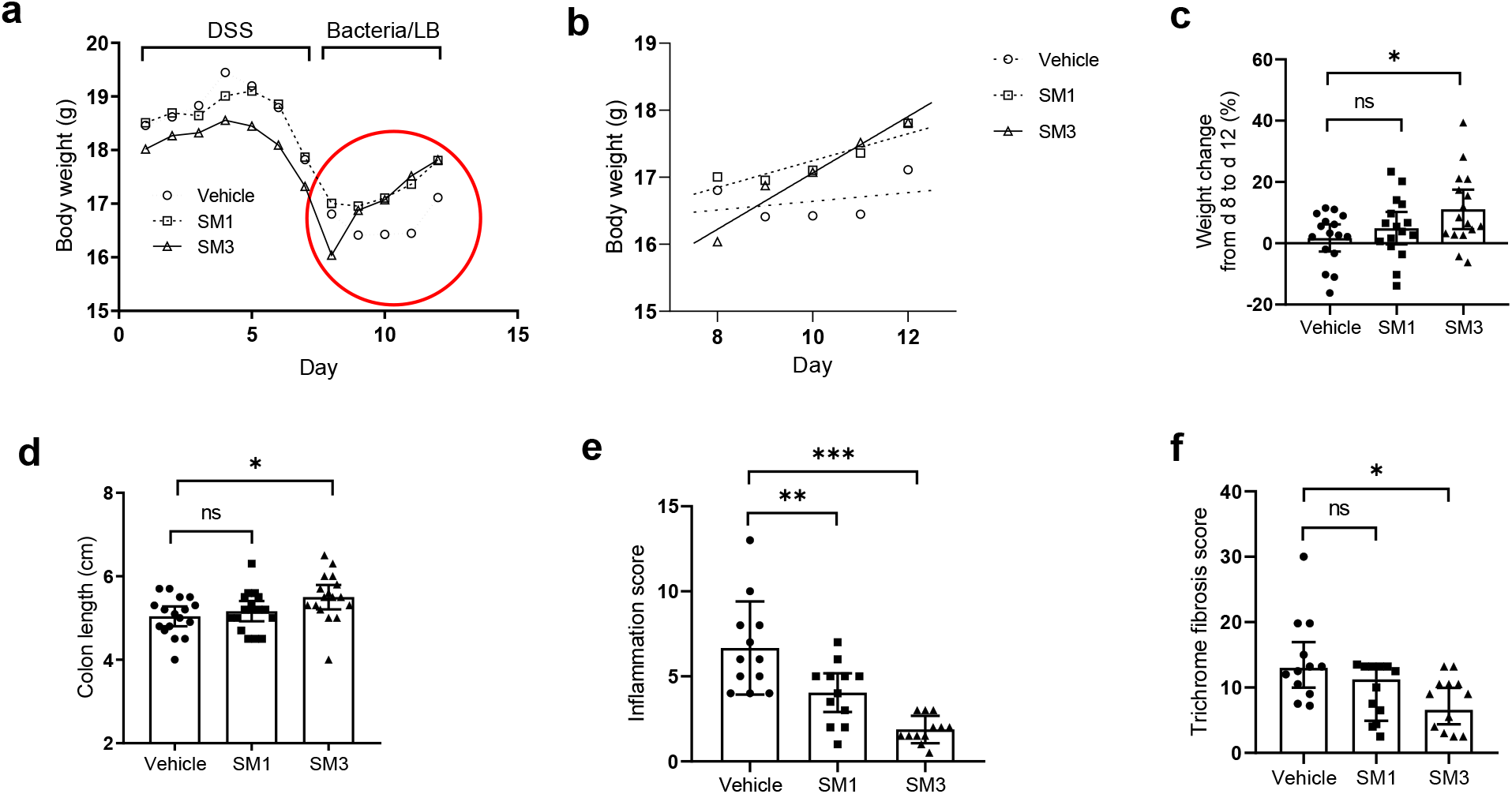
Effect of *Enterobacter* sp. SM1 or SM3 strain on DSS induced colitis in C57BL/6 mice during recovery phase. 8-week old mice were exposed to DSS water for 7 days. On day 8, DSS water was replaced with drinking water and mice were administered vehicle (LB), SM3 or SM1 for 5 days. **a-c**, indicates the weight change. **a**, Day by day weight change. **b**, Day by day weight change from day 8 to day 12 (healing phase, red circle in **a**) was separately plotted, and the best fitting line was added to each group using linear regression. The slopes for the regression lines are 0.421 (SM3), 0.201 (SM1) and 0.065 (Vehicle). The slope of SM3 group is significantly deviated from zero (P = 0.013), while the SM1 and vehicle group are not (P = 0.240, 0.754 respectively). **c**, Percent weight change from day 8 to day 12. **d-f**, indicates colon length (**d**), inflammation score (**e**), and trichrome fibrosis score **(f)** (n = 16 per treatment group except for **e** and **f**, four colon specimens per group were used for other experiments). Unless otherwise noted, data represented as mean and 95% CI, and significance tested using one-way ANOVA followed by Tukey’s post hoc test. **f**, data represented as median and interquartile range, and significance tested using Kruskal-Wallis followed by Dunn’s multiple comparisons test.

**Figure 4.**
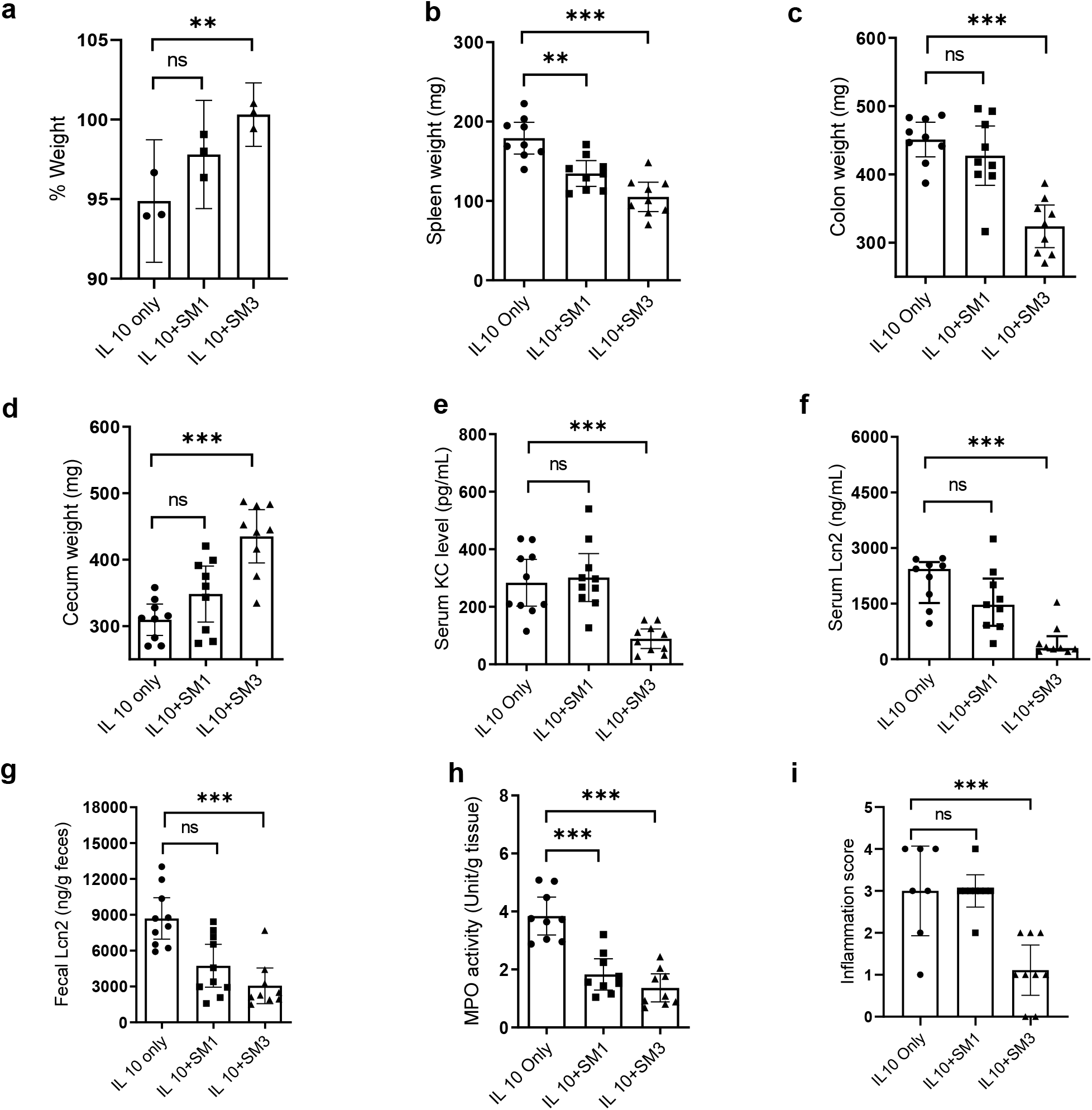
Effects of *Enterobacter* sp. SM strains on IL-10R neutralization-induced colitis in *Tlr5* KO mice. 8-week-old *Tlr5*KO mice were administered rat anti-IL-10R monoclonal antibody (1.0 mg/mouse, i.p.) (BioXcell) at day 0 and 7. SM1 or SM3 was gavaged every 3^rd^ day from day 1 until day 18. **a**, Weight percentage compared with day 0 (n = 3 per treatment group). **b**, Spleen weight (n = 9 per treatment group). **c**, Colon weight (n = 9 per treatment group). **d**, Cecum Weight (n = 9 per treatment group). **e**, Serum KC level (n = 10 per treatment group). **f**, Serum lipocalin (n = 9 per treatment group). **g**, Fecal lipocalin (n = 10 for each group, 1 fecal sample in SM3 group was used for other study). **h**, Myeloperoxidase (MPO) (n = 9 per treatment group). **i**, Inflammation score (n = 9 for each group, 2 tissue samples in IL 10 only group were used for other study). Unless otherwise noted, data are represented as mean and 95% CI, and significance tested using one-way ANOVA followed by Tukey’s post hoc test. **f-g**, data represented as median and interquartile range, and significance tested using Kruskal-Wallis followed by Dunn’s multiple comparisons test.

### SM3 mediated abrogation of intestinal stress is microbiome dependent

To determine if the anti-inflammatory role of SM3 is dependent on the conventional intestinal microbiome composition, germ-free mice transferred to specific pathogen-free conditions (GF/SPF) and exposed to DSS-induced colitis, were treated with SM3. This strain was unable to abrogate intestinal inflammation in GF/SPF mice (Fig. 5a). We analyzed fecal samples of colitic mice (conventional and GF/SPF) with SM3 administered using 16S rRNA gene profiling. In contrast to GF/SPF mice, conventional mice feces showed specific enrichment of anaerobes belonging to the family S24-7 and Lactobacillaceae within SM3 treated mice when compared to vehicle mice (Fig. 5b). Specifically, in conventional mice, we found a significant increase in the abundance of S24-7 with SM3 gavage compared to vehicle in DSS exposed mice (Fig. 5c). However, quantitative PCR analysis of the levels of S24-7 in the feces of DSS-induced colitis mice gavaged with SM1 or SM3_18 or SM3_24, that did not exhibit protection from intestinal inflammation, was significantly reduced (Fig. 6a). In mice not exposed to DSS, the levels of S24-7 bacteria remain stable in SM3 treated group when compared with the untreated group (Fig. 5c). Within DSS exposed conventional mice, we observed that enriched S24-7 negatively co-occurred with pathogenic taxa such as the Peptostreptococcaceae and Enterobacteriaceae (Fig. 5d). Together, these data suggest that protection from intestinal inflammation by SM3 is associated with the presence of beneficial S24-7 group of bacteria (Volk et al., 2019).

**Figure 5.**
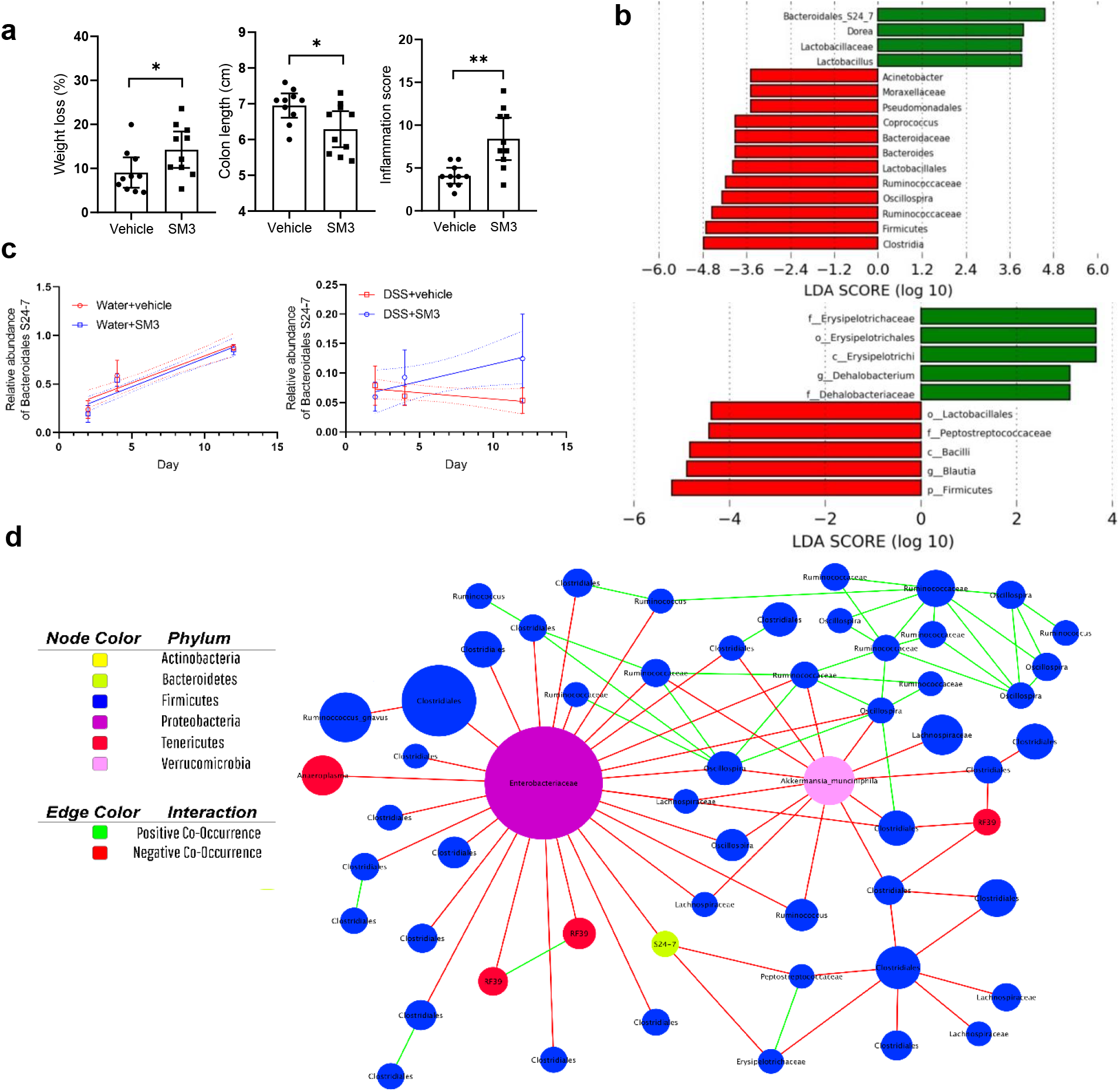
Effects of SM3 on the intestinal microbiota of GF/SPF and conventional mice. **a**, C57BL/6 GF/SPF mice (5-week old) were exposed to DSS water and treated with vehicle (LB) or SM3 for 6 days. **a** indicates weight loss (left), colon length (middle), and inflammation score (Borton et al.) (n = 10 per treatment group). **b**, Linear discriminant analysis (Walter et al.) Effect Size (LEfSe) plot of taxonomic biomarkers identified using feces of SM3 treated conventional (n = 10) (upper) and GF/SPF (n = 10) (lower) colitic mice on day 12 and day 6, respectively, as compared to vehicle (n = 10). Green and red bars indicate bacterial enrichment within SM3 treated and vehicle group respectively. All taxa that yielded an LDA score >3.0 are presented. **c**, Relative abundance of S24-7 in the feces from DSS (Allison et al., 1994; Borton et al.) and control (left) mice treated with SM3 or vehicle (n = 8 per treatment group). Linear regression line was fit to show the trend of the change (dotted lines show the 95% confidence bands). The slope of the SM3 treated group is similar to vehicle in water control group (*P* = 0.7827), but significantly different in DSS group (*P* = 0.0182). **d**, Co-occurrence network plot showing strong positive and negative correlations between OTU abundances. All networks were generated with CoNet and visualized in Cytoscape. Processing was applied to the dataset with CoNet. Input filtering constrained the minimum occurrence of OTUs and considered only those present in at least 50% of samples. Standardization normalized dataset columns. Networks were constructed using Spearman’s correlation methods with threshold setting at 0.9, Bray Curtis dissimilarity at the automatic threshold setting, and Kullback-Leibler dissimilarity at the automatic threshold setting; the edge selection parameter was set to 30 for the strongest positive and negative correlations. Randomization steps included permutations and bootstraps with filtering of unstable edges and Benjamini-Hochberg procedure with a P-value of 0.05. Node clusters with less than or equal to three edges were not shown in the final network. Edge coloration indicates copresence in green or mutual exclusion in red. Nodes were colored by taxonomic phylum and labeled by the highest taxonomic ranking available.Unless otherwise noted, data are represented as mean and 95% CI, and significance tested using a two-tailed Student’s t-test. OTU, Operational Taxonomic Unit; GF/SPF, Germ-Free mice transferred to specific pathogen free conditions.

**Figure 6.**
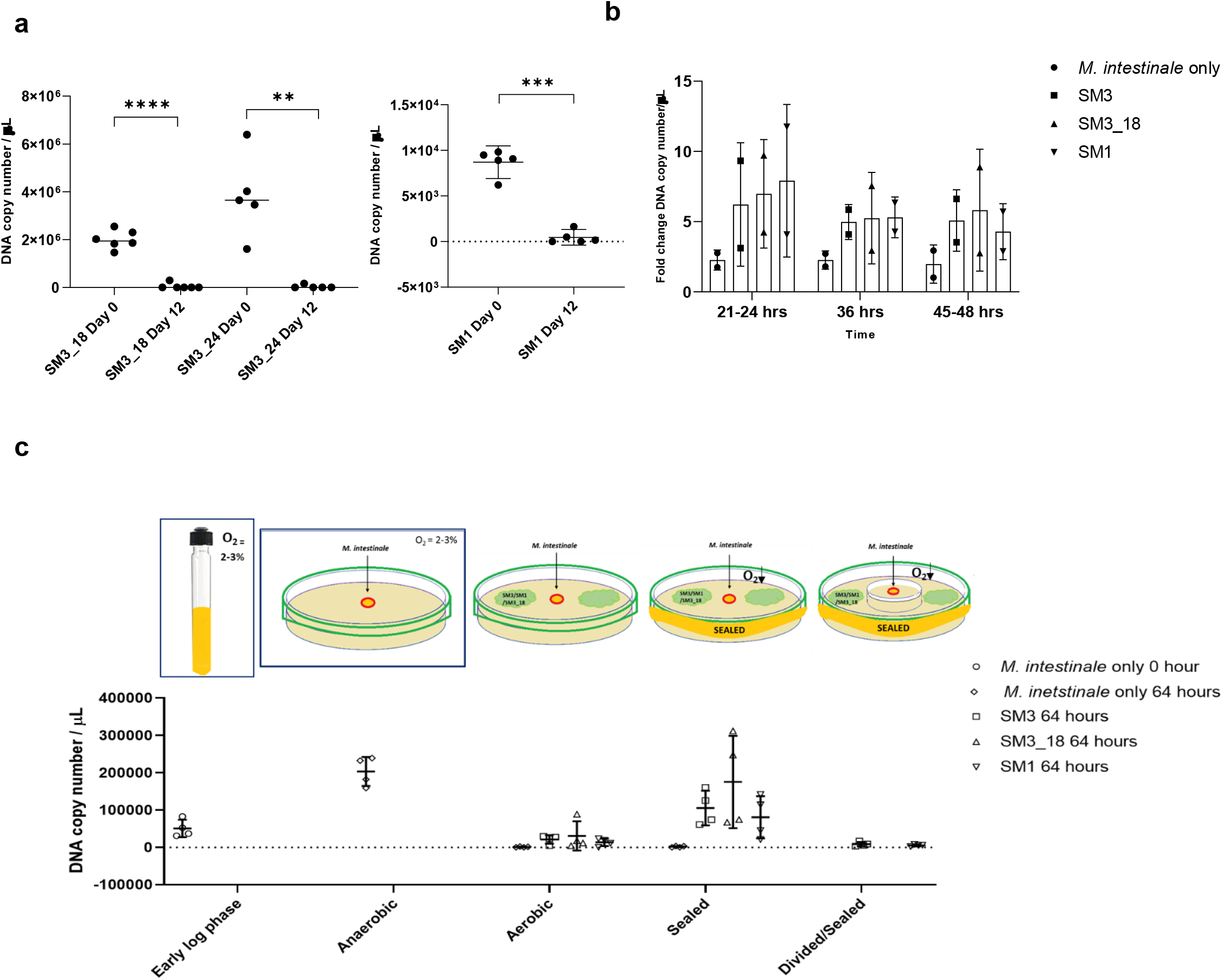
Effect on S24-7 levels in the presence of SM3 and the insufficient (or inefficient) swarming variants *in vivo* and *in vitro*. **a, Eight** (8)-week old mice (n ≥ 5 per treatment group) were exposed to DSS water and treated with SM3_18, SM3_24, and SM1 by oral gavage for 12 days. Total DNA was extracted from feces collected on day 0 and day 12, processed and assessed using qPCR. Five (5) ng of total DNA in conjunction with S24-7 specific primers were used to quantify bacterial copy numbers. In each assay, DNA copy number/μL was calculated based on an internal standard curve. **b-c**, *In vitro* co-culture assay using *M. intestinale* cells grown in Chopped meat medium under anaerobic condition until early log phase (OD_600_ ≈ 0.5) were used. **b**, Fold change DNA copy number/μL relative to *M. intestinale* monoculture. In broth-based assay, 2μL of early log phase cells of SM3, SM3_18, or SM1 was added to *M. intestinale* cells and mixed cells or monoculture of *M. intestinale* was collected at regular intervals (21-24, 36, 45-48 hrs). **c**, In swarming-plate based assay, early log phase *M. intestinale* was transferred in the bore-well and SM3, SM3_18, or SM1 was allowed to swarm either under aerobic or sealed condition at 37°C and RH ≈ 50%. Plates were sealed using parafilm to create and maintain anaerobiosis due to the act of swarming. *M. intestinale* grown under anaerobic condition was used as a positive control. In Divided/Sealed condition, swarming region was physically separated from the bore-well containing *M. intestinale* and sealed using parafilm. Closed boxes represent incubation in an anaerobic chamber. DNA extracted from equal volume of culture and resuspended in equal volume of TE buffer was used for qPCR in conjunction with *M. intestinale* specific primers. **a**, Data represented as mean and 95% CI, and significance tested using paired t-test. **b-c**, Data represented as mean (+ SD)(n = 2 independent experiments and 2 technical replicates for each).

### *Enterobacter sp.* SM3 promotes growth of *Muribaculum intestinale in vitro*

A recent study has reported the first cultured bacterium *Muribaculum intestinale* (DSM 28989) that belongs to Bacteriodales S24-7 family (Lagkouvardos et al., 2019). We used this strain to delineate any potential interspecies interaction with SM3 using an *in vitro* co-culture assay system. However, in precedence, we assessed if the strain *M. intestinale* shared sequence homology to any of the S24-7 taxa identified in our fecal 16S rDNA profile. OTU_5, which was found in highest abundance among all other OTU’s representing S24-7 taxa, exhibited >96% identity to *M. intestinale* (Fig. S7). Hence, we performed a broth-based co-culture assay using this strain and SM3 or SM1 or SM3_18. Interestingly, the proportion of *M. intestinale* during co-culture was higher compared to its monoculture at any tested time point. SM3 as well as the partially swarming deficient strains, SM1 and SM3_18, had a two-four (2-4) fold increase in DNA copy number/μL, when analyzed by qPCR using S24-7 specific primers (Fig. 6b). We also designed and developed a plate-based co-culture assay to compare the effects of swarming bacteria SM3 and swarming-deficient variants, SM1 or SM3_18, on the growth of *M. intestinale*. In this assay, swarming plates harbored a central bore well containing *M. intestinale* that guarantees a direct or indirect interaction with the spreading bacteria on agar of the same plate. The plates were sealed so that the act of swarming generated an anaerobic environment suitable for the growth of *M. intestinale*. At 64 hours, in congruence with the broth co-culture assay results, we observed an increase in *M. intestinale* counts with SM3, SM3_18, and SM1 (Fig. 6c). To better understand if the observed increase in *M. intestinale* levels is mediated by a direct or an indirect interaction during the co-culture studies, and not solely due to the reduced oxygen concentration in the agar plate, we developed a separate plate-based co-culture assay. In this assay, the swarming region was physically separated from the central bore well containing *M. intestinale* to prevent any direct or indirect interaction with the swarming bacterium. In this system, as the bacteria swarmed on the agar surface over 64h, oxygen levels were reduced. *M. intestinale* showed no growth under the conditions tested (Fig. 6c, Divided/Sealed). Overall, our results suggest, that both planktonic and swarming cells of SM3, SM1, or SM3_18, when co-cultured *in vitro*, can promote the growth of S24-7 family (*M. intestinale)*, independent of reduced oxygen concentrations in the environment. Coincidently, the development of significantly reduced oxygen concentrations in the environment are also observed *in vivo* but only with SM3 and not SM1 or the other SM3 mutant bacteria (Supplementary text and Fig. S8). Our results suggest that SM3 proximity to *M. intestinale* is necessary for the induction of the growth of the latter species.

### The act of bacterial swarming is likely a phenomenon *in vivo*

The intestinal mucosa is relatively uneven during inflammation due to the loss of mucin (Fig. S10)(Sasaki et al., 2008). We conjectured, therefore, that swarmers might have an added advantage in niche dominance on inflamed tissue. Indeed, a race assay designed on mouse mucosal surface demonstrated the biological relevance of swarming. On mouse mucosa, it is not possible to visualize bacterial cells directly, so we developed the mucosa racing experiment to determine the dominant motility type. We developed a hyperswarmer strain of SM1, HS2B, that swims slower but swarms at a similar rate and extent to that of SM3, and an isogenic mutant Δ*flhE* SM1 that is swarm deficient but swim competent when compared to SM1. In this experiment, identically sized segments of “normal or control” mouse colon was placed on a hybrid agar plate (1%/0.3%) (see STAR Method) and equal concentration (CFU/mL) of bacteria spotted. As a negative control, sterile LB and a non-motile strain of Δ*mot*A SM1 when spotted did not show any motility across the mucosal surface even after 20h (Video S4). In comparing the “motility rates” across the normal colon mucosa (see STAR Method), the hyperswarming HS2B demonstrated slow motility when compared to Δ*flhE* SM1 (Fig. S11c). However, when the same experiment was performed using inflamed mouse colon after exposure to DSS *in vivo*, the motility of HS2B was significantly faster than swarming deficient Δ*flhE* SM1 (Fig. S11d). We also used *S. marcescens* strains, swarming Db10 and non-swarming JESM267 (*swrA*) mutant to confirm our hypothesis. While on normal colon mucosa Db10 and JESM267 did not significantly differ in the motility rates, however, Db10 was significantly faster than JESM267 on colitic mucosa (Fig. S11g-h). This indicates that swarming bacteria finds an advantage in motility on a colitic mucosa compared to normal mucosa (Video S2-4). Parenthetically, we have recently developed and used a distinct Polydimethylsiloxane (PDMS) based confinement tool to show that a unique motility pattern associated with bacterial swarming is also seen on tissue from colitic mice gavaged with SM3 (doi: https://doi.org/10.1101/2020.08.30.274316). Furthermore, to understand if SM3 does swarm *in vivo*, we searched for the presence of transcriptomic markers in the feces that can be linked to SM3 swarming physiology. In agar-based studies of RNA sequencing of SM3 obtained from the edge of a swarming colony versus the pre-swarming colony at center, a singular pathway was significantly upregulated – the lipid A biosynthetic pathway (fold change 3 fold, q value = 0.0376). Meta-transcriptomic analysis of feces from SM3 treated DSS induced colitic mice identified steady increase of lauroyl acyltransferase transcript involved in Lipid A biosynthesis on Day 4 and Day 12 when compared to Day 0 (Fig. S12a). However, heat killed SM3 treatment showed reduction in transcript abundance by Day 12. Other genes known to be associated with swarming such as the sigma factor FliA and nitrate reductase NarH were also enriched in SM3 versus heat killed SM3 gavaged colitic mice (Fig. S12b). Normal mice gavaged with SM3 or heat killed SM3 did not show enrichment of these genes. Collectively, our data provides multiple lines of evidence suggesting that bacterial swarming is a likely phenomenon *in vivo*, and a motility form that is necessary for the induction of *M. intestinale* growth.

## Discussion

Our study finds that intestinal inflammation itself promotes a protective niche that allow enrichment of bacterial swarmers. The inflammatory milieu likely provides a permissive environment for stress adaptation and swarming behavior. Surprisingly, however, these bacterial swarmers when dosed in sufficient abundance abrogate intestinal inflammation in mice. We focused on a novel bacterium, *Enterobacter sp.* SM3, which is resident to the intestinal microflora of mice. *In vivo*, SM3, but not SM1, or SM3 swarming deficient mutants (poor swarmers), influenced the specific enrichment of S24-7 group of bacteria. Notably, the family of S24-7 (*Muribaculaceae*) are known to repair barrier function in inflamed mice intestines (Osaka et al., 2017; Volk et al., 2019). However, the *in vitro* co-culture experiment proved that a close interaction between SM3 and S24-7 group of bacteria is essential for its enrichment. Thus, we hypothesized that it is the relative hyperswarming activity of SM3 (but not very slow swarming SM1 or SM3 mutants) that may facilitate a close interaction with S24-7 group of bacteria, *in vivo*. Further support of this hypothesis comes from the ability of bacteria to swarm on a mucosal surface afflicted only by colitis (ex vivo mucosal race assay) and the meta-transcriptomic analysis demonstrating that at least two major metabolic pathways known to be associated with motility are upregulated in coltic mice administered SM3 (but not its heat-killed counterpart). The present mechanism implicates swarming SM3 to directly enhance S24-7 (*Muribaculaceae*) which then suppresses host inflammation (Graphical Abstract). Nevertheless, we do not exclude other direct or indirect effects of the swarming SM3 on mucosal inflammation and healing. However, if present, it would assist in suppressing host inflammation in conjunction with enrichment of S24-7 group of bacteria in the gut.

Increased prevalence of swarmers from stressed intestinal contents of mammals (humans, pigs, and rodents) encouraged us to investigate its relevance on host health during colitis. We developed a method to isolate dominant swarmers resident in the host from fecal and colonoscopic washing samples. Screening of several bacterial species allowed us to identify genetically identical gram-negative bacterial strains SM1 and SM3 from feces of normal and DSS induced colitic mice, respectively. Bacterial size (Be’er et al., 2013), swimming speed (Sokolov and Aranson, 2009), cell density (Zhang et al., 2010), and several other factors (Angelini et al., 2009) are known to affect swarming and swarm patterns *in vitro*. Comparing these factors that could affect the swarming ability of SM1 and SM3, we found that the swimming speeds and growth rates were similar (Fig. S3a-e). Swarming bacteria secrete surfactants, such as surfactin, that facilitate during motility on a solid surface (Kearns, 2010). As surfactin is reported to attenuate TNBS induced colitis, possibly by differentially regulating anti-inflammatory and pro-inflammatory cytokines (Selvam et al., 2009), we determined surfactant levels using blood agar assay, drop-collapse assay (Bodour and Miller-Maier, 1998) and drop-counting assay based on modified Stalagmometric Method (Dilmohamud et al., 2005). None of the strains showed significant difference in surfactant production when compared to its isogenic mutants, at the conditions tested (Supplementary text, Fig. S13).

Furthermore, as DSS is a polymer of sulfated anhydroglucose we performed a separate experiment to show that SM3 does not influence DSS induced cell cytotoxicity using a cell line model of intestinal epithelial differentiation (Caco-2) (Fig. S14). Thus, it is unlikely that SM3 can either metabolize and/or inactivate DSS *per se*. To identify if the pathophysiological effect observed with bacterial swarmers was specifically due to its swarming physiology, we focused on generating isogenic mutants that were swarm deficient but proficient in all other features that may otherwise affect swarming (swimming, growth rate, surfactant production). To emphasize the importance of functional flagella in protection during intestinal inflammation, we used a motor protein abrogated strain of SM1 (Δ*mot*A SM1) in mice with DSS induced colitis. Notably, this mutant strain that had equivalent growth rate and an intact flagellum but was unable to either swim or swarm showed no protection against intestinal inflammation in a DSS mice model study (Fig. S15). As a proof of concept that swarming SM3 protects from inflammation, we used two isogenic strains SM3_18 and SM3_24 that had significantly reduced potential to swarm, *albeit* with similar swimming speeds and growth rates compared to the wildtype. Transpositions in SM3_18 and SM3_24 were found to locate within the putative structural genes encoding N6-hydroxylysine O-acetyltransferase or aerobactin synthesis protein (*iuc*B) and isocitrate/isopropylmalate dehydrogenase/ADP-ribose pyrophosphate of COG1058 family, respectively. Nevertheless, transposon integration in SM3_18 led to a polar insertion that will only disrupt the expression of downstream genes *iucC*, *iucD* and *iutA* located within the operon, hampering aerobactin synthesis only. Fundamentally, genes *iucD* and *iutA* aid in iron acquisition in bacteria during nutrient limiting condition (Koster et al., 1994). A single study has also shown the dependence of bacterial phytopathogen *Pantoea stewartii* swarming on aerobactin synthesis (Burbank et al., 2015). Similarly, in SM3_24 non-polar insertion within the gene coding for putative isocitrate/isopropylmalate dehydrogenase/ADP-ribose pyrophosphate of COG1058 family, may not affect the transcription of downstream gene that has two possible promoter sequences at the 3’end of the insertion site, as determined by BPROM (Solovyev, 2011).

TLR-5 knockout mice fail to express anti-flagellin antibodies, an immunomodulatory pathway that aids in maintaining intestinal homeostasis (Cullender et al., 2013). Even in the absence of TLR5, colitic mice harboring normal microbiota showed significant protection when treated with SM3. In contrast, in a germ-free condition, SM3 lost protection allowing us to speculate the role of intestinal microbiome in the observed effect. Indeed, oral gavage of SM3 in conventional colitic mice showed enrichment of beneficial anaerobes and microaerophiles. These anaerobes belong to the family Bacteroidales S24-7 and Lactobacillaceae. S24-7, recently classified to Muribaculaceae family, forms one of the major taxa in mouse gut (Lagkouvardos et al., 2019) and has been associated with disease remission (Borton et al., 2017; Osaka et al., 2017). Similarly, *Lactobacillus* producing lactate is known to promote the proliferation of intestinal epithelial cells (Okada et al., 2013). In view of the fact that oxygen content of the intestinal lumen increases during intestinal inflammation (a shift from anoxic to oxic)(Colgan and Taylor, 2010) (Fig. S8a), it was unexpected to find enrichment of obligate anaerobes such as Bacteriodales S24-7 in SM3 treated mice. Recent metagenomic analyses have revealed the potential of S24-7 bacteria to be “nanaerobe”, permitting growth in nanomolar concentrations of oxygen (Ormerod et al., 2016). This observation allowed us to hypothesize the possible role of SM3 in the rapid depletion of oxygen in the lumen, and in turn, favoring the growth of resident anaerobes. In congruence, we observed SM3 fed colitic mice had significantly lower oxygen concentration compared to the colitic mice treated with swarming deficient variants. We conjectured the possible role of swarming movement of SM3, if occurring *in vivo*, in reducing oxygen concentration. To link swarming activity and oxygen necessity, we showed by *in vitro* experiments that SM3 cannot swarm at low concentrations of oxygen. It was further corroborated by the increase in anaerobic taxa in the feces of GF/SPF mice treated with SM3. Nevertheless, a steady increase of S24-7 specific OTU’s in SM3 treated DSS-colitic mice pointed towards a potential mechanism underlying the observed protection. Hence, we designed a broth and plate-based co-culture assay to identify possible specific interaction between SM3 and the first cultured bacterium belonging to the S24-7 family, *M. intestinale*. Both SM3 and the less swarming variants promoted growth of *M. intestinale* in co-culture assay. However, linking this observation with decrease in the levels of S24-7 in the fecal DNA obtained from SM1, SM3_18, and SM3_24 led us to speculate the essential role of swarming by SM3 in exhibiting protection. We conjectured that in addition to an anaerobic environment generated by the act of swarming on the agar plate, all the tested strains either required a direct cell-cell contact or produced a secretome, which promoted growth of *M. intestinale*. This was further validated by a plate-based assay that allowed physical separation of swarming SM3 from *M. intestinale*, but at the same time creating an anaerobic condition in the system suitable for the growth of *M. intestinale*. Together this suggest, a close spatial interaction between SM3 and S24-7 group of bacteria allow enrichment of the latter preferably in a microenvironment. *In vivo*, SM3 may aid in re-establishing hypoxia, and consequently creating an optimal environment for the growth of S24-7 and other anaerobes. The present mechanism strongly suggests that swarming (but not swimming or slow swarming) SM3 directly enhances S24-7 (*Muribaculaceae*) by attaining proximity to it *in vivo* (Graphical Abstract).

In broadening the scope of our hypothesis, we observed that even known commensal swarming strains *Bacillus subtilis* 3610 and *Serratia marcescens* Db10 protected against DSS-induced colitis in mice when compared to its isogenic swarming deficient mutants. Similar results were also observed with a clinical strain of *S. marcescens* isolated from the colonoscopic aspirates of a human colitis patient, which also swarmed *in vitro*. Since this *Serratia* isolate was obtained from human feces and not characterized at the genome sequence level, we used wildtype and heat-killed clinical strains to study their effects on mice with DSS-induced colitis. We found that the wildtype, but not the heat-killed strain protected against colitis (Fig. S6). Such protection observed with different bacteria belonging to a distinct phylogenetic group may also preclude the possibility of any loss of protection seen in case of isogenic mutants due to other pleiotropic effects. Similarly, a pathogenic swarmer strain of *Salmonella enterica* serovar Typhimurium had reduced severity and less inflammation compared to its swarm defective mutant Δ*fli*L (Fig. S16, Supplementary text). Overall, these observations suggested that bacterial swarmers and likely the act of swarming play a major role in mitigating disease severity in a mouse model of colitis.

Bacterial swarming is a fundamental process in certain groups of bacteria characterized by collective and rapid movement across a surface (Kearns, 2010), (Be’er and Ariel, 2019) and has been reported *in vitro*. We focused on understanding if swarming is likely a phenomenon *in vivo*. As an initial approach, we developed an *ex vivo* mucosal race assay that proved motility dominance of bacterial swarmers on a biological surface. Similarly, we have developed a PDMS based tool to distinguish bacterial swimming from swarming, which exhibited the possibility of the latter movement in micro pockets on DSS treated mucosal tissue. However, technical limitations to demonstrate swarming *in vivo* in a mouse model limits generalization of our data. First, *in vivo*, intestines are continually moving through a process termed by Baylis & Starling as “peristalsis” (Spencer et al., 2016). These myogenic ripples actively move food and other particles along the intestines to the anal region for defecation. If we could image our bacteria *in vivo*, the artifact of peristalsis would cloud any interpretation of swarming movement *in vivo*. Further, poor resolution in contemporary microscopic endoscopy is a limitation. Second, in order to detect collective motion *in vivo*, we would need to image the bacteria studied. GFP labeling of SM3 knocks out its swarming ability. We have also labeled the outer membrane of SM3 using click chemistry, however, upon gavage, we posit that the signal will be lost as the bacteria replicate in the gastrointestinal tract. Both limitations, at present, preclude any ability to efficiently and functionally label SM3 *in vivo*. Due to these limitations, we have demonstrated the possibility of swarming movement *in vivo* by identifying genes that are known to be associated with swarming. One of the upregulated genes in SM3 treated colitic mice, Lauroyl acyltransferase, is an enzyme involved in lipopolysaccharide (LPS) biosynthesis, which forms an integral part of cell envelope biogenesis. LPS synthesis genes are upregulated in swarmer cells that act as a lubricant, promoting swarming (Partridge and Harshey, 2013). Arguably, LPS is an important biomarker during intestinal inflammation (Im et al., 2012). Although KEGG pathway analysis showed enrichment of LPS biosynthesis pathways in SM3 treated colitic mice compared to Heat killed SM3 treatment, protection observed may indicate the upregulated pathways in association with bacterial swarming. The nitrate reductase beta chain, NarH, involved in nitrogen dissimilation is also linked to rhamnolipid biosynthesis in *P. aeruginosa* (Van Alst et al., 2007). Rhamnolipids are essential for swarming behavior in *Pseudomonas* (Caiazza et al., 2005). A similar role for NarH could be at play in *Enterobacter sp.* SM3, as there is a significant increase in nitrate reductase beta-chain transcripts found in pre-swarming SM3 cells collected from the center of the agar plate (q = 1.27E-224). Two-component system analysis using fecal metatranscriptome also identified pathways with upregulation of specific sigma factors. FliA is an alternate sigma factor that directs transcription of Class III flagellar genes and is also essential in bacterial swarming(Young et al., 1999). The HptB-sigma factor signaling system, as identified in KEGG analysis (Fig S12b), also utilizes FliA to upregulate flagellar biogenesis essential in swarming by *P. aeruginosa*(Bhuwan et al., 2012). Overall, these data together support the notion that swarming movement likely occurs *in vivo*.

In summary, our work demonstrates the unique and unprecedented role that bacterial swarmers play in intestinal homeostasis. Our work demonstrates the potential for a new personalized “probiotic” approach stemming from the ability to isolate and bank swarming microbes during colitis flares for clinical treatment during concurrent or subsequent colitic episodes.

## Acknowledgement

We thank Steve Almo, Andrew Gewirtz, Cait Costello, Jeffrey Pessin, Matthew R. Redinbo and John March for valuable discussions. We aslo thank Brad Tricomi for developing the assay “Cytotoxicity of DSS on Caco-2 cell lines in the presence or absence of viable SM3 cells”, Ehsan Khafipour for providing pig specimens (feces) and performing clinical scoring of histopathology, Cori Bargmann at Rockfeller University for gifting us the bacterial strains Serratia marcescens Db10 and JESM267. Additional assistance was obtained from Amanda Beck DVM (Histology and Comparative Pathology Core, AECOM), Olga C. Arionadis, Thomas Ullmann and Azal Al Ani (Department of Medicine, AECOM), Winfred Edelmann and Eleni Tosti (Department of Cell Biology, AECOM). The studies presented here were supported in part by the Broad Medical Research Program at CCFA (Crohn’s & Colitis Foundation of America; Grant# 362520) (to S.M); NIH R01 CA127231; CA 161879; 1R01ES030197-01 and Department of Defense Partnering PI (W81XWH-17-1-0479; PR160167) (S.M.), Diabetes Research Center Grant (P30 DK020541); Cancer Center Grant (P30CA013330 PI: David Goldman); 1S10OD019961-01 NIH Instrument Award (PI: John Condeelis); LTQ Orbitrap Velos Mass Spectrometer System (1S10RR029398); and NIH CTSA (1 UL1 TR001073). Peer Reviewed Cancer Research Program Career Development Award from the United States Department of Defense (CA171019, PI: Libusha Kelly).

## Author Contributions

H.L., S.M. conceptualized the discovery. H.L., D.K., W.C., J.T., S.M. designed and executed the swarming assays. D.L. was the Principal Investigator of the Clinical Study and provided specimens. L.K. performed genome assembly and annotation. J.W., R.L., S.M. designed and executed all the 16S, metagenomic and strain-specific PCR assays. A.D. designed; A.D., W.C., S.M., S.G. characterized bacterial mutants. B.S.Y. and M.V-K. performed tlr5KO mice study and repeat swarming assays for reproducibility. A.D., H.L., W.C. and S.M. wrote and edited the paper. S.C. and W.C. performed statistical analyses. X.L. assisted H.L. in mouse model studies. S.G. has performed a single independent mice model study. A.B. analyzed the clinical data and revised the paper. K.S. did the histological preparations and examination. C.J. and Z.H. performed gnotobiotic mouse model studies. W.S. identified bacteria strains using MALDI-TOF.

## Graphical Abstract

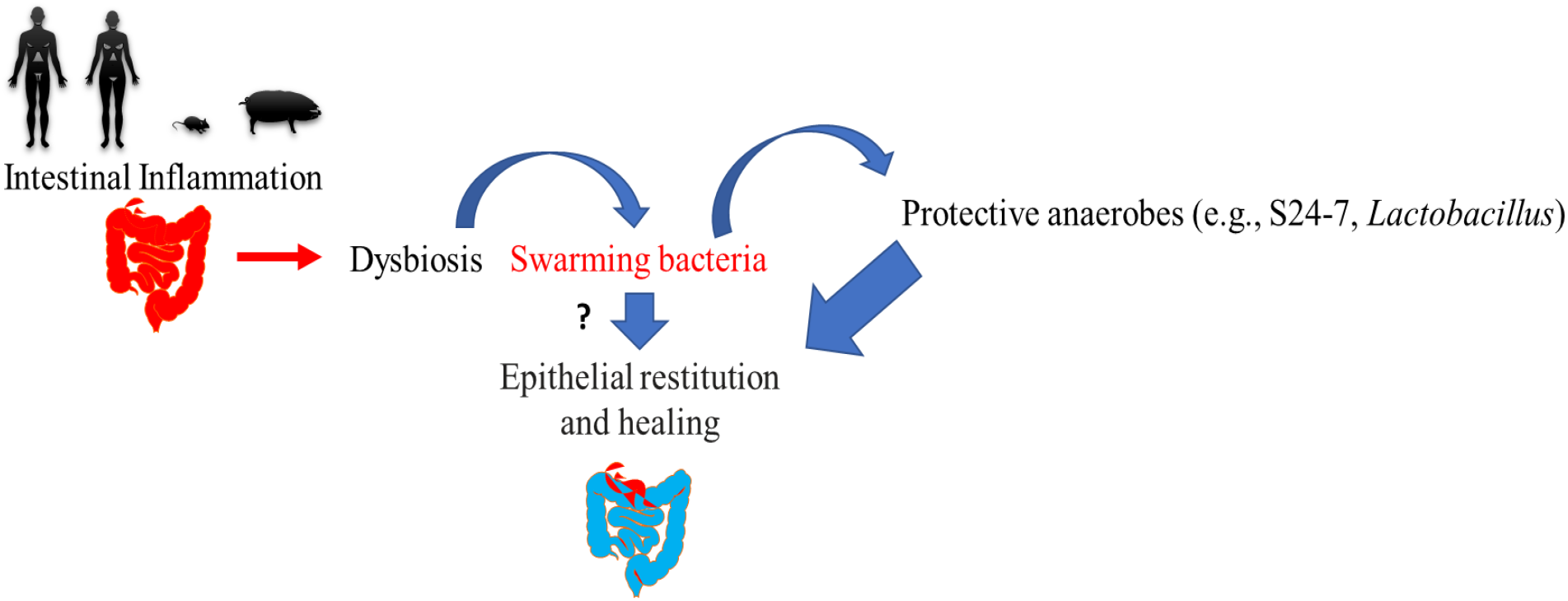
Schematic of proposed mechanism of cause and consequence of bacterial swarming during intestinal stress. Acute intestinal stress (e.g., colitis), as opposed to a homeostatic colonic lumen, induces growth of bacteria with swarming properties. If the abundance of swarming bacteria has reached sufficient levels passing some threshold CFU, it may exhibit epithelial restitution and healing by a direct or indirect mechanism and reduce local luminal oxygen levels in the intestine creating a favorable environment for enrichment of anaerobes. The bloom of protective anaerobes such as belonging to family Bacteriodales S24-7 and Lactobacillaceae in turn correlates with the accelerated resolution of inflammation and healing in colitic mice. Thus, bacterial swarming is a protective response to intestinal stress and one that is garnered during the evolution of colitis. Dashed line represents a possible unknown link, “?” represent an underlying possible direct mechanism; CFU, Colony Forming Units.

## STAR Method

### Clinical Study

From August 2014 through January 2018, sixty-three (63) patients were consented to participate in a colonoscopy aspirate or fecal collection study that was approved by the Institutional Review Board (IRB) (#2015-4465; #2009-446; #2007-554). The patients eligible for colonoscopy were enrolled sequentially after they provided study consent (#2015-4465; NCT 04089501). This study was audited by the IRB on April 24, 2019. All patients were screened and consented by a single gastroenterologist and Inflammatory Bowel Disease specialist (DL). Patients were enrolled if they had a diagnosis of inflammatory bowel disease (Crohn’s disease or Ulcerative colitis) or were undergoing routine screening colonoscopy for colorectal polyps/cancer or required a colonoscopy as part of their medical management of any gastrointestinal disorder as clinically indicated. The following information was collected in the clinic and codified (numerically) by the gastroenterologist (DL). Laboratory personnel receiving the aspirate sample were blinded to the patient, diagnosis and therapy outcome. The results of the swarming assay were then associated by un-blinding the clinical data (MB, DL). The clinical data collected included age, gender, pathology, other clinical diagnoses, and medications taken at the time of the colonoscopy. There were no dietary restrictions or special instructions for patients to follow prior to colonoscopy except for routine fasting prior to the procedure. The colonoscopy preparative regimen (split dose polyethylene glycol) was used for all patients. Patients (n = 62 of 63) successfully underwent complete colonoscopy and aspirates were taken from the region or mucosa of pathology or from descending colon while exiting from a grossly “normal” colonoscopy. One patient had a truncated colonoscopy due to incomplete cleansing and no specimen was obtained in this case. Specimens were collected in sterile fecal specimen cups without any preservative (Fisher, 650 mL), kept at 4°C for at most 15 minutes prior to transport to the laboratory. The specimens were then transported at ambient temperature (~ 15 minutes) to the laboratory for processing. Ten (10) de-identified random frozen fecal samples from pre-screened healthy volunteers were also obtained from OpenBiome (Boston, MA) (www.openBiome.org). Glycerol can facilitate swarming of bacterial cells on soft agar medium (Kim and Surette, 2005). Samples from OpenBiome are stored in glycerol. In order to avoid any external determinant that can influence swarming of bacterial species in these fecal samples, we washed the fecal samples in sterile PBS and then incubated in 2 mL LB broth overnight at 37°C, 200 rpm. Swarming assay was performed using these revived fecal cultures. Qualitative scoring of the swarmers in the clinical specimens was made based on the detection of bacterial spread with surfactant rim over a 72 hours incubation period. Samples showing swarming were scored as ≤ 24 hours (Score 3), 24-48 hours (Score 2) and 48-72 hours (Score 1). Samples that didn’t show swarming over 72 hours of incubation were considered as “non-swarmers” (Score 0).

### Isolation and identification of bacterial swarmers from feces

Swarming assay was performed on Luria Bertani (LB) swarming agar medium (10 g/L tryptone (Sigma), 5 g/L yeast extract (Acumedia) 10 g/L NaCl (Fisher), 5 g/L Agar (RPI)) with some modifications to an established method (Morales-Soto et al., 2015). To isolate a singular dominant swarmer from a polymicrobial mix of bacteria (such as feces), we initially focused on developing an assay to isolate swarmers using known polymicrobial mixed cultures of bacteria. Single bacterial species (up to seven strains belonging to different taxa) grown in LB [OD_600_ of 1.0-1.3] were mixed in a 1:1 ratio and, 5μL of this mix was spotted on 0.5% agar plates. Following air drying at room temperature, the plates were and incubated at 37°C, 40% RH (relative humidity) for 10 hours. Bacterial swarm front was swabbed using a sterile tooth-pick from the edge of swarming colony at different locations (see arrows, Fig. S2) and after re-streaking on separate agar plates and scaled by growth in LB, samples were identified using Matrix Assisted Laser Desorption and Ionization-Time of Flight (MALDI-TOF). Swarmers or hyperswarmers present in the fecal or colonoscopic samples were isolated and determined using an identical approach. Fecal pellets and/or colonoscopy aspirates from the clinic and/or feces of mice and pigs were collected in sterile tubes, and freshly prepared and used for swarming assays before freezing in small aliquots at −80°C. Feces from stressed intestinal model of pigs (Munyaka et al., 2016) were obtained on dry ice from Ehsan Khafipour at University of Manitoba Winnipeg, Canada. Five (5) μL of homogenized feces in Phosphate buffered saline PBS, pH 7.2 (100 mg/mL) or colonoscopic aspirates were spotted on swarming agar plates (optimized to 0.5% agar) and were incubated at 37°C and 40% RH, for 120 hours. Most bacterial swarmers, however, were detected within the first 48-72h from incubation. Dominant swarmers from the edge of the colony were identified using MALDI-TOF. Once identified, cells from the same aliquot were plated on to 1.5% LB agar and serially passaged from a single colony to obtain a pure culture of the strain.

### Bacterial swarming and time-lapse imaging

Swarming ability of a single bacterial species using a pure culture of *Enterobacter* sp. SM1 and its isogenic mutant, *Enterobacter* sp. SM3 and its transposon mutants, *Serratia marcescens* Db10 and JESM267, clinical isolate of *Serratia marcescens*, *Bacillus subtilis* 3610 and its isogenic mutant DS215 was always determined on LB swarming agar at 37°C and 40% RH prior to any experiments using these strains. *B. subtilis* 3610 and its isogenic mutant were compared on LB swarming agar containing 0.7% agar (Kearns and Losick, 2005). Briefly, 2 μL of an overnight culture grown in LB medium from a glycerol stock was spotted on swarming agar plate, followed by air drying at room temperature and incubating the plate overnight as stated above. Freshly made swarming agar plates, no more than 12 hours old when stored at 4°C, were used throughout the study. In order to capture real time swarming motility, a temperature and humidity-controlled incubator equipped with time lapse photography was built (see accompanying publication on Nature Protocol Exchange for detailed protocol, doi:10.21203/rs.2.9946/v1). As swarming is dependent upon RH (Kearns, 2010), we used an optimized RH of 40% that allowed image capturing without condensation on the lid of the Petridish. Unless otherwise stated, swarming potential of isogenic strains was always compared on the same swarming agar plate to nullify the difference due to the condition of the medium, which may vary between plates. Swarming area was calculated using a python-based script (available online, via Nature Protocol, doi:10.21203/rs.2.9946/v1) to identify the swarming edge using the time-lapse images. Swarming under anerobic condition (0.1 ppm) and 10% oxygen (3.3 ppm) were performed at 37°C and 40% RH in an anaerobic chamber (Coy Labs Inc.). In order to compare swarming motility between two strains, bacterial cultures were inoculated on a single plate and the colony areas were measured when the faster swarmer has covered half of the agar plate.

### Dextran Sulfate Sodium (DSS) induced acute colitis in conventional and gnotobiotic mice

Four to six-week old female C57BL/6 mice (Jackson Laboratories, Bar Harbor, ME; # 000664) were purchased and co-housed for acclimatization at The Albert Einstein College of Medicine vivarium for 2 weeks prior to randomization by coin toss as previously described (Venkatesh et al., 2014). To induce acute colitis, mice were administered 3% (w/v) DSS (MW 36-50 KDa) (MP Biomedicals, LLC; Cat. no. 160110) in animal facility drinking water throughout the course of the experiment. By contrast, the control group or the normal mice always received animal facility drinking water. To determine the effect of swarming and swarming deficient strains during colitis, mice were orally gavaged with 100 μL (~ 4×10^9^ CFU/mL) test bacteria or LB as vehicle, daily for 9-12 days until the weight of vehicle group dropped >20%. Daily gavage of bacterial strains absolutely required use of unwashed bacterial strains grown in fresh LB (OD_600_ ~ 1.0). In dose-optimization studies, bacterial dilutions were prepared in its own spent medium to vary cell number as the only determinant factor. Mice underwent daily monitoring for body weight, clinical signs and symptoms (e.g., occult blood, diarrhea, activity), gross water consumption (measuring water marked level), and visual inspection of rectal mucosa. At the end of the experiment, mice were euthanized using isoflurane anesthesia and intestines harvested for histopathology. The histology slides were prepared using a swiss role technique of intestines embedded in paraffin as previously published (Whittem et al., 2010). Scoring of inflammatory pathology was based on a published reference with minor modifications (Erben et al., 2014). The experimenter was not blinded to treatment allocation; however, the pathologist (K. S.) evaluating histologic scores was blinded to treatment allocation. All studies were approved by the Institute of Animal Studies at the Albert Einstein College of Medicine, INC (IACUC # 20160706 and preceding protocols).

Colitis was induced in five-week old germ-free (GF) wildtype (WT) C57BL/6 mice under specific pathogen free (SPF) conditions (McCafferty et al., 2013) using 3% DSS in drinking water and treated with 100 μL (~ 4×10^9^ CFU/mL) of test strain or LB as vehicle for 7 days when most mice had >10% weight drop. Mice were euthanized by CO_2_ asphyxiation and histology specimens were prepared using swiss roll technique. Scoring of inflammatory pathology was based on a published reference with minor modifications (Erben et al., 2014). Inflammation score and Trichrome Fibrosis score was calculated based on published references (Ding et al., 2012). All GF mouse protocols were approved by the Institutional Animal Care and Use Committee of the University of Florida (IACUC#201308038). The experimenter (ZH, CJ) was not blinded to treatment allocation; however, the pathologist (KS) evaluating histologic scores was blinded to treatment allocation. In addition, a second pathologist (QL), randomly re-read a subset of the colitis histology slides which had a high correlation with original pathologist read (KS) (Spearman ρ = 0.96, 95% CI 0.92 – 0.98, *P* < 0.0001, two-tailed).

### DSS-induced intestinal injury recovery model

In a study to determine the healing effect of SM3 in colitis, C57BL/6 mice were administered 3% DSS in drinking water for 7 days (when most mice had a weight loss >10% of their pre-DSS exposure weight). Subsequently, mice received animal facility drinking water without DSS and were further randomized by coin-toss to a treatment group delivered 4×10^9^ CFU/mL of bacterial cells or LB by oral gavage for 5 days. Colon samples were prepared for Hematoxylin-Eosin (H&E) staining and histology and processed as described above. The precise number of mice used for each experiment are stated in the Figure legends and are visible as separate plots in the graph. In the preliminary experiments (see Fig. 2d and 7c), the computed means (±SD’s) of the inflammation score for vehicle [7.28 –9.28 (+ 1.77 – 2.4)] and wildtype bacteria gavaged mice [2.59–5.66 (+ 1.31 – 3.2)] allowed for determination of the D value (~ ranging between 2-3). The exact sample size was determined under consideration of the D value, available mice per order, and ethical aspects (implementing replication studies, use of coin toss 1:1 randomization, and cage space at any given time point) as well as an assumed estimated inflation factor of ≤ 10%. For certain experiments, if insufficient number of mice were available for a reliable significance prediction, biologically independent repetition experiments were performed and data pooled for analysis (e.g., Fig. 2a-f, Fig. 7a-e).

### Construction of transposon mutants

In order to generate an isogenic swarming deficient strain of SM3, we adopted an *in vivo* transposition approach using pSAM_Ec with some modifications (Wiles et al., 2013). pSAM_Ec was a gift from Matthew Mulvey (Addgene plasmid #102939; http://n2t.net/addgene:102939; RRID: Addgene_102939). In short, donor strain – EcS17/pSAM_Ec was grown from an overnight culture in pre-mating medium (M9 salts containing 40μg/mL threonine and proline, 1μg/mL of thiamine) with 0.2% glucose until mid-exponential phase (OD_600_ 0.5-0.6). Similarly, the recipient strain SM3 was grown in pre-mating medium containing 0.4% lactose until early exponential phase (OD_600_ 0.2-0.3). After heat shock treatment of the recipient strain at 50°C for 30 minutes the cell density of SM3 was scaled up to obtain similar number of cells as that of the donor strain. For conjugation, 750 μL of both the strains were mixed, the cells were washed twice in M9 salts and re-suspended again in the same medium. The mixture of donor and recipient was placed on a sterile 0.45 μm membrane disc (Millipore) rested on mating agar plate (1x M9-thr-pro-thi-glucose agar) and incubated upright at 37°C overnight. Next day, the cells were dislodged from the membrane in M9 medium by vortexing and plated on selective agar medium (1x M9-threonine-glucose-kanamycin agar) containing kanamycin. Individual colonies were spotted on LB swarm agar plate to screen non-swarming or swarming deficient isogenic strain of SM3. The presence of transposon was confirmed by using transposon specific primer while the location of transposon insertion was verified by APPCR (Saavedra et al., 2017) followed by Sanger sequencing and mapping into the SM3 genome (STAR Method Table 1).

### 16S rRNA profiling to identify shift in colon microbiome

16S rRNA meta-analyses of the fecal samples from mice were conducted at Wright Labs, LLC. Fecal samples were shipped to Wright Labs, LLC on dry ice, and underwent DNA isolation using a Qiagen DNeasy Powersoil DNA Isolation kit following the manufacturer’s instructions (Qiagen, Frederick, MD). DNA was quantified and checked for its quality using the double stranded DNA high sensitivity assay on the Qubit 2.0 Fluorometer (Life Technologies, Carlsbad, CA). The 16S rRNA gene was amplified using Illumina iTag Polymerase Chain Reactions (PCR) based on the Earth Microbiome Project’s 16S rRNA amplification protocol (Walters et al., 2015). Amplified DNA was pooled, gel purified at ~400bp and multiplexed with other pure libraries to form a sequencing library normalized to the final concentration of library observed within each sample. The sequencing library was sequenced using an Illumina MiSeq V2 500 cycle kit cassette with 16S rRNA library sequencing primers set for 250 basepair (bp) paired-end reads at Laragen Inc (Culver City, CA).The paired-end sequences were merged with a minimum overlap of 200 bases, trimmed at a length of 251 bp, and quality filtered at an expected error of less than 0.5% using USEARCH (Edgar, 2010). The reads were analyzed using the QIIME 1.9.1 software package (Caporaso et al., 2010; Caporaso et al., 2011). Chimeric sequences were identified and assigned operational taxonomic units (OTU) using UPARSE at 97% identity (Edgar, 2013). The taxonomy was assigned using the Greengenes 16S rRNA gene database (13.5 release) (DeSantis et al., 2006). Linear discriminant analysis Effect Size (LEfSe) analysis was conducted to identify significantly enriched taxa within categorical groups of interest (Segata et al., 2011). For all comparisons, a Kruskal-Wallis alpha (α) was set at 0.05 to identify significantly enriched taxa, and a pairwise Wilcoxon rank sum test was utilized to test biological consistency across all subgroups (α = 0.05). Linear discriminant analysis (Walters et al., 2015) was calculated to determine effect size, and the 5 most strongly enriched taxa within each cohort were plotted. Co-occurrence network analysis was conducted on an unrarified OTU table containing bacterial abundance data from DSS+ SM3 treated samples and created within the Cytoscape plugin Conet (Faust and Raes, 2016); (Shannon et al., 2003). A spearman’s rho threshold of | 0.7 | was implemented prior to network plotting.

### Identification of bacterial strains isolated from a polymicrobial culture using MALDI-TOF

Bacterial cells picked from the leading edge of the swarming colony within the surfactant layer were sub-cultured on LB and then repeatedly sub-cultured onto MacConkey II, Columbia Nalidixic Acid (CNA) with 5% sheep blood, and Trypticase Soy Agar with 5% sheep blood plates (Becton Dickinson, Sparks, MD). Culture plates were incubated overnight at 35°C. A minimum of three colonies from each sample were identified by MALDI-TOF analysis using a MALDI Biotyper (Bruker Daltonics, Billerica, MA) in conjunction with Real Time Classification software (Bruker Daltonics, version 3.1). When evident, colonies with varied lactose fermentation reactions and/or colony morphologies were chosen for MALDI identification. Colonies were identified by directly transferring the bacteria to a MALDI target plate followed by the addition of 70% formic acid (Sigma-Aldrich, St. Louis, MO) and HCCA (α-cyano-4-hydroxycinnamic acid) matrix (Bruker Daltonics). When necessary, colonies with low MALDI identification scores (0-1.999) were sub-cultured, and a tube-based extraction was performed to attempt to improve the identification score. Briefly, colonies were added to 300 μL of water (Sigma-Aldrich) and emulsified followed by the addition of 100% ethanol (Sigma-Aldrich). The bacterial suspension was centrifuged (13,000 rpm, 2 minutes, RT) and the supernatant removed from the bacterial pellet. To extract the bacterial proteins, 50 μL of 70% formic acid and 50 μL of acetonitrile (Sigma-Aldrich) were added to the bacterial pellet, the sample was vigorously vortexed, and again centrifuged. The supernatant (1 μL) was spotted onto MALDI targets in triplicate for identification. MALDI identification scores of 1.7-1.999 were considered indicative of a reliable genus level identification whereas a MALDI score ≥ 2.0 indicated reliable genus and species, unless otherwise indicated.

### Bacterial strains and medium

All the bacterial strains used in this study were grown in LB broth (10 g/L tryptone, 5 g/L, 10g/L NaCl) at 37°C with agitation at 200 rpm, unless otherwise stated. Gram staining of SM1 and SM3 was performed using a commercially available Gram Staining kit (Becton Dickinson Microbiology Systems, Sparks, Maryland, USA) according to established protocol. For swarming, swimming and growth curve assays, bacterial strains were always grown from a glycerol stock stored at −80°C. Growth fitness of bacterial cells in LB at 37°C when grown with agitation was assessed spectrophotometrically at 600 nm using Beckman Spectrophotometer DU 640B. Growth curve was fitted using absorbance of culture as a function of time. Specifically, for growth analysis of SM1 and SM3, growth in LB was monitored at 600 nm every 20 minutes using a plate reader (PERKIN ELMER HTS7000) at 37°C for 24 hours.

### Measurement of microlevels of oxygen in mouse lumen

Oxygen concentration in the mouse lumen was assessed using a profiling oxygen microsensor (PresensIMP-PSt7-02) with a flat tip that has the ability to detect in the range of 0-1400 μM oxygen with an accuracy of ± 3%. The control or DSS treated mice were first anesthetized in isoflurane for at least 3 minutes, and then the microsensor probe was inserted from the anal verge. The oxygen concentration was monitored for one minute at different locations across the colon (0.5, 1 and 2 cm from the anus) using “Presens Measurement Studio 2 (version 3.0.1.1413)”. In order to avoid damage of the probe and mucosa while inserting through the anus, we used an Ethylenetetrafluoroethylene (ETFE) tube (outer diameter: 1mm; inner diameter: 0.7 mm) to house the probe. The housing was retracted to expose the probe in the designated location and cleaned before moving to the next location within the colon.

### Consumption of residual oxygen on swarming plates

Swarming plates were prepared as described previously and a fine hole (3 mm × 1mm) was made on the lid of the plate to fix a syringe-based oxygen microsensor probe (Presens, NTH-PSt7-02). After inoculation of the test bacterial culture, the probe was inserted into the swarming agar medium through the hole on the lid, finely adjusted using a manual micromanipulator (Presens) and then sealed using silicon oil. The side of the Petri dish was sealed using parafilm, and this whole unit was placed in the indigenously built environmental controlled incubator at 37°C. The oxygen consumption within the agar plate over time was monitored every 5 minutes for 20 hours using “Presens Measurement Studio 2”. The average oxygen consumption rate in a sealed container was calculated by dividing the change in oxygen concentration with time at which the oxygen levels reached a plateau phase. Consistently, we have observed that during swarming activity of SM3, the plateau phase stabilizes at an oxygen concentration of 0.003 ppm. This validated that the system used in this study was properly sealed from the outside environment.

### Swarming on mucosal surface

We used colon tissue from mice that had received 3% DSS water or water for 10 days to develop a mucosal race experiment. Normal or DSS treated mice were euthanized, and the large intestines were cut open and cleaned to remove residual feces. After rinsing thoroughly twice in 35% (v/v) ethanol and PBS, the intestines were sectioned into small segments of around 1.5-2.5 cm each. A hybrid plate with sterile swimming agar (3 g/L) and hard agar (15 g/L) was prepared, where one half of the plate had 1.5% agar and the other half was filled with 0.3% agar containing LB. To make such hybrid agar plate, 1.5% agar was poured first and once solidified half of the gel was removed using a sterilized spatula to fill the rest of the Petri dish with swimming agar. The tissue pieces were placed on 1.5% agar in a way so as to have one end of the tissue precisely overlapping with the border between 1.5% agar and the swimming agar. Overnight bacterial cultures were serially diluted 1012 times to reach cell concentration of 106 CFU/mL, 2 μL of which was inoculated on a 2 mm × 2 mm sterilized filter membrane (MF-Millipore, 0.45 μm). Bacterial cells adsorbed on membrane was then used as a source of inoculum on the mucosal surface. This avoided wetting of tissue surface that may facilitate free swimming and free flowing of bacterial cells on tissue surface. The motility of a swarming deficient and its wild type was always compared using a piece of tissue that belonged to the same region of the colon in mice. The plates were dried in the laminar hood for 20-30 minutes before incubating at 37°C and 40% RH overnight. Drying of plates allowed removal of excess moisture from the topmost layer of the tissue. Time-lapse photos were captured to evaluate the time at which bacterial test strain reached the other end of the intestinal tissue indicated by the swimming of bacteria on 0.3% LB agar. Distance travelled by the bacterial strain was measured in ImageJ according to the pixel/length ratio. The motility rates were calculated as Distance travelled / Time duration in which the test strain reached the swim agar.

### Swimming assays

Free swimming of bacterial cells was observed in fresh cultures that were grown in LB from an overnight culture (1:100 dilution in fresh LB) until OD_600_ ~ 0.3. At this point, cells were further diluted in PBS (1:50) and spotted on a glass slide with a cover slip placed on top of it. Swimming cells were observed under a phase contrast microscope (OMAX M837ZL, 40X) and videos captured using software (OMAX ToupView 3.7). The videos were captured at a frame rate of 18 fps for ~1 second for each trajectory and then processed in ImageJ (ver. 1.52g) to analyze swimming speeds of the test bacterium. Ten (10) straight trajectories of motion were picked randomly and the average speed was calculated as Trajectory length/Time.

For soft-agar swimming assay that may be relevant to *in vivo* conditions, 2 μL of overnight culture of the test bacterium was inoculated on LB swimming plate (10 g/L tryptone, 5 g/L, 10 g/L NaCl, 3 g/L Agar) and incubated at 37°C, 40% RH in our indigenously made incubator. To compare swimming potential of isogenic mutants, the isogenic pairs were inoculated on the same swim agar plate. Captured image in which the swimming colonies have not merged in due time, were used to calculate swim area using “Free selection” tool in ImageJ (ver. 1.59g).

### Genome sequencing, assembly, and annotation

We used a combination of short and long read sequencing to sequence and assemble two strains, SM1 and SM3. Short read library construction and sequencing was done at the New York Genome Center using a 2×300 paired end MiSeq run. Because we had many contigs for each strain, we next did PacBio® Single Molecule Real Time (SMRT) long read sequencing to improve our assemblies. SMRT sequencing was done at the Yale Center for Genome Analysis. The Hierarchical Genome Assembly Process (HGAP) was used for assembly of the genomes. SMRT sequencing and HGAP assembly yielded one contig for each strain. We then did a combined assembly with the PacBio and Illumina data using SPAdes (version 3.6.2) (Bankevich et al., 2012), using the PacBio contig with the ‘trusted_contigs’ parameter for each strain with otherwise default parameters. The CLC command clc_mapper (version 4.4.2.133896) was used to assess the quality of the final assembly for each strain. The final assembly for each strain was annotated using Prokka (version 1.3) (Seemann, 2014) annotating as “Bacteria” with otherwise default parameters.

### Genome comparisons

The SM1 and SM3 strains were compared to other available bacterial genomes using three approaches: 1) comparing the SM1/SM3 16S rRNA gene sequences to a large database of bacterial 16S sequences; 2) comparing conserved genes in SM1/SM3 to conserved genes in other bacterial genomes using multi-locus sequence typing (MLST); and 3) whole genome BLAST comparisons. Prokka identified eight 16S rRNA gene sequences in each strain. For 16S based analysis, these sequences were input to the Silva ACT: Alignment, Classification and Tree Service (Yilmaz et al., 2014). We used the “search and classify” and “compute tree” options with default parameters excepting that we restricted our search to one neighbor per query sequence. For MLST analysis, we input the SM1 and SM3 strains to autoMLST (Alanjary et al., 2019), an automated webserver that uses a precomputed set of marker genes to generate a multi locus species tree that placed SM1 and SM3 with their closest genomic neighbors. Both 16S and MLST analysis identified SM1 and SM3 as *Enterobacter* strains. We therefore used the BLAST Ring Generator (BRIG) (Alikhan et al., 2011) to compare the complete genomes of SM1 and SM3 with two clinically important close neighbors, *Enterobacter cloacae* subsp. cloacae ATCC 13047 and *Enterobacter asburiae* strain ATCC 35953. To run BRIG (Alikhan et al., 2011), we used the longest contig from SM1 (5107194 bp) as the reference genome and compared the other three genomes (SM3, *E. cloacae*, and *E. asburiae*) to SM1.

### Quantification of bacteria in feces using qPCR

To estimate species specific bacterial abundance (SM3, SM3_18, SM3_24, *B. subtilis* 3610 and DS215, *S. marcescens* Db10 and JESM267) present in murine feces, we approached a qPCR-based assay using total microbial DNA extracted from feces. DNA was extracted from feces samples collected on Day 0, 4 and 10 using DNeasy Powersoil kit (Qiagen). In case of glycerol stocks of feces, stocks were pelleted at 5,000 x g for 8 minutes, glycerol removed and re-suspended in 400 μL PBS solution. A total of 400 μL of the PBS/bacteria solution was then added to the lysing tube provided with the kit. SM3 specific primers (Just_F1 and Just_R1, Supplementary Table 2) were designed that did not show any amplification with *E. coli* DH5α and *Enterobacter cloacae* (the nearest neighbors by MLST). *Bacillus subtilis* and *Serratia marcescens* specific primers were adapted from previous studies (Bussalleu and Althouse, 2018; Fall et al., 2004). qPCR was performed using either Sensifast kit (Bioline) or PowerUp SYBR Green master mix (Applied Biosystem’s) following manufacturer’s protocol. Quantification of amplicon (copy number/μL) was calculated based on a standard curve generated using amplicon of interest as template DNA.

### Construction of isogenic mutants

Isogenic mutants of either *Enterobacter* sp. SM1 or *Salmonella enterica* serovar typhimurium were constructed using recombineering and PCR Ligation mutagenesis approach (Datta et al., 2006; Lau et al., 2002). Briefly, red recombineering plasmids pSIM5 or pSIM6 (graciously donated by Donald Court at National Cancer Institute, MD, USA) was electroporated in electrocompetent cells of SM1 or *Salmonella* (Electric field strength 12.5KV/cm & 200Ω) using Gene Pulser (Biorad). Bacterial cells harboring these plasmids were grown in LB from an overnight culture until the cell density reached OD_600_ 0.4-0.5. At this point Red recombinase proteins were induced by further incubating the cells at 42°C for 15 minutes followed by transferring the cells on ice for additional 15 minutes in order to make the cells electrocompetent. To make gene specific deletion a gene cassette containing FRT-Kan-FRT fragment or kanamycin resistance marker flanked by upstream and downstream gene sequences of the target gene was generated. Upstream and downstream regions were PCR amplified from genomic DNA, while FRT-Kan-FRT was amplified from the plasmid pKD4 as a template, using gene specific primers (Supplementary Table 2). Kanamycin resistance marker without FRT flanking region was amplified from pCAM48 (Hooven et al., 2016). Restriction digestion, ligation and PCR amplification using primers that can anneal at the 5’ end of the upstream region and 3’ end of the downstream region (upstream forward and downstream reverse primers) generated a gene cassette that can facilitate homologous recombination. Purified DNA was electroporated in bacterial cells in which lambda recombinase was expressed, followed by revival of cells in SOC medium at 30°C for 2 hours before selection of the clonal mutants on LB Agar containing 75 μg/mL kanamycin during an overnight incubation. At least two colonies were selected and transformed with flippase encoding pCP20 (Cherepanov and Wackernagel, 1995) in order to facilitate markerless gene deletion. Homologous recombination followed by excision of kanamycin resistance gene at the loci of interest was verified by Sanger sequencing.

### Qualitative measurement of surfactant production using blood agar hemolysis

Blood agar hemolysis assay was optimized based on previously established method (Walter et al., 2010). Briefly, 10 μL of overnight culture was inoculated on 15 mL Columbia Base agar plate supplemented with 2.5% defibrinated sheep blood. All the plates were incubated for 48h at 37°C, unless otherwise stated. The area of the zone of hemolysis was calculated using “Oval selection” tool in ImageJ.

### Qualitative measurement of surface tension using drop-collapse assay

Overnight bacteria culture was spun down at 4,500 rpm for 4 minutes. 5 μL of bacteria supernatant were pipetted on a polystyrene 96-well plate lid (Falcon). The droplet was placed at the center of a well and the lid was scanned (Bodour and Miller-Maier, 1998). Cross-sectional area of the droplet was calculated using “Free selection” tool in ImageJ.

### Qualitative measurement of surface tension using drop-counting assay

Drop-counting assay was performed with some modifications to an established method (Dilmohamud et al., 2005). Overnight bacteria culture was spun down at 4,500 rpm for 4 minutes. A glass Pasteur pipet was then used to transfer the supernatant to a plastic cuvette drop by drop. Number of droplets was counted for the spent medium supernatant to reach a total volume of 1 mL.

### Cytotoxicity of DSS on model intestinal human epithelia in the presence or absence of viable SM3 cells

To verify if SM3 can breakdown DSS affecting its cytotoxic efficacy, we treated Caco-2 cells (ATCC HTB-37), a colorectal adenocarcinoma cell line, with DSS that was incubated with SM3. Caco-2 cells were grown in complete medium: EMEM medium supplemented with 2mM L-glutamine, 20% Fetal bovine serum and 1X Anti-Anti (antimicrobial cocktail, Gibco) at 37°C, 5% CO_2_. When at confluence, cells were washed in sterile PBS (pH 7.2), trypsinized for 5 minutes and then resuspended in complete medium before seeding into 96 well plates. The seeding concentration was 30,000 cells/well. MTS assay (Cell Titer 96 Aqueous One Solution Cell Proliferation Assay, Promega) was used to determine cytotoxicity of DSS on Caco-2 cells. A dose– response assay using various concentrations of DSS in cell culture medium (8% - 0.03%, by serial dilution) showed a linear relation between DSS concentration and Caco-2 cytotoxicity. We chose to incubate SM3 cells in 8% DSS-medium. Bacterial cells (10^8^ CFU/mL), obtained from a fresh culture in LB (OD_600_ ≈ 0.1) after centrifugation followed by removal of spent medium, was re-suspended in 8% DSS-medium and incubated at 37°C for an hour with slow agitation. To avoid contamination of Caco-2 cells by SM3, the bacterial cells in DSS were heat killed at 95°C for 10 minutes and then spun down to collect the supernatant. 100 μL of this supernatant was used to treat Caco-2 cells grown in 96 well plate for durations of 1h, 12h and 24h. Similarly, heat killed SM3 (at 95°C for 10 minutes) prior to incubation in 8% DSS, 8% DSS alone, SM3 in complete medium, and complete medium alone were treated identically and used as controls in each assay. The cytotoxic efficacy was evaluated using MTS assay. This assay, at each time point, was performed in triplicate with three internal repeats. Cell survival (%) at each time point and in each condition is reported as a normalized value to the medium only at that specific time point.

### Tissue RT-qPCR

Colon total RNA (4 mice each group) were isolated using Trizol Reagent (Ambion) and reverse transcribed using High Capacity cDNA Reverse Transcription Kit (Thermofisher). RT-qPCR was performed using PowerUp SYBR Green Master Mix (Thermofisher) with primers of TNFα (FW: 5’-ATGAGAAGTTCCCAAATGGC-3’, RE: 5’-AGCTGCTCCTCCACTTGGTGG-3’), IL10 (FW: 5’-TGAGGCGCTGTCGTCATCGATTTC TCCC-3’, RE: 5’-ACCTGCTCCACTGCCTTGCT-3’), TNFR2 (FW: 5’-CAGGTTGTCTTGA CACCCTAC-3’, RE: 5’-GCACAGCACATCTGAGCCT-3’), IL6 (FW: 5’-GAGGTAAAAGA TTTACATAAA-3’, RE: 5’-CAAGATGAATTGGATGGTC-3’), and TBP (FW: 5’-ACCGT GAATCTTGGCTGTAAAC-3’, RE: 5’-GCAGCAAATCGCTTGGGATTA-3’). PCR was repeated in quadruplicate. The mRNA expression was normalized to internal control, TBP (TATA-Box Binding Protein). The entire experiment was repeated twice for reproducibility.

### Histologic scoring of mouse intestinal tissue exposed to *Salmonella*

Mice with 3% DSS (in drinking water)-induced colitis exposed to *S. enterica* serovar Typhimurium (day 1-5) were euthanized on day 5. Weights were measured every day and on day 5 intestinal tissue (SI, small intestines; colon) and liver was processed for histology (H&E stain). Scoring of degree of intestinal damage (n = 3 slides per treatment group; 1 slide per mouse) first followed using criteria established for colitis in a prior publication (Coburn et al., 2005) and consisted of determining extent of involvement and severity of luminal tissue (necrotic epithelial cells, polymorphonuclear leukocytes, and presence or absence of blood); mucosal lesions (extent of epithelial desquamation, ulceration, crypt abscess, mono- and polymorphonuclear leukocyte infiltrates and presence or absence of granulation tissue); and submucosal lesions (mononuclear leukocytes, polymorphonuclear leukocyte infiltrates, and edema). The small intestines were scored in similar fashion without determining crypt abscess formation which was absent. H-scores were computed by multiplying the severity of the lesion with extent (in terms of % of area involved in that slide for that lesion) (Geesala et al., 2019).

### Mucin Staining

Alcian Blue was used to stain mucin layer of intestine based on an established method described elsewhere (Dong et al., 2012). Briefly, intestinal tissues were deparaffinized and hydrated in distilled water. After mordanting in 3% Acetic Acid (v/v) for 3 minutes the tissue was stained in 1% Alcian Blue in 3% Acetic Acid (pH 2.5) for 30 minutes. Next, the specimens were dehydrated in 95% Ethanol and cleaned in xylene before mounting with Poly Mount.

### Lipocalin assay

LCN2 assay was performed using Mouse Lcn2/NGAL Duoset ELISA kit (R&D System, DY1857) according to manufacturer’s protocol. Briefly, fecal samples were collected and resuspended in PBS to a final concentration of 100 mg/mL. Samples were either diluted at a ratio of 1:10 (day 0 samples) or 1:1000 (day 10 samples) and added to precoated ELISA wells (100 μL/well). Plates were processed and read at an absorbance of 450 nm, corrected from optical imperfections by subtracting the reads using measurements at 570 nm, and the corrected reads were plotted against a standard curve to calculate lipocalin concentrations.

### RNA Extraction and metatranscriptome analysis

Cells were scraped from swarming plate and resuspended in 500μL RNA*Later* (Invitrogen) solution before freezing. Samples were sent to Wright Labs LLC. for further processing and analyses. RNA was extracted using the RNeasy PowerMicrobiome Kit (Qiagen). All RNA extracts were quantified using the Qubit RNA High Sensitivity Kit (Invitrogen, Carlsbad, CA, USA) to confirm complete DNase treatment of the RNA extracts (DNA concentration < 0.05 ng/uL). Subsequently, approximately 100 ng of extracted RNA was subject to NEBNext Ultra RNA Library Prep (New England BioLabs, Ipswich, MA, USA) for double stranded cDNA synthesis and RNA-Seq library preparation. Quality of the final library was assessed using a high sensitivity bioanalyzer chip (Agilent). The same library quantification, pooling, and purification methods were used as described above. Purified metatranscriptome libraries was sequenced using Illumina NextSeq. Raw read quality was assessed using the program FastQC to obtain average Q scores across the read length of all sequences and quality filtered using fastp. A sliding window filtration was utilized to cut reads at a 4-base average Q score of 28 or lower and reads trimmed below 60% of original length were discarded (Chen et al., 2018). Filtered RNA-Seq data were aligned against the SM3 genome using the Rockhopper RNA-Seq analysis platform using default parameters (Tjaden, 2015). Normalized expression counts were then subject to weighted Jaccard principal coordinates analyses within the QIIME2 software analysis package (Hall and Beiko, 2018). Analysis of similarity (ANOSIM) test statistics were calculated to assess the significance of differential clustering between categorical cohorts of interest.

### IL-10 receptor neutralization-induced chronic colitis

Mice deficient in Toll-like receptor 5 (*Tlr5*KO) were employed in this study given their heightened response to IL-10R neutralization-induced chronic colitis (Singh et al., 2015). Colitis was induced in 8-week-old female *Tlr5*KO mice by administering rat anti-IL-10R monoclonal antibody (1.0 mg/mouse, i.p.) (BioXcell) on Day 0 and 7. Mice were orally gavaged with SM1 or SM3 every third day from Day 1 onwards. Control mice were given isotype control antibody, *i.e.* rat anti-mouse IgG1 (BioXcell, West Lebanon, NH). Body weights were measured once every two days until euthanasia on Day 18. At euthanasia, blood was collected into BD microtainer (BD Biosciences, San Jose, CA) via cardiac puncture. Hemolysis-free sera were obtained after centrifugation and stored at −80°C until further analysis. Fecal samples were collected and prepared as described previously (Chassaing et al., 2012). Fecal lipocalin 2 (Lcn2), serum Lcn2 and serum keratinocyte-derived chemokine (KC) were quantified using Duoset ELISA kits from R&D Systems (Minneapolis, MN) according to the manufacturer’s protocol. Histology scoring for inflammatory damage was performed according to published criteria for colonic inflammation as a consequence of cytokine imbalance (Erben et al., 2014).

### *In vitro* co-culture assay using *M. intestinale*

To develop either a broth-based or swarm plate-based co-culture assay the growth kinetics of *Muribaculum intestinale* (DSM 28989) in Chopped meat carbohydrate broth, PR II (BD, BBL) in an anaerobic chamber, at 37°C (O_2_ = 1-2%) was determined (Fig. S7). Early exponential phase cells (OD_600_ 0.5-0.6) were used to establish the assay. For broth-based assay, 1mL of *M. intestinale* cells were mixed to 2μL of SM3/ SM3_18/ SM1 grown in LB medium and collected at mid-exponential phase, in a Hungate tube. Cells were collected at different time points (21/24, 36, 45/48 hrs), washed in PBS at 5000 rpm for 10 mins and total DNA extracted. DNA was resuspended in equal volume of TE buffer.

For swarm-plate based assay, 50mL overlay plates containing 30mL LB agar overlaid with 20mL LB swarming agar were prepared. An overnight culture of SM3/SM3_18/SM1 was prepared in LB and spotted, as mentioned before. *M. intestinale* grown in chopped meat medium was transferred into a bore-well at the center of the plate and incubated at different conditions for 64 hours (aerobic, sealed or anaerobic) at 37°C. For sealed condition, plates were taped carefully using parafilm to maintain anaerobiosis throughout the experiment. Plates incubated in aerobic conditions were maintained at a relative humidity of 50%. For Divided/Sealed condition, a small Petridish was placed inside a big Petridish and an overlay agar prepared then. The bore well containing *M. intestinale* was stationed in the small Petridish, while the swarming or less swarming strains were spotted on agar present in the big Petridish. Plates were sealed using parafilm, and the presence of an air space between the plates was ensured (please see caricature in Fig. 6). This allowed physical separation of *M. intestinale* from the swarming bacteria, nevertheless maintaining an anaerobic condition in the system. Plates were incubated at 37°C. At the end of the experiment, well containing cells were collected, washed twice in PBS and then proceeded further for DNA extraction using phenol-chloroform method. DNA was resuspended in equal volume of TE buffer. DNA samples were diluted at least 2 log fold to ascertain DNA copy number/μL within the range of standard. qPCR analysis was performed using equal volume of each diluted DNA sample and *M. intestinale* specific primers (Supplementary Table 2). Quantification of amplicon (copy number/μL) was calculated based on a standard curve generated using amplicon of interest as template DNA. The specificity of the primers was verified using SM3 genomic DNA in a qPCR reaction.

### Statistical analysis

P values of data were obtained by parametric or non-parametric methods, as indicated in the figure legends, with 95% confidence interval (CI). Normality (Gaussian distribution) was not assumed and for each dataset this was either tested for or transformed (e.g., log normality) to discern whether the data fit a Gaussian distribution. Through visual inspection, sample size assessment, and tests for normality, a determination was made to use a parametric or non-parametric statistical test, as indicated. All statistical tests, except where otherwise indicated, were performed with Graph Pad Prism v.8.2.0; * P < 0.05, ** P < 0.01, *** P < 0.001; ns, not significant. All plots are shown as mean and 95% CI except where otherwise indicated.

## STAR Method Tables

**STAR Method Table 1.**
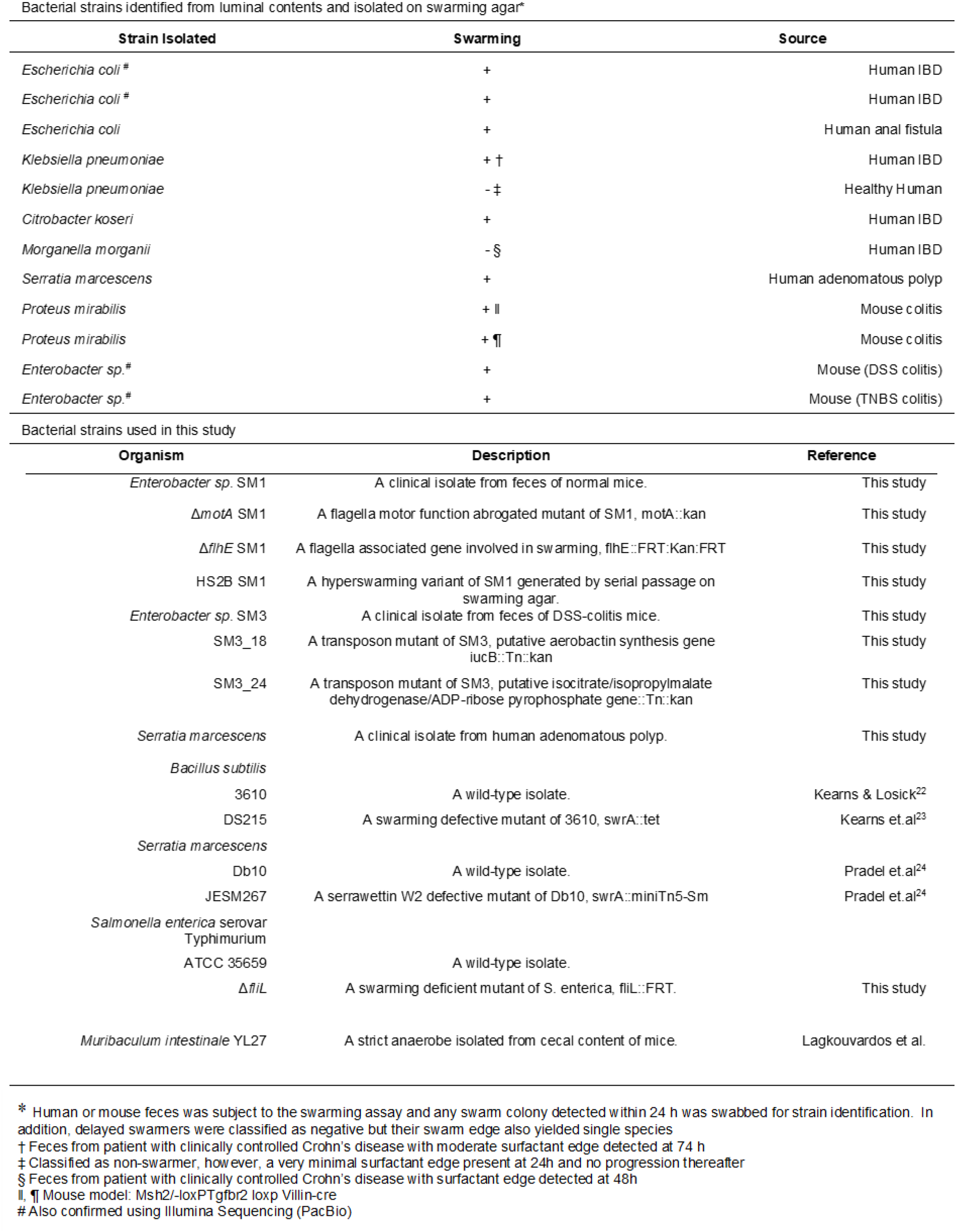
Bacterial Strains isolated and used in this study.

## References

Alanjary, M., Steinke, K., and Ziemert, N. (2019). AutoMLST: an automated web server for generating multi-locus species trees highlighting natural product potential. Nucleic Acids Res 47, W276–W282.

Alikhan, N.F., Petty, N.K., Ben Zakour, N.L., and Beatson, S.A. (2011). BLAST Ring Image Generator (BRIG): simple prokaryote genome comparisons. BMC Genomics 12, 402.

Allison, C., Emody, L., Coleman, N., and Hughes, C. (1994). The role of swarm cell differentiation and multicellular migration in the uropathogenicity of Proteus mirabilis. J Infect Dis 169, 1155–1158.

Angelini, T.E., Roper, M., Kolter, R., Weitz, D.A., and Brenner, M.P. (2009). Bacillus subtilis spreads by surfing on waves of surfactant. Proc Natl Acad Sci U S A 106, 18109–18113.

Bankevich, A., Nurk, S., Antipov, D., Gurevich, A.A., Dvorkin, M., Kulikov, A.S., Lesin, V.M., Nikolenko, S.I., Pham, S., Prjibelski, A.D., et al. (2012). SPAdes: a new genome assembly algorithm and its applications to single-cell sequencing. J Comput Biol 19, 455–477.

Barak, J.D., Gorski, L., Liang, A.S., and Narm, K.E. (2009). Previously uncharacterized Salmonella enterica genes required for swarming play a role in seedling colonization. Microbiology 155, 3701–3709.

Be’er, A., and Ariel, G. (2019). A statistical physics view of swarming bacteria. Mov Ecol 7, 9.

Be’er, A., Strain, S.K., Hernandez, R.A., Ben-Jacob, E., and Florin, E.L. (2013). Periodic reversals in Paenibacillus dendritiformis swarming. J Bacteriol 195, 2709–2717.

Bhuwan, M., Lee, H.J., Peng, H.L., and Chang, H.Y. (2012). Histidine-containing phosphotransfer protein-B (HptB) regulates swarming motility through partner-switching system in Pseudomonas aeruginosa PAO1 strain. J Biol Chem 287, 1903–1914.

Bodour, A.A., and Miller-Maier, R.M. (1998). Application of a modified drop-collapse technique for surfactant quantitation and screening of biosurfactant-producing microorganisms. Journal of Microbiological Methods 32, 273–280.

Borton, M.A., Sabag-Daigle, A., Wu, J., Solden, L.M., O’Banion, B.S., Daly, R.A., Wolfe, R.A., Gonzalez, J.F., Wysocki, V.H., Ahmer, B.M.M., et al. (2017). Chemical and pathogen-induced inflammation disrupt the murine intestinal microbiome. Microbiome 5, 47.

Burbank, L., Mohammadi, M., and Roper, M.C. (2015). Siderophore-Mediated Iron Acquisition Influences Motility and Is Required for Full Virulence of the Xylem-Dwelling Bacterial Phytopathogen Pantoea stewartii subsp stewartii. Applied and Environmental Microbiology 81, 139–148.

Butler, M.T., Wang, Q., and Harshey, R.M. (2010). Cell density and mobility protect swarming bacteria against antibiotics. Proc Natl Acad Sci U S A 107, 3776–3781.

Caiazza, N.C., Shanks, R.M., and O’Toole, G.A. (2005). Rhamnolipids modulate swarming motility patterns of Pseudomonas aeruginosa. J Bacteriol 187, 7351–7361.

Caporaso, J.G., Kuczynski, J., Stombaugh, J., Bittinger, K., Bushman, F.D., Costello, E.K., Fierer, N., Pena, A.G., Goodrich, J.K., Gordon, J.I., et al. (2010). QIIME allows analysis of high-throughput community sequencing data. Nat Methods 7, 335–336.

Caporaso, J.G., Lauber, C.L., Walters, W.A., Berg-Lyons, D., Lozupone, C.A., Turnbaugh, P.J., Fierer, N., and Knight, R. (2011). Global patterns of 16S rRNA diversity at a depth of millions of sequences per sample. Proc Natl Acad Sci U S A 108 Suppl 1, 4516–4522.

Chassaing, B., Aitken, J.D., Malleshappa, M., and Vijay-Kumar, M. (2014a). Dextran sulfate sodium (DSS)-induced colitis in mice. Curr Protoc Immunol 104, Unit 15 25.

Chassaing, B., Ley, R.E., and Gewirtz, A.T. (2014b). Intestinal epithelial cell toll-like receptor 5 regulates the intestinal microbiota to prevent low-grade inflammation and metabolic syndrome in mice. Gastroenterology 147, 1363–1377 e1317.

Chen, S., Zhou, Y., Chen, Y., and Gu, J. (2018). fastp: an ultra-fast all-in-one FASTQ preprocessor. Bioinformatics 34, i884–i890.

Cherepanov, P.P., and Wackernagel, W. (1995). Gene disruption in Escherichia coli: TcR and KmR cassettes with the option of Flp-catalyzed excision of the antibiotic-resistance determinant. Gene 158, 9–14.

Coburn, B., Li, Y., Owen, D., Vallance, B.A., and Finlay, B.B. (2005). Salmonella enterica serovar Typhimurium pathogenicity island 2 is necessary for complete virulence in a mouse model of infectious enterocolitis. Infect Immun 73, 3219–3227.

Colgan, S.P., and Taylor, C.T. (2010). Hypoxia: an alarm signal during intestinal inflammation. Nat Rev Gastroenterol Hepatol 7, 281–287.

Crespo-Sanjuán, J., Calvo-Nieves, M.D., Aguirre-Gervás, B., Herreros-Rodríguez, J., Velayos-Jiménez, B., Castro-Alija, M.J., Muñoz-Moreno, M.F., Sánchez, D., Zamora-González, N., Bajo-Grañeras, R., et al. (2015). Early detection of high oxidative activity in patients with adenomatous intestinal polyps and colorectal adenocarcinoma: myeloperoxidase and oxidized low-density lipoprotein in serum as new markers of oxidative stress in colorectal cancer. Lab Med 46, 123–135.

Cullender, T.C., Chassaing, B., Janzon, A., Kumar, K., Muller, C.E., Werner, J.J., Angenent, L.T., Bell, M.E., Hay, A.G., Peterson, D.A., et al. (2013). Innate and adaptive immunity interact to quench microbiome flagellar motility in the gut. Cell Host Microbe 14, 571–581.

Datta, S., Costantino, N., and Court, D.L. (2006). A set of recombineering plasmids for gram-negative bacteria. Gene 379, 109–115.

DeSantis, T.Z., Hugenholtz, P., Larsen, N., Rojas, M., Brodie, E.L., Keller, K., Huber, T., Dalevi, D., Hu, P., and Andersen, G.L. (2006). Greengenes, a chimera-checked 16S rRNA gene database and workbench compatible with ARB. Appl Environ Microbiol 72, 5069–5072.

Dilmohamud, B.A., Seeneevassen, J., Rughooputh, S.D.D.V., and Ramasami, P. (2005). Surface tension and related thermodynamic parameters of alcohols using the Traube stalagmometer. European Journal of Physics 26, 1079–1084.

Ding, S., Walton, K.L., Blue, R.E., McNaughton, K., Magness, S.T., and Lund, P.K. (2012). Mucosal healing and fibrosis after acute or chronic inflammation in wild type FVB-N mice and C57BL6 procollagen alpha1(I)-promoter-GFP reporter mice. PLoS One 7, e42568.

Dong, W., Matsuno, Y.K., and Kameyama, A. (2012). A procedure for Alcian blue staining of mucins on polyvinylidene difluoride membranes. Anal Chem 84, 8461–8466.

Edgar, R.C. (2010). Search and clustering orders of magnitude faster than BLAST. Bioinformatics 26, 2460–2461.

Edgar, R.C. (2013). UPARSE: highly accurate OTU sequences from microbial amplicon reads. Nat Methods 10, 996–998.

Erben, U., Loddenkemper, C., Doerfel, K., Spieckermann, S., Haller, D., Heimesaat, M.M., Zeitz, M., Siegmund, B., and Kuhl, A.A. (2014). A guide to histomorphological evaluation of intestinal inflammation in mouse models. Int J Clin Exp Pathol 7, 4557–4576.

Faust, K., and Raes, J. (2016). CoNet app: inference of biological association networks using Cytoscape. F1000Res 5, 1519.

Finkelshtein, A., Roth, D., Ben Jacob, E., and Ingham, C.J. (2015). Bacterial Swarms Recruit Cargo Bacteria To Pave the Way in Toxic Environments. mBio 6.

Geesala, R., Schanz, W., Biggs, M., Dixit, G., Skurski, J., Gurung, P., Meyerholz, D.K., Elliott, D., Issuree, P.D., and Maretzky, T. (2019). Loss of RHBDF2 results in an early-onset spontaneous murine colitis. J Leukoc Biol 105, 767–781.

Gude, S., Pince, E., Taute, K.M., Seinen, A.B., Shimizu, T.S., and Tans, S.J. (2020). Bacterial coexistence driven by motility and spatial competition. Nature 578, 588–592.

Hall, M., and Beiko, R.G. (2018). 16S rRNA Gene Analysis with QIIME2. Methods Mol Biol 1849, 113–129.

Hsu, C.C., Okumura, R., and Takeda, K. (2017). Human LYPD8 protein inhibits motility of flagellated bacteria. Inflamm Regen 37, 23.

Im, E., Riegler, F.M., Pothoulakis, C., and Rhee, S.H. (2012). Elevated lipopolysaccharide in the colon evokes intestinal inflammation, aggravated in immune modulator-impaired mice. Am J Physiol Gastrointest Liver Physiol 303, G490–497.

Jass, J.R. (2003). Hyperplastic-like polyps as precursors of microsatellite-unstable colorectal cancer. Am J Clin Pathol 119, 773–775.

Kearns, D.B. (2010). A field guide to bacterial swarming motility. Nat Rev Microbiol 8, 634–644.

Kearns, D.B., and Losick, R. (2005). Cell population heterogeneity during growth of Bacillus subtilis. Genes Dev 19, 3083–3094.

Koster, M., van Klompenburg, W., Bitter, W., Leong, J., and Weisbeek, P. (1994). Role for the outer membrane ferric siderophore receptor PupB in signal transduction across the bacterial cell envelope. EMBO J 13, 2805–2813.

Lagkouvardos, I., Lesker, T.R., Hitch, T.C.A., Galvez, E.J.C., Smit, N., Neuhaus, K., Wang, J., Baines, J.F., Abt, B., Stecher, B., et al. (2019). Sequence and cultivation study of Muribaculaceae reveals novel species, host preference, and functional potential of this yet undescribed family. Microbiome 7, 28.

Lau, P.C., Sung, C.K., Lee, J.H., Morrison, D.A., and Cvitkovitch, D.G. (2002). PCR ligation mutagenesis in transformable streptococci: application and efficiency. J Microbiol Methods 49, 193–205.

Lodes, M.J., Cong, Y., Elson, C.O., Mohamath, R., Landers, C.J., Targan, S.R., Fort, M., and Hershberg, R.M. (2004). Bacterial flagellin is a dominant antigen in Crohn disease. J Clin Invest 113, 1296–1306.

McCafferty, J., Muhlbauer, M., Gharaibeh, R.Z., Arthur, J.C., Perez-Chanona, E., Sha, W., Jobin, C., and Fodor, A.A. (2013). Stochastic changes over time and not founder effects drive cage effects in microbial community assembly in a mouse model. ISME J 7, 2116–2125.

Morales-Soto, N., Anyan, M.E., Mattingly, A.E., Madukoma, C.S., Harvey, C.W., Alber, M., Deziel, E., Kearns, D.B., and Shrout, J.D. (2015). Preparation, imaging, and quantification of bacterial surface motility assays. J Vis Exp.

Munyaka, P.M., Sepehri, S., Ghia, J.E., and Khafipour, E. (2016). Carrageenan Gum and Adherent Invasive Escherichia coli in a Piglet Model of Inflammatory Bowel Disease: Impact on Intestinal Mucosa-associated Microbiota. Front Microbiol 7, 462.

Okada, T., Fukuda, S., Hase, K., Nishiumi, S., Izumi, Y., Yoshida, M., Hagiwara, T., Kawashima, R., Yamazaki, M., Oshio, T., et al. (2013). Microbiota-derived lactate accelerates colon epithelial cell turnover in starvation-refed mice. Nat Commun 4, 1654.

Okada, T., Kanda, T., Ueda, N., Ikebuchi, Y., Hashiguchi, K., Nakao, K., and Isomoto, H. (2020). IL-8 and LYPD8 expression levels are associated with the inflammatory response in the colon of patients with ulcerative colitis. Biomed Rep 12, 193–198.

Okumura, R., Kurakawa, T., Nakano, T., Kayama, H., Kinoshita, M., Motooka, D., Gotoh, K., Kimura, T., Kamiyama, N., Kusu, T., et al. (2016). Lypd8 promotes the segregation of flagellated microbiota and colonic epithelia. Nature 532, 117–121.

Ormerod, K.L., Wood, D.L., Lachner, N., Gellatly, S.L., Daly, J.N., Parsons, J.D., Dal’Molin, C.G., Palfreyman, R.W., Nielsen, L.K., Cooper, M.A., et al. (2016). Genomic characterization of the uncultured Bacteroidales family S24-7 inhabiting the guts of homeothermic animals. Microbiome 4, 36.

Osaka, T., Moriyama, E., Arai, S., Date, Y., Yagi, J., Kikuchi, J., and Tsuneda, S. (2017). Meta-Analysis of Fecal Microbiota and Metabolites in Experimental Colitic Mice during the Inflammatory and Healing Phases. Nutrients 9.

Overhage, J., Bains, M., Brazas, M.D., and Hancock, R.E. (2008). Swarming of Pseudomonas aeruginosa is a complex adaptation leading to increased production of virulence factors and antibiotic resistance. J Bacteriol 190, 2671–2679.

Partridge, J.D., and Harshey, R.M. (2013). Swarming: flexible roaming plans. J Bacteriol 195, 909–918.

Perse, M., and Cerar, A. (2012). Dextran sodium sulphate colitis mouse model: traps and tricks. J Biomed Biotechnol 2012, 718617.

Rooks, M.G., Veiga, P., Wardwell-Scott, L.H., Tickle, T., Segata, N., Michaud, M., Gallini, C.A., Beal, C., van Hylckama-Vlieg, J.E., Ballal, S.A., et al. (2014). Gut microbiome composition and function in experimental colitis during active disease and treatment-induced remission. ISME J 8, 1403–1417.

Saavedra, J.T., Schwartzman, J.A., and Gilmore, M.S. (2017). Mapping Transposon Insertions in Bacterial Genomes by Arbitrarily Primed PCR. Curr Protoc Mol Biol 118, 15 15 11–15 15 15.

Sasaki, Y., Fukuda, S., Mikam, T., and Hada, R. (2008). Endoscopic quantification of mucosal surface roughness for grading severity of ulcerative colitis. Digestive Endoscopy 20, 67–72.

Seemann, T. (2014). Prokka: rapid prokaryotic genome annotation. Bioinformatics 30, 2068–2069.

Segata, N., Izard, J., Waldron, L., Gevers, D., Miropolsky, L., Garrett, W.S., and Huttenhower, C. (2011). Metagenomic biomarker discovery and explanation. Genome Biol 12, R60.

Selvam, R., Maheswari, P., Kavitha, P., Ravichandran, M., Sas, B., and Ramchand, C.N. (2009). Effect of Bacillus subtilis PB6, a natural probiotic on colon mucosal inflammation and plasma cytokines levels in inflammatory bowel disease. Indian J Biochem Biophys 46, 79–85.

Shannon, P., Markiel, A., Ozier, O., Baliga, N.S., Wang, J.T., Ramage, D., Amin, N., Schwikowski, B., and Ideker, T. (2003). Cytoscape: a software environment for integrated models of biomolecular interaction networks. Genome Res 13, 2498–2504.

Singh, V., Yeoh, B.S., Carvalho, F., Gewirtz, A.T., and Vijay-Kumar, M. (2015). Proneness of TLR5 deficient mice to develop colitis is microbiota dependent. Gut Microbes 6, 279–283.

Sokolov, A., and Aranson, I.S. (2009). Reduction of viscosity in suspension of swimming bacteria. Phys Rev Lett 103, 148101.

Solovyev, V. (2011). V. Solovyev, A Salamov (2011) Automatic Annotation of Microbial Genomes and Metagenomic Sequences. In Metagenomics and its Applications in Agriculture, Biomedicine and Environmental Studies (Ed. R.W. Li), Nova Science Publishers, p. 61–78. In, pp. 61–78.

Spencer, N.J., Dinning, P.G., Brookes, S.J., and Costa, M. (2016). Insights into the mechanisms underlying colonic motor patterns. J Physiol-London 594, 4099–4116.

Stanton, T.B., and Savage, D.C. (1984). Motility as a factor in bowel colonization by Roseburia cecicola, an obligately anaerobic bacterium from the mouse caecum. J Gen Microbiol 130, 173–183.

Suzuki, K., Arumugam, S., Yokoyama, J., Kawauchi, Y., Honda, Y., Sato, H., Aoyagi, Y., Terai, S., Okazaki, K., Suzuki, Y., et al. (2016). Pivotal Role of Carbohydrate Sulfotransferase 15 in Fibrosis and Mucosal Healing in Mouse Colitis. PLoS One 11, e0158967.

Tjaden, B. (2015). De novo assembly of bacterial transcriptomes from RNA-seq data. Genome Biol 16, 1.

Tran, H.Q., Ley, R.E., Gewirtz, A.T., and Chassaing, B. (2019). Flagellin-elicited adaptive immunity suppresses flagellated microbiota and vaccinates against chronic inflammatory diseases. Nat Commun 10, 5650.

Van Alst, N.E., Picardo, K.F., Iglewski, B.H., and Haidaris, C.G. (2007). Nitrate sensing and metabolism modulate motility, biofilm formation, and virulence in Pseudomonas aeruginosa. Infect Immun 75, 3780–3790.

Venkatesh, M., Mukherjee, S., Wang, H., Li, H., Sun, K., Benechet, A.P., Qiu, Z., Maher, L., Redinbo, M.R., Phillips, R.S., et al. (2014). Symbiotic bacterial metabolites regulate gastrointestinal barrier function via the xenobiotic sensor PXR and Toll-like receptor 4. Immunity 41, 296–310.

Volk, J.K., Nystrom, E.E.L., van der Post, S., Abad, B.M., Schroeder, B.O., Johansson, A., Svensson, F., Javerfelt, S., Johansson, M.E.V., Hansson, G.C., et al. (2019). The Nlrp6 inflammasome is not required for baseline colonic inner mucus layer formation or function. J Exp Med 216, 2602–2618.

Walter, V., Syldatk, C., and Hausmann, R. (2010). Screening Concepts for the Isolation of Biosurfactant Producing Microorganisms. Biosurfactants 672, 1–13.

Walters, W., Hyde, E.R., Berg-Lyons, D., Ackermann, G., Humphrey, G., Parada, A., Gilbert, J.A., Jansson, J.K., Caporaso, J.G., Fuhrman, J.A., et al. (2015). Improved Bacterial 16S rRNA Gene (V4 and V4-5) and Fungal Internal Transcribed Spacer Marker Gene Primers for Microbial Community Surveys. mSystems 1.

Whittem, C.G., Williams, A.D., and Williams, C.S. (2010). Murine Colitis modeling using Dextran Sulfate Sodium (DSS). J Vis Exp.

Wiles, T.J., Norton, J.P., Russell, C.W., Dalley, B.K., Fischer, K.F., and Mulvey, M.A. (2013). Combining quantitative genetic footprinting and trait enrichment analysis to identify fitness determinants of a bacterial pathogen. PLoS Genet 9, e1003716.

Wiles, T.J., Schlomann, B.H., Wall, E.S., Betancourt, R., Parthasarathy, R., and Guillemin, K. (2020). Swimming motility of a gut bacterial symbiont promotes resistance to intestinal expulsion and enhances inflammation. PLoS Biol 18, e3000661.

Yang, Y., and Jobin, C. (2014). Microbial imbalance and intestinal pathologies: connections and contributions. Dis Model Mech 7, 1131–1142.

Yilmaz, P., Parfrey, L.W., Yarza, P., Gerken, J., Pruesse, E., Quast, C., Schweer, T., Peplies, J., Ludwig, W., and Glockner, F.O. (2014). The SILVA and “All-species Living Tree Project (LTP)” taxonomic frameworks. Nucleic Acids Res 42, D643–648.

Yoon, S.H., Ha, S.M., Lim, J., Kwon, S., and Chun, J. (2017). A large-scale evaluation of algorithms to calculate average nucleotide identity. Antonie Van Leeuwenhoek 110, 1281–1286.

Young, G.M., Smith, M.J., Minnich, S.A., and Miller, V.L. (1999). The Yersinia enterocolitica motility master regulatory operon, flhDC, is required for flagellin production, swimming motility, and swarming motility. J Bacteriol 181, 2823–2833.

Zhang, H.P., Be’er, A., Florin, E.L., and Swinney, H.L. (2010). Collective motion and density fluctuations in bacterial colonies. Proc Natl Acad Sci U S A 107, 13626–13630.

