## Supplementary text and figures for "Bacterial Swarmers exhibit a Protective Response to Intestinal Stress"

#### Supplementary Information

#### Supplementary Text

**Commensal bacterial swarmers also abrogate intestinal stress.** To generalize this concept across multiple strains, mice with DSS induced colitis were administered *Bacillus subtilis* 3610 (wildtype)(Kearns and Losick, 2003) or its swarming deficient *swrA* isogenic mutant DS215 (Kearns et al., 2004) using the identical protocol as that used for SM3. In comparison with strain DS215, the wildtype significantly protected mice from intestinal inflammation (Fig. S5a-e). Similarly, swarming *Serratia marcescens* Db10, in contrast to the swarming deficient JESM267 isogenic mutant (Pradel et al., 2007), protected against inflammation in the identical mouse model (Fig. S5f-h). The bacterial levels in feces, collected on day 4 were not different compared to its respective mutant (Fig. S4c-d). Incidentally, a clinical strain of *S. marcescens* (isolated from the surface washing of a human dysplastic polyp) also protected against DSS induced inflammation in mice (Fig. S6). Together, our results confirm that from a diverse set of genes and pathways altered in different bacterial strains, the swarming phenotype of bacteria correlates with protection against inflammation.

***Enterobacter sp.* SM3 contributes in reduction of luminal oxygen concentration *in vivo*.** The enrichment of certain specific anaerobes when treated with SM3 suggested a reduction in oxygen content in the intestine; however, during inflammation, the median oxygen concentration in the lumen increases (Fig. S8a). Similarly, fecal 16S rDNA profile of GF/SPF mice exposed to DSS and treated with SM3 also showed enrichment of anaerobic and microaerophilic taxa compared to the vehicle group (Fig. S9). By contrast, the fecal microbiota of vehicle group was enriched in taxa that are aerobic and/or facultative anaerobic. Hence, we determined the oxygen concentrations within the intestinal lumen of mice at various lengths along the colon. In control conventional C57BL/6 mice, the colonic lumen is uniformly "hypoxic or anoxic". In colitic mice, however, we found a significant increase in the oxygen levels (ppm) in the colonic lumen (measured at different lengths from 0.5 to 2 cm proximal to the anal verge) (Fig. S8a). In DSS exposed mice treated with SM3, we observed a significant reduction in the luminal oxygen concentrations when compared to the mice that were treated with SM1 and the swarming deficient mutant strains (Fig. S8b-c). SM3_18 and SM3_24 did not significantly affect oxygen concentrations compared with vehicle control (Fig. S8c). In vitro experiments further proved that the swarming behavior of SM3 is dependent on oxygen concentration (Fig. S8d), which in turn reduces the oxygen levels in a closed system at a significantly higher rate than the slow swarming variants (Fig. S8e). Hence, we hypothesize that it is likely the act of swarming by SM3, *in vivo*, which might contribute to reducing the median oxygen concentrations in the intestinal lumen and concomitantly aid in establishing an anaerobic microenvironment. Indeed, the events leading to healing could also contribute to the reduced luminal oxygen levels.

**Anti-inflammatory effect is likely not due to surfactant production.** Swarming bacteria secrete surfactants, such as surfactin, which reduce surface tension during motility on a solid surface (Kearns, 2010). Surfactin is reported to attenuate TNBS induced colitis, possibly by differentially regulating anti-inflammatory and pro-inflammatory cytokines (Selvam et al., 2009). This finding leads us to speculate that higher levels of protection exhibited by SM3 in comparison to its identical strain SM1 could likely be due to differences in surfactin levels secreted by these strains. We used an indirect blood hemolysis readout assay to test for the presence or absence of surfactin (or equivalent surfactants with surfactin-like activity). In this assay, blood agar hemolysis by SM1 and SM3 demonstrated similar zones of hemolysis. This observation suggested that the expression of surfactins might be similar in SM1 and SM3, at least under the conditions tested. We observed similar results for *Bacillus subtilis* 3610 and its isogenic *swr*A mutant, and *S. marcescens* Db10 and its swarming deficient mutant JESM267 at 37°C (Fig. S13). Notably, wildtype *S. marcescens* and JESM267 show differences in the zone of hemolysis at 30°C but not at 37°C on blood agar plates. However, in the context of mice *in vivo* experiments, the hemolysis phenotype exhibited by strains at 37°C *in vitro* is more relevant in extrapolating to the *in vivo* condition (i.e. body temperature is ~37°C) (Newsom et al., 2004). Indeed, quantification of genes essential for surfactin synthesis (*srf*AA) and surfactin secretion (*sfp*) by RT-qPCR did not show any significant difference between *B. subtilis* 3610 and the non-swarming strain DS215 (data not shown). Similar quantification for SM1 and SM3 could not be done as the genes responsible for surfactin synthesis or production are not characterized yet. A BLAST search for homologous genes in SM1 identified a putative *sfp* gene. However, in-frame deletion of this gene did not show any difference in surfactin production as estimated using blood agar assay as well as in its swarming potential compared to SM1 (data not shown). In addition to the blood agar assay, drop-collapse assay (Bodour and Miller-Maier, 1998) and drop-counting assay based on modified Stalagmometric

Method (Dilmohamud et al., 2005) did not show any significant difference between the isogenic pairs of SM3, *B. subtilis* 3610 and *S. marcescens* Db10 (Fig. S13d-e).

**Genome characterization.** Genome assembly generated one major contig of 5,107,194 bp both in the case of *Enterobacter sp.* SM1 and *Enterobacter sp.* SM3 (NCBI BioProject PRJNA558971) with 56% GC content. Other contigs were substantially smaller (<1500 bp). Prokka (Seemann, 2014) annotation revealed 85 tRNAs and 25 rRNAs encoded by the genome of SM1 and SM3, and 4608 CDS for SM1 and 4618 CDS for SM3, respectively. Progressive Mauve (Darling et al., 2010) alignment of genome sequences found 7 potential SNPs, among which two are non-synonymous mutations. Notably, there are several mutations also present on 16S rDNA sequence (Supplementary Table 1). Based on Multi Locus Sequence Typing (MLST) our strains were found to be closest to *Enterobacter asburiae*. Considering four close sequences from the MLST generated tree (*Enterobacter mori, Enterobacter kobei, Lelliottia nimipressuralis,* and *Enterobacter asburiae*), we did ANI analysis using OrthoANI (Yoon et al., 2017) and found ~96% identity ANI between SM1/SM3 and *E. asburiae*.

### Supplementary Figures

#### Figure S1

**a**

Top

Middle

Bottom

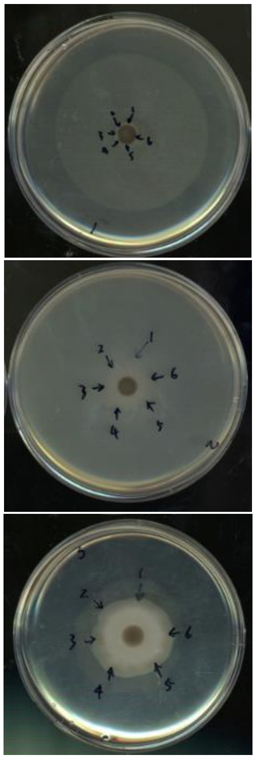

**b**

| Sample Number | MALDI Identification | MALDI Identification Score (Mean) * |
|---|---|---|
| A-1 | *Klebsiella pneumoniae*<br>*Escherichia coli* | 2.26<br>2.06 |
| A-2 | *Klebsiella pneumoniae*<br>*Escherichia coli* | 2.24<br>2.33 |
| A-3 | *Klebsiella pneumoniae*<br>*Escherichia coli*<br>*Acinetobacter sp.* | 2.20<br>2.33<br>1.82 |
| A-4 | *Klebsiella pneumoniae*<br>*Acinetobacter sp.* | 2.12<br>1.76 |
| A-5 | *Klebsiella pneumoniae* | 2.12 |
| A-6 | *Klebsiella pneumoniae*<br>*Escherichia coli* | 2.24<br>2.14 |
| B-1 | *Bacillus sp.* | 1.81 |
| B-2 | *Bacillus sp.* | 1.83 |
| B-3 | *Bacillus sp.* | 1.79 |
| B-4 | *Bacillus sp.* | 1.80 |
| B-5 | *Bacillus sp.* | 1.80 |
| B-6 | *Bacillus sp.* | 1.84 |
| C-1 | *Enterobacter asburiae*† | 2.24 |
| C-2 | *Enterobacter asburiae* | 2.28 |
| C-3 | *Enterobacter asburiae* | 2.31 |
| C-4 | *Enterobacter asburiae* | 2.25 |
| C-5 | *Enterobacter asburiae* | 2.27 |
| C-6 | *Enterobacter asburiae* | 2.21 |

* MALDI identification scores ≥ 2.0 indicate reliable species level identification. Identification scores between 1.7-1.99 indicate a reliable identification to the genus level only.

† *Enterobacter asburiae* is a member of *Enterobacter cloacae* complex. The species level identification *of E. cloacae* complex members by MALDI was confirmed using 16S sequencing.

**Figure S1 │ Identification of dominant swarming bacteria within a polymicrobial culture. a,** 1:1 ratio mix of bacteria were used for swarming assay on 0.5% LB agar for 10 hours. *Top*: five non-swarming bacteria were mixed and applied on 0.5% agar. Six random picks as shown in arrows were placed on the edge of colony (1. *Klebsiella pneumoniae* 2. *Escherichia coli* 3. *Acinetobacter* sp. 4. *Bordetella hinzii* 5. *Staphylococcus xylosus*) and 1.5% LB agar streaks performed - single viable colonies were subjected to MALDI-TOF identification. *Middle*: five non-swarming bacteria as above plus two known swarming bacteria SM3 (*Enterobacter asburiae*) and *Bacillus* sp. were mixed, and experiment repeated as per *Top* panel. Six random picks as shown in arrows were placed on the edge of the complex. *Bottom*: five non-swarming bacteria as above plus one known swarming bacteria SM3 (*Enterobacter asburiae*). Six random picks as shown in arrows were placed on the edge of complex**. b,** Table showing results of MALDI-TOF identification of bacterial colonies isolated from swarming edge. A1-A6 are picks from **a** *Top*. B1-B6 from **a** *Middle*, and C1-C6 from **a** *Bottom*. "A" represents mix of bacterial species *Klebsiella pneumoniae, Escherichia coli, Acinetobacter* sp., and *Bordetella hiuzii*, *Staphylococcus xylosus*; "B" represents mix of "A", *Bacillus pumilus*, and SM3; "C" represents mix of "A" and SM3.

#### Figure S2

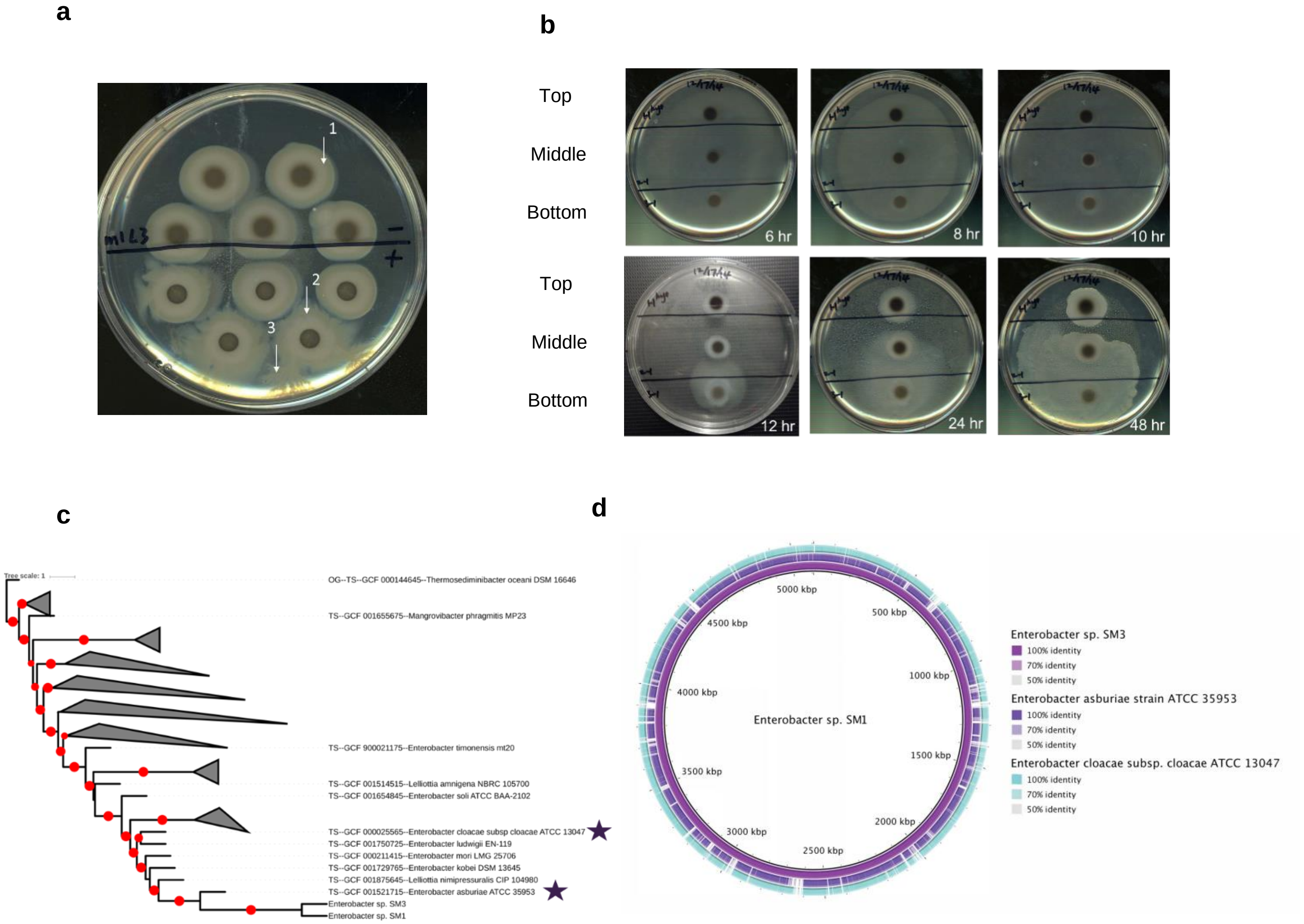


**Figure S2 │Isolation and characterization of *Enterobacter* sp. a,** Five replicate fecal spots from pooled fecal pellets of mice administered water (above black line) or 3% DSS water (below black line) (n = 3, day 7). The white arrows indicate 1, swarm edge isolation from control feces (SM1); 2, swarm edge isolation from feces of mice exposed to DSS (SM2); 3, swarm colony isolation from spontaneous "burst" activity from feces at 24h from plating (SM3). The mouse experiments were repeated at least twice. **b,** The bacterial clones isolated from **a** were re-plated as pure strains on 0.5% LB agar and the swarming assay performed over time. Two solid black marker lines divide each plate into 3 regions, holding spots of the 3 strains – Top: Strain 1 (SM1), Middle: Strain 2 (SM2), Bottom: Strain 3 (SM3). These strains have been repeatedly (≥25 times) plated in swarming assays from all aliquots stored from the original isolation (August 2014) and the results confirm that SM3 is a stable hyperswarmer. **c**, Phylogenetic tree showing multi-locus sequencing typing-based genetic relatedness between *Enterobacter* sp. SM1, SM3 and reference genomes. Tree was generated with autoMLST (CITE) and drawn using iTOL (CITE). Red dots indicate bootstrap support > 0.8. Stars represent related strains used for comparison with the genome sequences of SM1 and SM3 in panel **d**. **d,** Genome comparison of related *Enterobacter* strains. *Enterobacter*sp. SM1 was compared to *Enterobacter*sp. SM3 (purple) and the related strains *Enterobacter asburiae* ATCC 35953 (violet) and *Enterobacter cloacae* ATCC 13047 (cyan), and plotted in BLAST Ring Generator (BRIG) http://brig.sourceforge.net/ PMID: 21824423. DSS, Dextran Sulfate Sodium; LB, Luria-Bertani broth.

### Figure S3

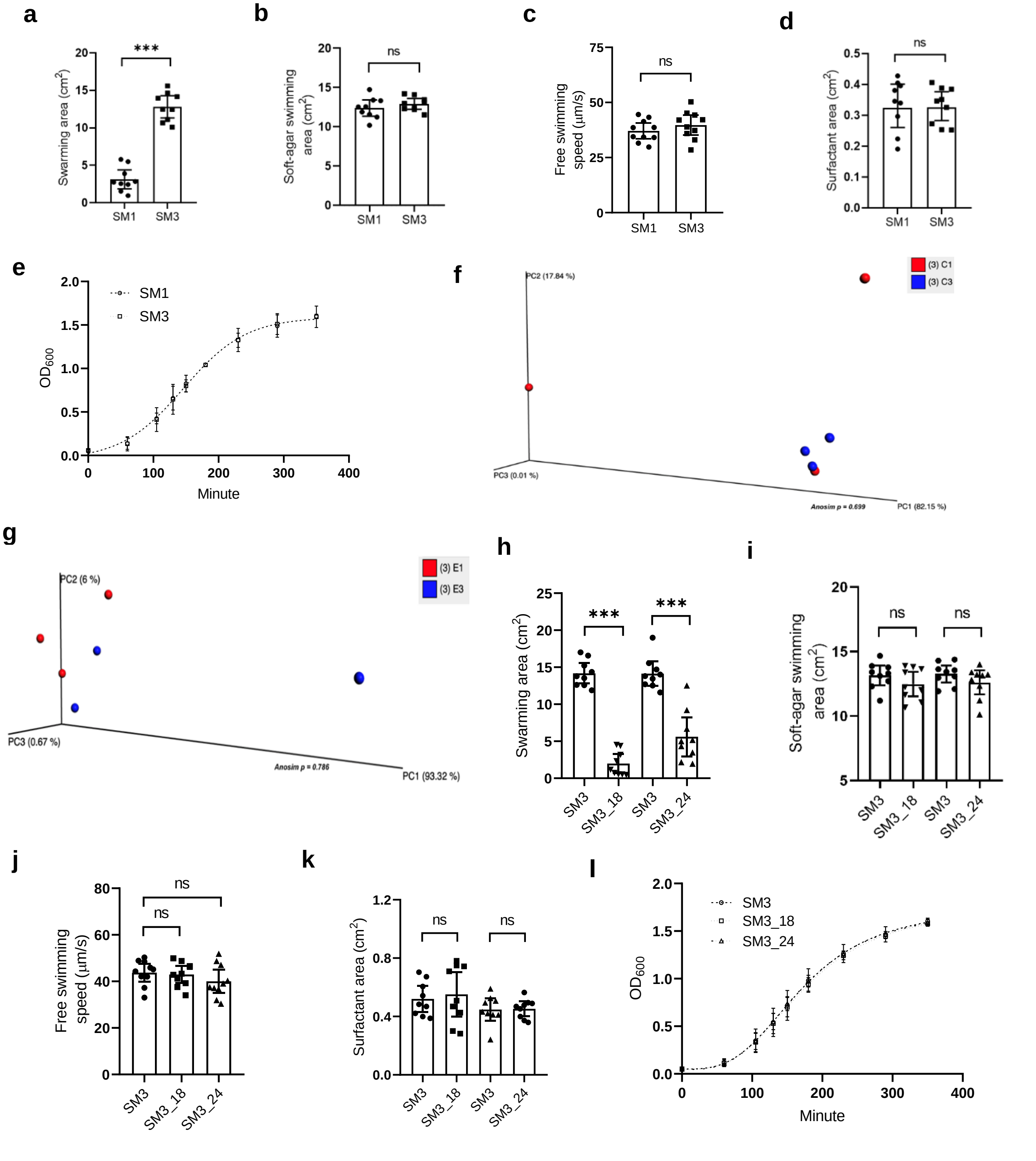


**Figure S3 │ Characterization of motility, growth, and surfactant production by *Enterobacter* sp. SM1, SM3 and its mutant strains. a-e,** SM3 and SM1, swarming motility (**a**), soft-agar swimming motility (**b**), free swimming motility (**c**), surfactant production (**d**) and growth rate (**e**) (n = 3, each in triplicate except for **e**, n = 3, each in singlet). **f-g,** Principal Coordinate Analysis (PCoA) plots of weighted Jaccard distance generated to determine global differences in expression of virulence and multi-drug resistance associated genes between cells collected from the center **(f)** and edge **(g)** of swarming colonies of SM1 and SM3 on agar. C1, center of SM1; C3, center of SM3; E1, edge of SM1 and E3, edge of SM3. **h-l,** SM3 and mutants (SM3_18 and SM3_24), swarming motility (**h**), soft-agar swimming motility (**i**), free swimming motility (**j**), surfactant production (**k**), and growth rate (**l**) (n = 3, each in triplicate except for **j**, n = 3, each in singlet). Unless otherwise noted, data are presented as mean and 95% CI, and significance tested using a two-tailed Student's t-test. **f-g,** significance tested using ANOSIM showing no significant difference between the tested groups $p = 0.699$ (**f**) and 0.786 (**g**). **j,** significance tested using one-way ANOVA followed by Tukey's post hoc test.

### Figure S4

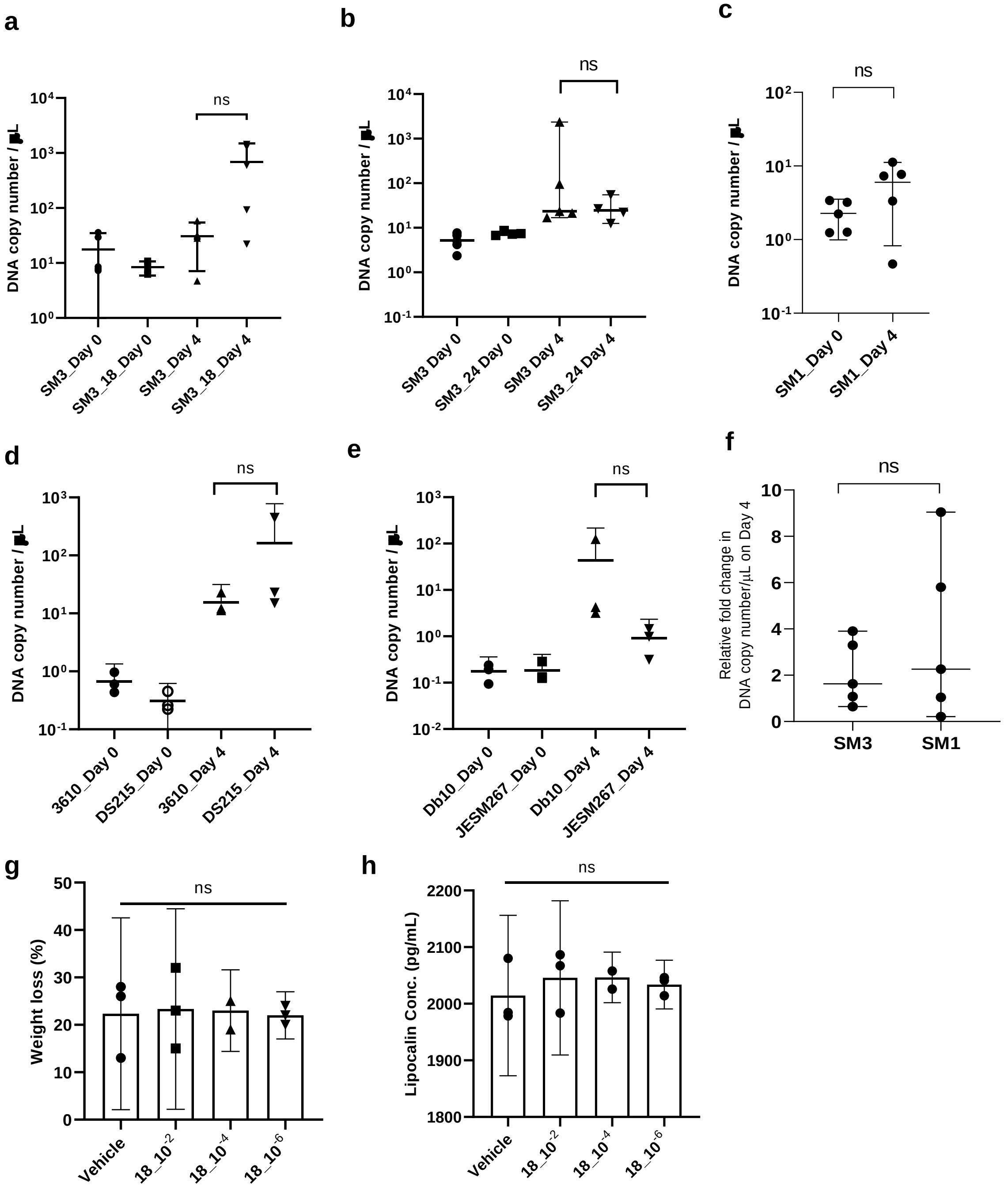


**Figure S4│Bacterial enumeration in feces using qPCR and dose-optimization effect of SM3_18 on DSS induced colitis in C57BL/6 mice**. **a-d**, 8-week old mice were exposed to DSS water for 4 days. Total DNA was extracted from feces collected on day 0 and day 4, processed and assessed using qPCR. 5ng of total DNA in conjunction with strain specific primers were used to quantify bacterial copy numbers. In each assay, DNA copy number/μL was calculated based on an internal standard curve and compared between samples from mice treated with SM3 and SM3_18 (**a**), SM3 and SM3_24 (**b**), SM1 (**c**), *B. subtilis* 3610 and DS215 (**d**) and *S. marcescens* Db10 and JESM267 (**e**) ($n \geq 3$ mice per treatment group, each performed as triplicate technical repeats). **f,** Relative abundance of SM3 or SM1 in feces on Day 4 when compared to Day 0, represented as ratio of DNA copy number/μL. **g-h**, In a separate experiment, 8-week old mice were exposed to DSS water and treated with different dilutions (10-2, 10-4, 10-6) of SM3_18 culture, which was grown in LB until 3 hours (O.D.600 ≈ 1.0), for 10 days. **e**, Weight loss (n = 3 mice per treatment group) and **f**, Lipocalin concentration (pg/mL) (n = 3, each in triplicates). Data are represented as mean and 95% CI, and significance tested using Fisher's Exact test unless otherwise stated. c, significance tested using Paired t-test. f, significance tested using Unpaired t-test with Welch's correction.

#### Figure S5

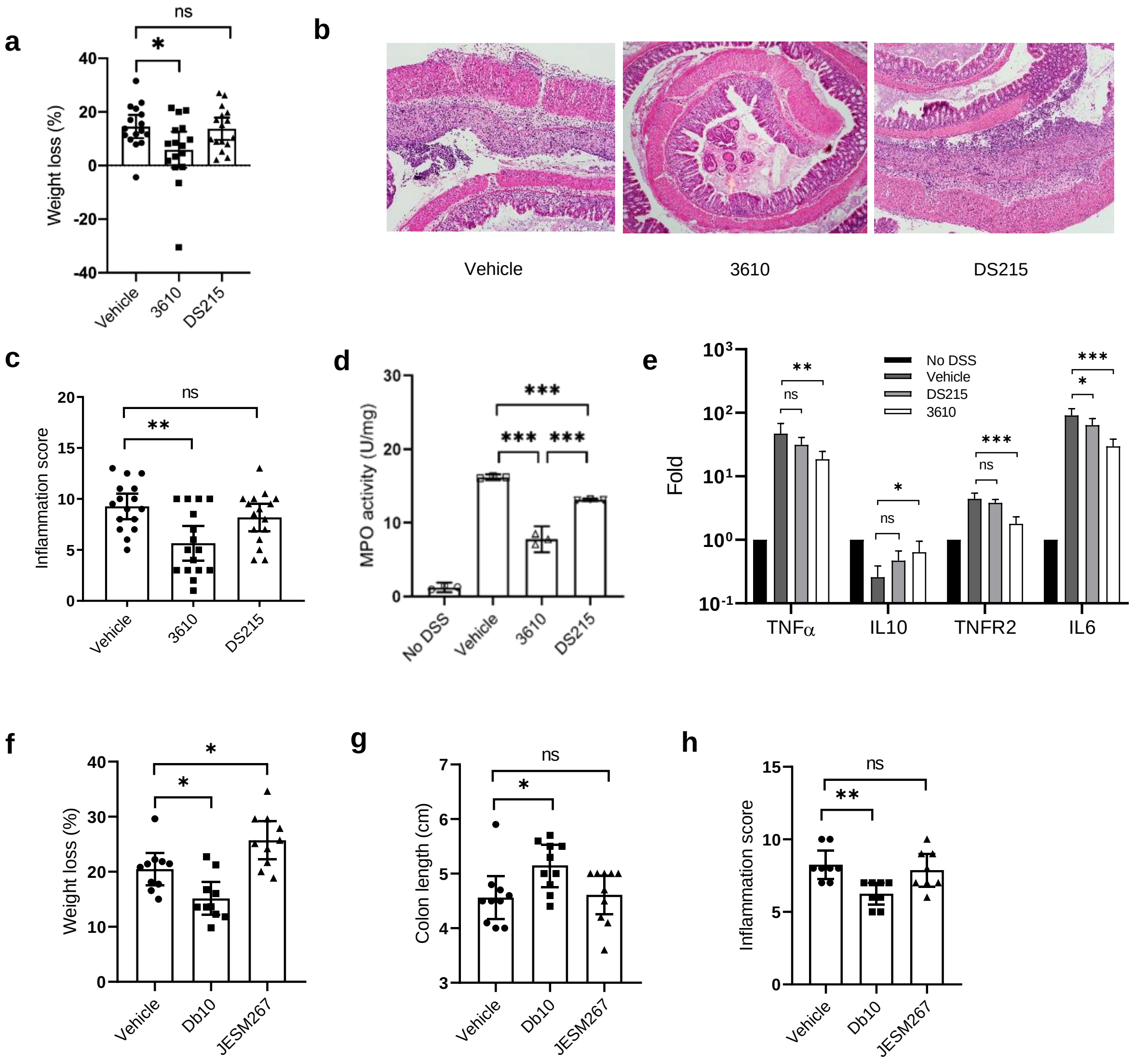


**Figure S5│Effects of *B. subtilis* and *S. marcescens* on DSS induced colitis in C57BL/6 mice. a-e,** 8-week old mice were exposed to DSS water and treated with vehicle (LB), *B. subtilis* 3610 or *B. subtilis* DS215 by oral gavage for 10 days. **a,** Weight loss (n = 16 per treatment group). **b,** Representative images (100x magnification) of H&E stained colonic section treated with vehicle (left), 3610 (middle) and DS215 (Borton et al.). **c,** Inflammation score (n = 16 per treatment group). **d-e,** In a separate experiment, myeloperoxidase (MPO) enzyme activity was determined (n = 3, each in duplicate) (**d**). Colon total RNA (n = 4) were isolated and reverse transcribed to cDNA. RT-qPCR data show fold induction of mRNA (TNFα, IL10, TNFR2, IL6). PCR was repeated in quadruplicate. The expression was normalized to internal control, TBP. The entire experiment was repeated n = 2 for reproducibility (**e**). **f-h,** In a separate experiment, C57BL/6 mice (8-week old) were exposed to DSS water and administered vehicle (LB), *S. marcescens* Db10 or *S. marcescens* JESM267 for 10 days. **f-h** indicates weight loss (**f**), colon length (**g**) and inflammation score (**h**) (n = 10 per treatment group except for **h**, for which n = 8; two colon specimens per group were used for other experiments). Unless otherwise noted, data represented as mean and 95% CI, and significance tested using one-way ANOVA followed by Tukey's post hoc test. **g,** data represented as median and interquartile range, and significance tested using Kruskal-Wallis followed by Dunn’s multiple comparisons test.

### Figure S6

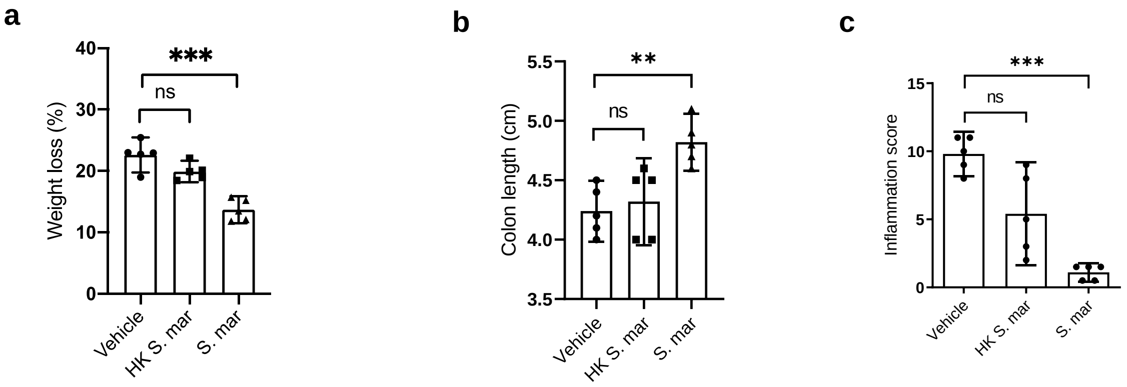


**Figure S6 | Effects of *Serratia marcescens* on DSS-induced colitis in C57BL/6 mice.** 8-week old mice were exposed to DSS water and treated with vehicle (LB), *S. marcescens* or heat killed *S. marcescens* by oral gavage for 10 days. **a**-**c** indicates weight loss (**a**), colon length (**b**), and inflammation score (**c**) (n = 5 per treatment group). Data are represented as mean and 95% CI, and significance tested using one-way ANOVA followed by Tukey's post hoc test. HK, Heat killed; S. mar, *S. marcescens*.

### Figure S7

**a**

| Query id | Subject id | Bit score | Evalue | query length | % identity | mismatches | subject length | alignment length |
|---|---|---|---|---|---|---|---|---|
| *OTU_5* | Muribaculum_intestinale__YL27__KR364784 | 130 | 5.71E-35 | 80 | 97.368 | 2 | 1460 | 76 |
| *OTU_3882* | Muribaculum_intestinale__YL27__KR364784 | 124 | 2.66E-33 | 80 | 96.053 | 3 | 1460 | 76 |
| *OTU_9* | Muribaculum_intestinale__YL27__KR364784 | 119 | 1.24E-31 | 80 | 94.805 | 2 | 1460 | 77 |
| *OTU_4541* | Muribaculum_intestinale__YL27__KR364784 | 119 | 1.24E-31 | 80 | 94.737 | 4 | 1460 | 76 |
| *OTU_5360* | Muribaculum_intestinale__YL27__KR364784 | 119 | 1.24E-31 | 80 | 94.737 | 4 | 1460 | 76 |
| *OTU_1* | Muribaculum_intestinale__YL27__KR364784 | 113 | 5.75E-30 | 80 | 93.421 | 5 | 1460 | 76 |
| *OTU_2569* | Muribaculum_intestinale__YL27__KR364784 | 113 | 5.75E-30 | 80 | 93.421 | 5 | 1460 | 76 |
| *OTU_5379* | Muribaculum_intestinale__YL27__KR364784 | 115 | 1.60E-30 | 80 | 92.5 | 6 | 1460 | 80 |
| *OTU_4806* | Muribaculum_intestinale__YL27__KR364784 | 91.6 | 2.69E-23 | 80 | 88.312 | 7 | 1460 | 77 |

**b**

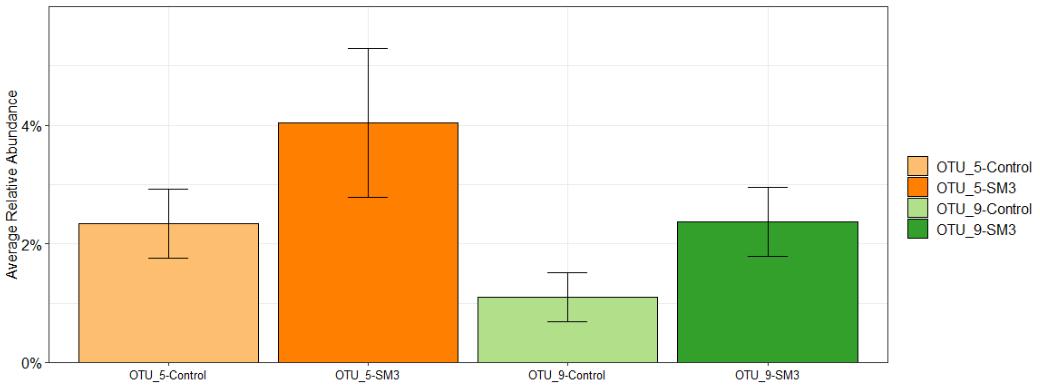


**c**

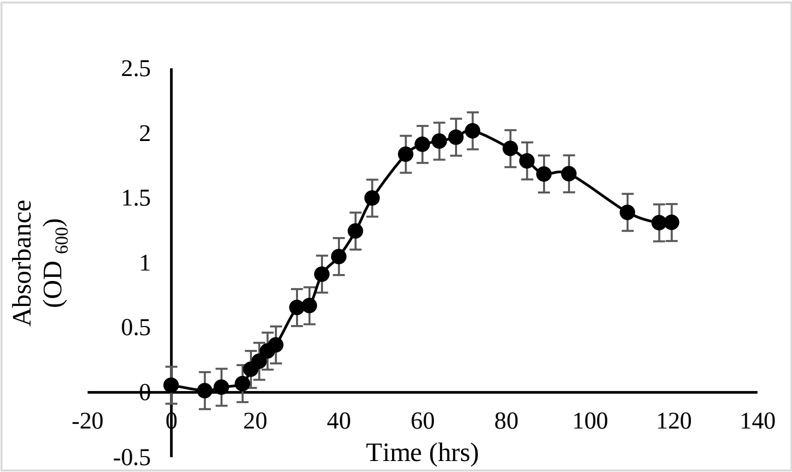


**Figure S7│In vitro growth kinetics and sequence homology of *Muribaculum intestinale* YL27 to 16S rDNA profile of feces from SM3 treated DSS- induced colitic mice. a,** Percent identity of *M. intestinale* 16S sequence to S24-7 specific OTU's identified in feces of SM3 treated mice using BLAST. **b,** Comparison of relative abundance of OTU_5 (sharing 97.34% identity and 2 mismatches to *M. intestinale*) and OTU_9 (sharing 94.8% identity and 2 mismatches to *M. intestinale*) in feces from mice treated or untreated with SM3 (n = 10). **c**, Growth kinetics of M. intestinale in Chopped meat carbohydrate medium, PR II at 37$^0$C in an anaerobic chamber ($O_2$ = 1-2%) (n = 2 independent replicate). Data represented as Mean (±SD).

### Figure S8

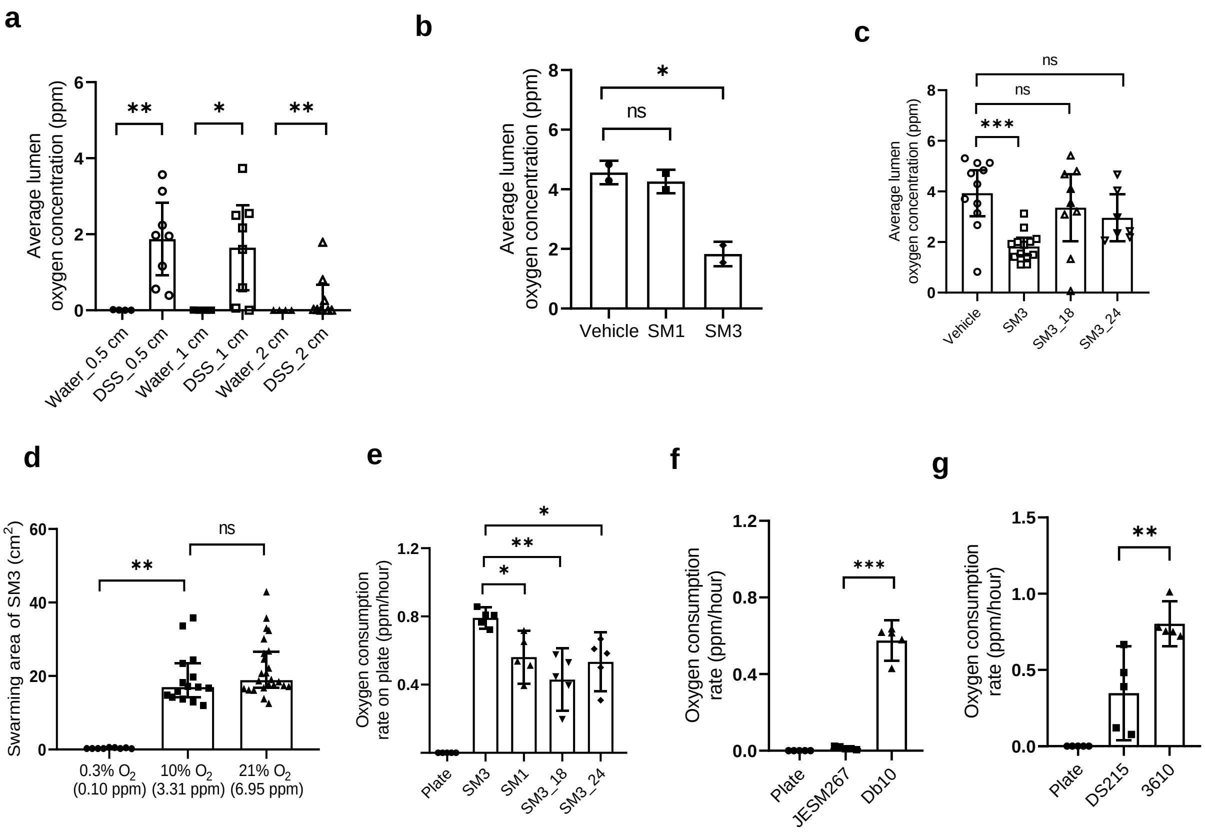


**Figure S8 │ Oxygen measurements *in vivo* and *in vitro* using a microsensor probe. a,** C57BL/6 mice were exposed to water or DSS water for 10 days. Average lumen oxygen concentration (0.5, 1, and 2 cm from the anus) was measured (normal, n = 4; DSS, n = 8). **b,** C57BL/6 mice were exposed to DSS water and treated with SM3 and SM1 for 10 days. Average lumen oxygen concentration was measured in a single experiment (n = 2). **c,** In a separate experiment, C57BL/6 mice were exposed to DSS water and treated with SM3 or its mutants (SM3_18 or SM3_24) for 10 days. Average lumen oxygen concentration was measured (n = 3, at least 2 mice each separate experiment). **d**, The swarming area of SM3 on LB agar plate in 8 hours under different concentration of oxygen (0.3%: n = 3, each in triplicate; 10%: n = 5, each in triplicate; 21%: n = 6, each in quadruplicate). **e-g,** Oxygen consumption rate was measured for different strains: SM1, SM3, and its mutant strains (**e**); Db10 and JESM267 (**f**); 3610 and DS215 (**g**) on LB agar plate (n = 5, each in singlet). Plate indicates oxygen consumption rate in LB agar with no bacteria. Unless otherwise noted, data are presented as mean and 95% CI, and significance tested using one-way ANOVA followed by Tukey's post hoc test. **a,** data are presented as median and interquartile range, and significance tested using Kruskal-Wallis test. **b,** for 0.5 cm and 1 cm groups, significance tested using a two-tailed Student’s t-test; for 2 cm groups, data are presented as median and interquartile range, and significance tested using Mann Whitney test.

### Figure S9

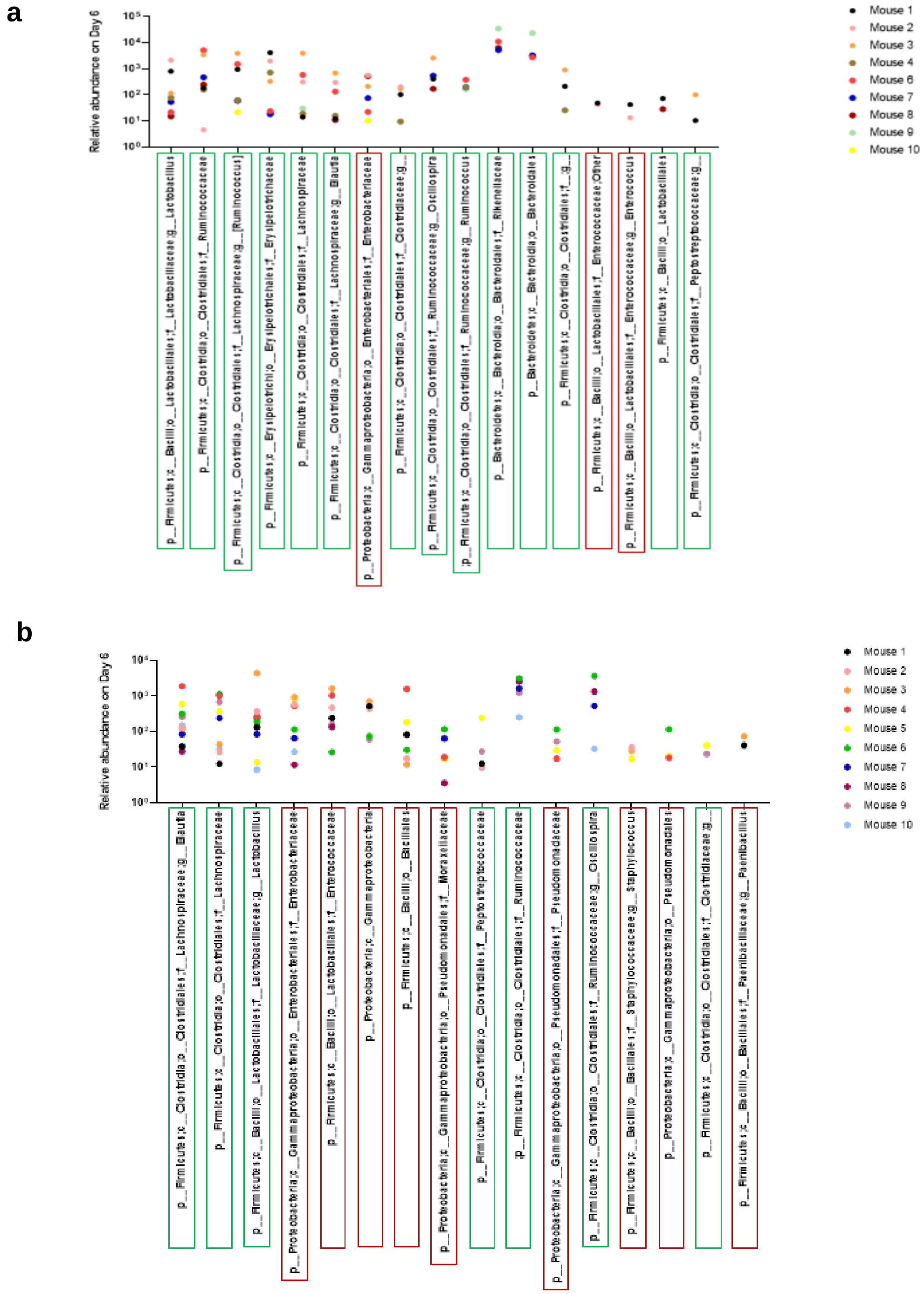


**Figure S9 │ Relative abundance of different taxa in the feces of gnotobiotic mice. a-b,** 16S rDNA profiling of feces samples from GF/SPF mice exposed to DSS and treated with SM3 **(a)** or vehicle **(b)** for 6 days were analyzed. The ratio of each taxa (represented as OTU normalized to total OTU's per sample) on Day 6 to its normalized OTU values in feces collected on Day 0 or Day 3, in each individual mouse, was calculated. All taxa with an abundance ratio > 10 and found at least in two individual mice are presented. Enrichment of these taxa indicates favorable conditions that allowed its growth during the course of the experiment, *in vivo*. Taxa highlighted in boxes are either microaerophilic and anaerobic (green) or aerobic or facultative anaerobic (Venkatesh et al.).

### Figure S10

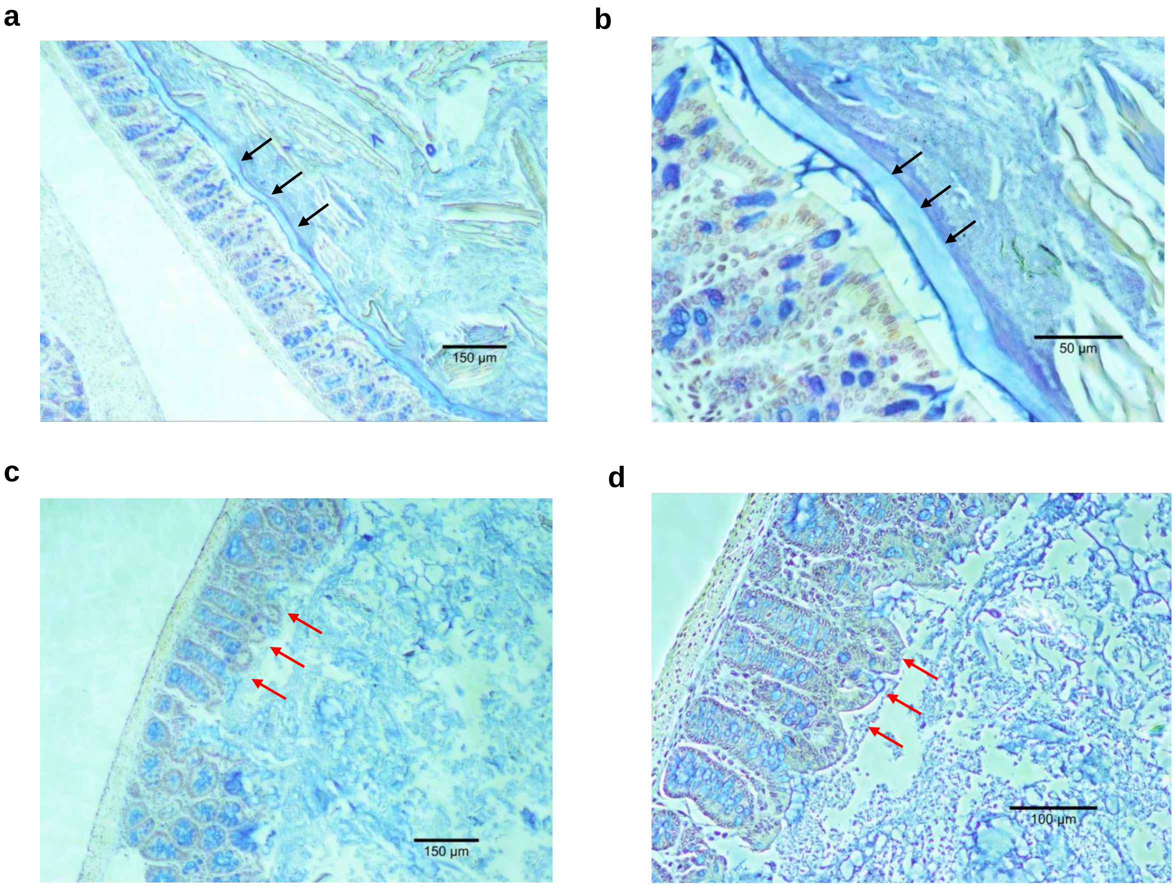


**Figure S10 │ Representative images of mucin stained mice intestinal tissue.** Large intestine obtained from 8-week old C57BL/6 mice exposed to water (**a, b**) or DSS water (**c, d**) for 10 days and stained for mucin using Alcian Blue. **a-b,** Black arrows indicate the mucin layer on normal large intestine. **c-d.** Red arrows indicate the loss of mucin layer on colitic tissue. **a, c** 10x objectives; **b,** 40x objectives; **d,** 20x objectives.

#### Figure S11

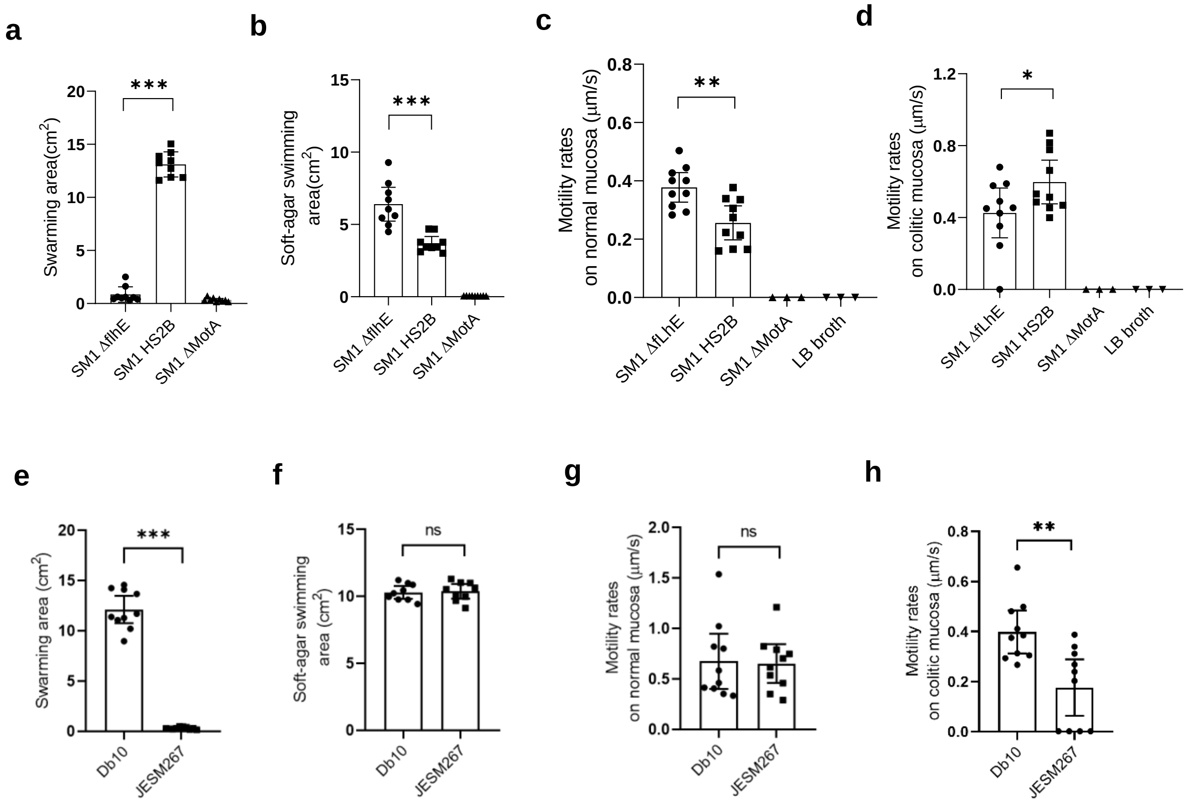


**Figure S11 │ Motility rates on different media. a-b,** *ΔflhE* SM1, HS2B SM1 and Δ*motA* SM1 swarming motility (**a**), soft-agar swimming motility (**b**) (n = 3, each in triplicate). **c-d,** *ΔflhE* SM1, HS2B SM1, Δ*motA* SM1 and LB spotted as negative control, motility rates on normal (**g**) and colitic (**h**) mucosal surface of C57BL/6 mouse (n = 10, and n = 3 for Δ*motA* SM1, LB). **e-f,** *S. marcescens* Db10 and *S. marcescens* JESM267, swarming motility (**e**), soft-agar swimming motility (**f**) (n = 3, each in triplicate). **g-h,** *S. marcescens* Db10 and *S. marcescens* JESM267, motility rates on normal (**g**) and colitic (**h**) mucosal surface of C57BL/6 mouse (n = 4, at least in duplicate). Unless otherwise noted, data are represented as mean and 95% CI, and significance tested using a two-tailed Student's t-test.

### Figure S12

a

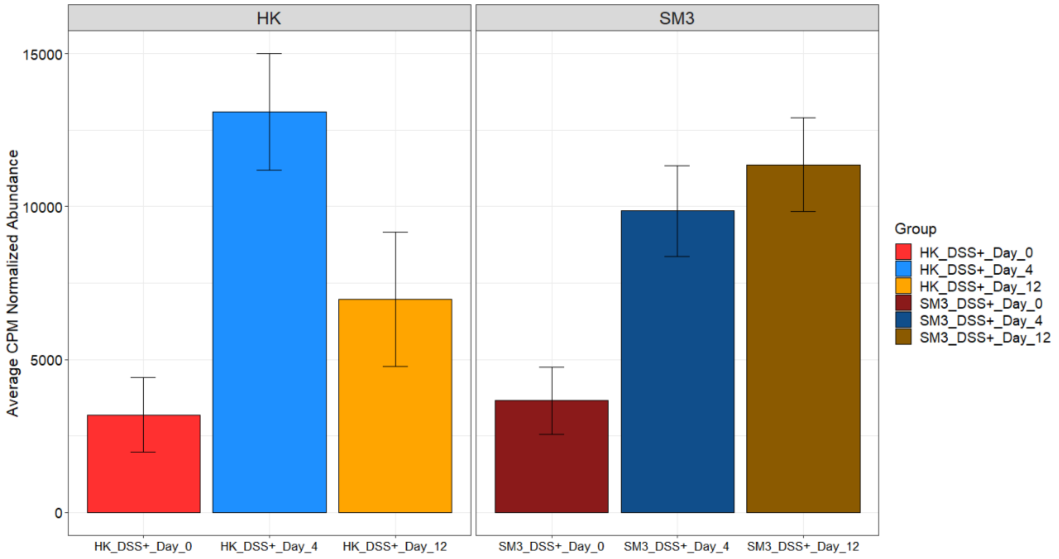


b

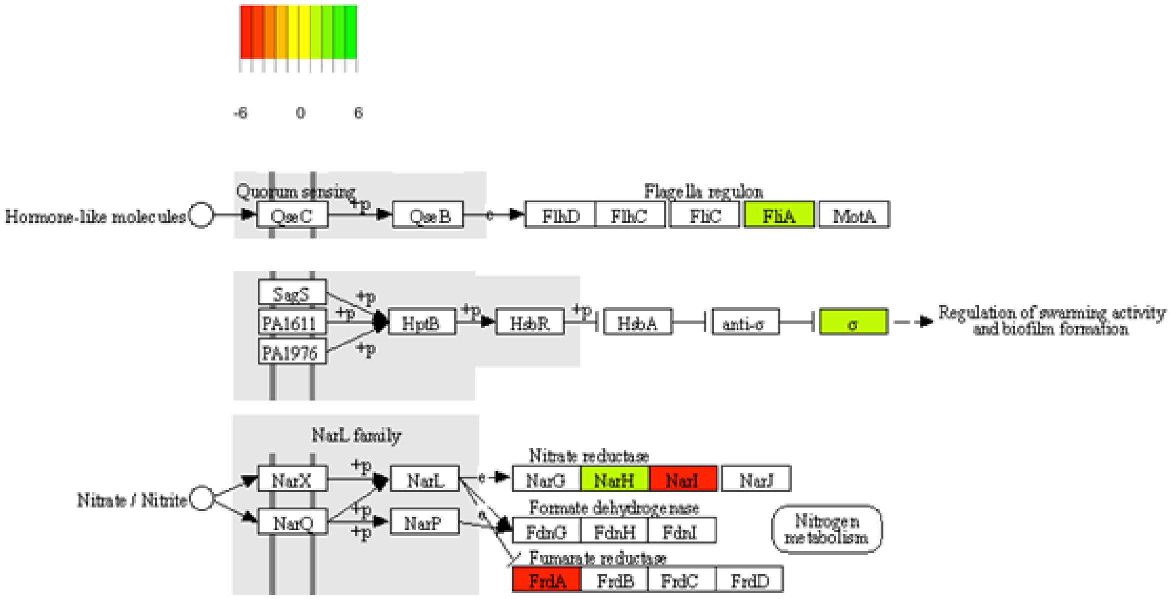


**Figure S12│Swarming associated genes in fecal metatranscriptome. a-b,** 8-week old mice were exposed to DSS water and treated with either heat-killed SM3 (HK) or SM3 for 12 days. **a,** Metatranscriptome analysis of feces collected on Day 0, 4 and 12 showed steady increase in abundance of lauroyl acyltransferase involved in Lipid A biosynthesis on day 12 in SM3 treated mice when compared to Day 0 in conjunction with mucosal healing. HK showed decrease in abundance of lauroyl acyltransferase on Day 12, in conjunction with mucosal inflammation. **b,** Relative abundance of swarming associated transcripts identified in SM3 treated mice when compared to HK. Pathway rendered using KEGG Pathways.

### Figure S13

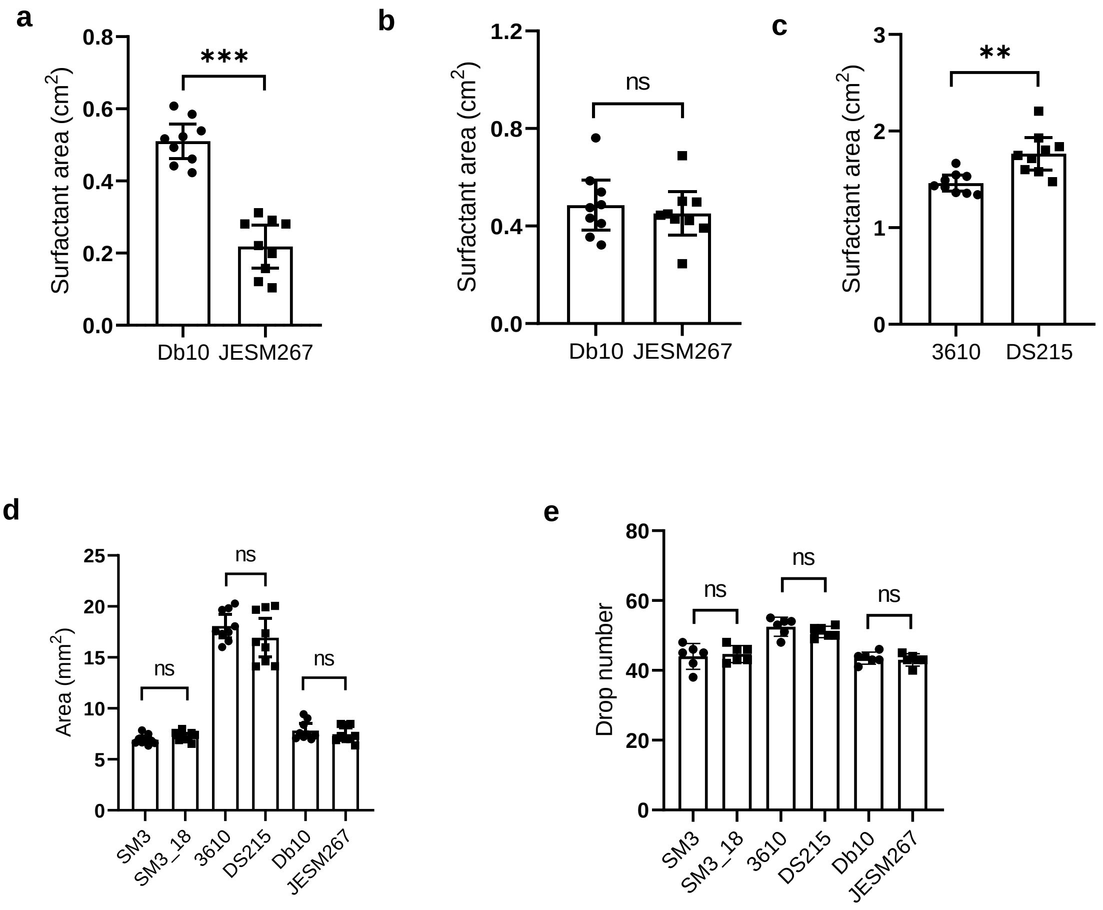


**Figure S13│Surfactant production by swarming bacteria. a-b,** Overnight bacterial cultures of wildtype *S. marcescens* Db10 and its non-swarmer mutant JESM267 spotted on 2.5% blood agar and incubated at 30°C (n = 3, each in triplicate) **(a)**, and 37°C (n = 3, each in triplicate) **(b)**, for 36-48 hours. **c,** Identical method was used to estimate surfactin production by swarmer strain of *Bacillus subtilis* 3610 and its non-swarmer mutant DS215 at 37°C for 36-48 hours (n = 3, each in triplicate). **d,** Cross-sectional area of bacteria supernatant droplets (5 μL) on 96-wellpolystyrene plate lid (n = 3, each in triplicate). **e,** Drop numbers of bacteria supernatant droplets dropping from a glass Pasteur pipet to refill the volume of 1 mL (n =3, each in duplicate). Data are represented as mean and 95% CI, and significance tested using a two-tailed Student's t-test. $^{**}P < 0.01$; $^{***}P < 0.001$; ns, not significant; CI, Confidence Interval.

**Figure S14**

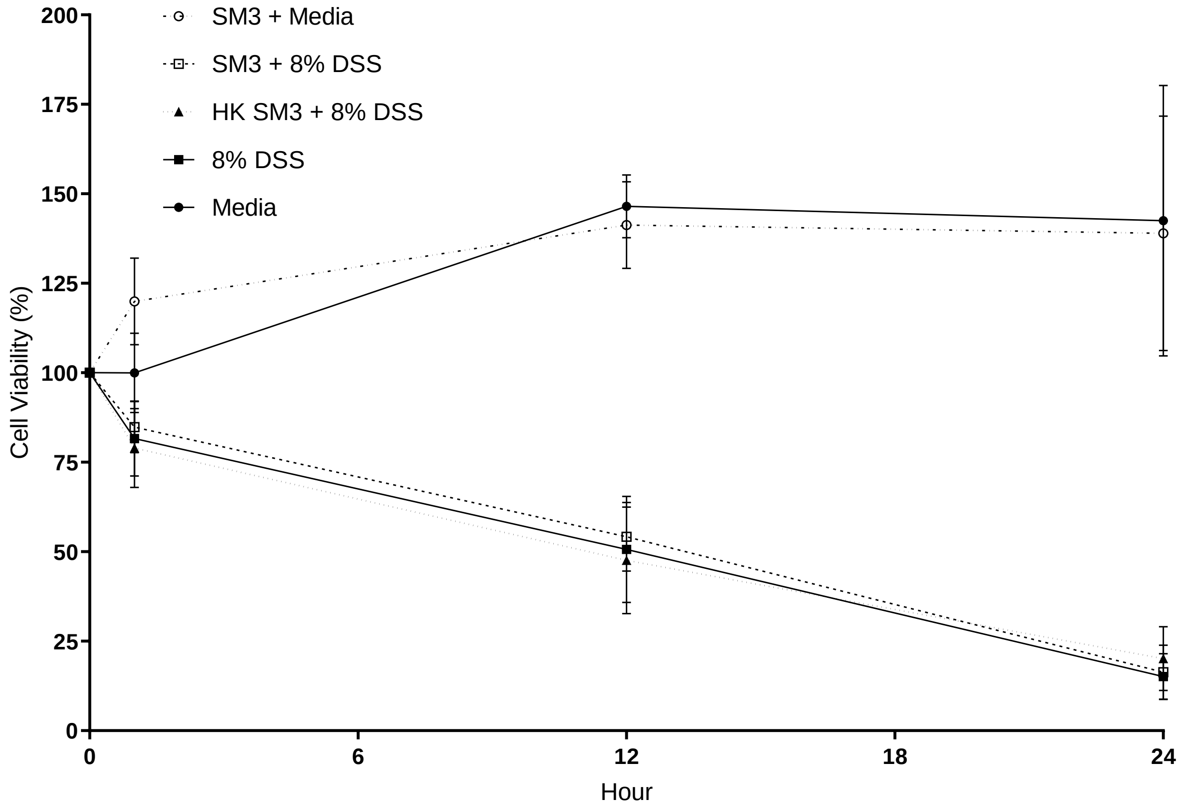


**Figure S14│Effect of DSS on Caco-2 cell line cytotoxicity.** Data represent % survival of Caco-2 cells treated in each tested condition (SM3+8% DSS, HK SM3+8% DSS and 8% DSS) compared to media only at each time point (0, 12 and 24 hours). MTT assay (n = 3, each in triplicate) using treated cell lines was performed to quantify cell survival. Data are represented as mean and 95% CI. DSS, Dextran Sulphate Sodium; HK, Heat Killed.

### Figure S15

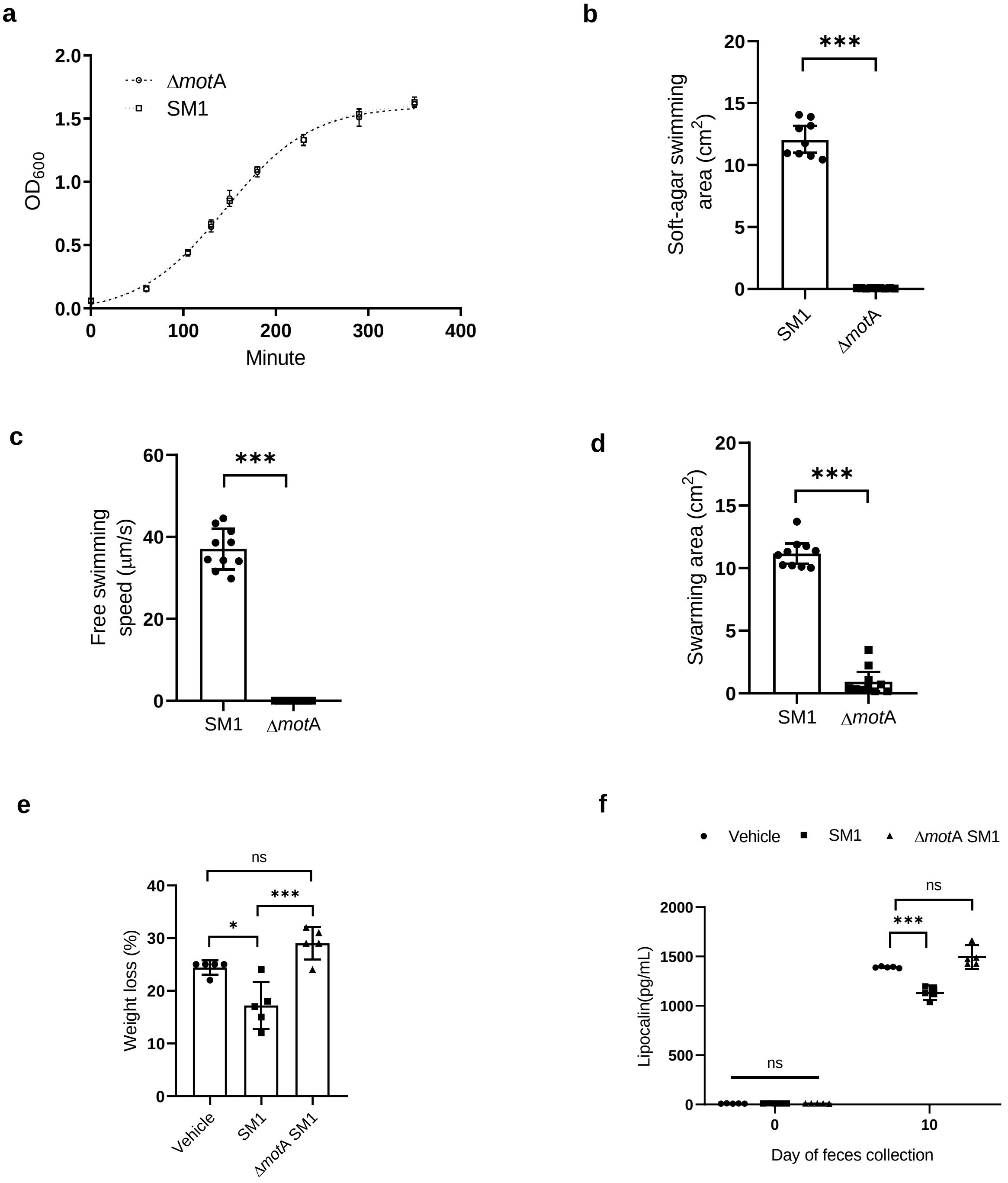


**Figure S15│Characterization of non-motile Δ*mot*A *Enterobacter* sp. SM1. a-d,** indicates comparison of growth curve (n = 3, independent experiment) **(a)**, swimming ability in 0.3% LB agar (n = 3, each in triplicate) **(b)**, swimming in liquid medium(n = 5, each in duplicate) **(c)**, and swarming ability (n = 5, each in duplicate) **(d)** on 0.5% LB agar, between wildtype and the mutant strain. Swimming in agar and swarming comparison between SM1 and Δ*mot*A was performed on the same plate in each experiment. **e-f**, 8-week old mice were exposed to DSS water and treated with vehicle (LB), *Enterobacter sp*. SM1 (SM1) or Δ*mot*A *Enterobacter sp*. SM1 (Δ*mot*A SM1) by oral gavage for 10 days. **e,** Weight loss (n = 5) per treatment group). **f**, Biochemical assessment of lipocalin in feces collected from mice on day 0 and 10 (n = 5) per treatment group. **a-f**, Data are represented as mean and 95% CI. **b-d**, significance tested using a two-tailed Student's t-test. **e-f**, significance tested using one-way ANOVA followed by Tukey's post hoc test.

### Figure S16

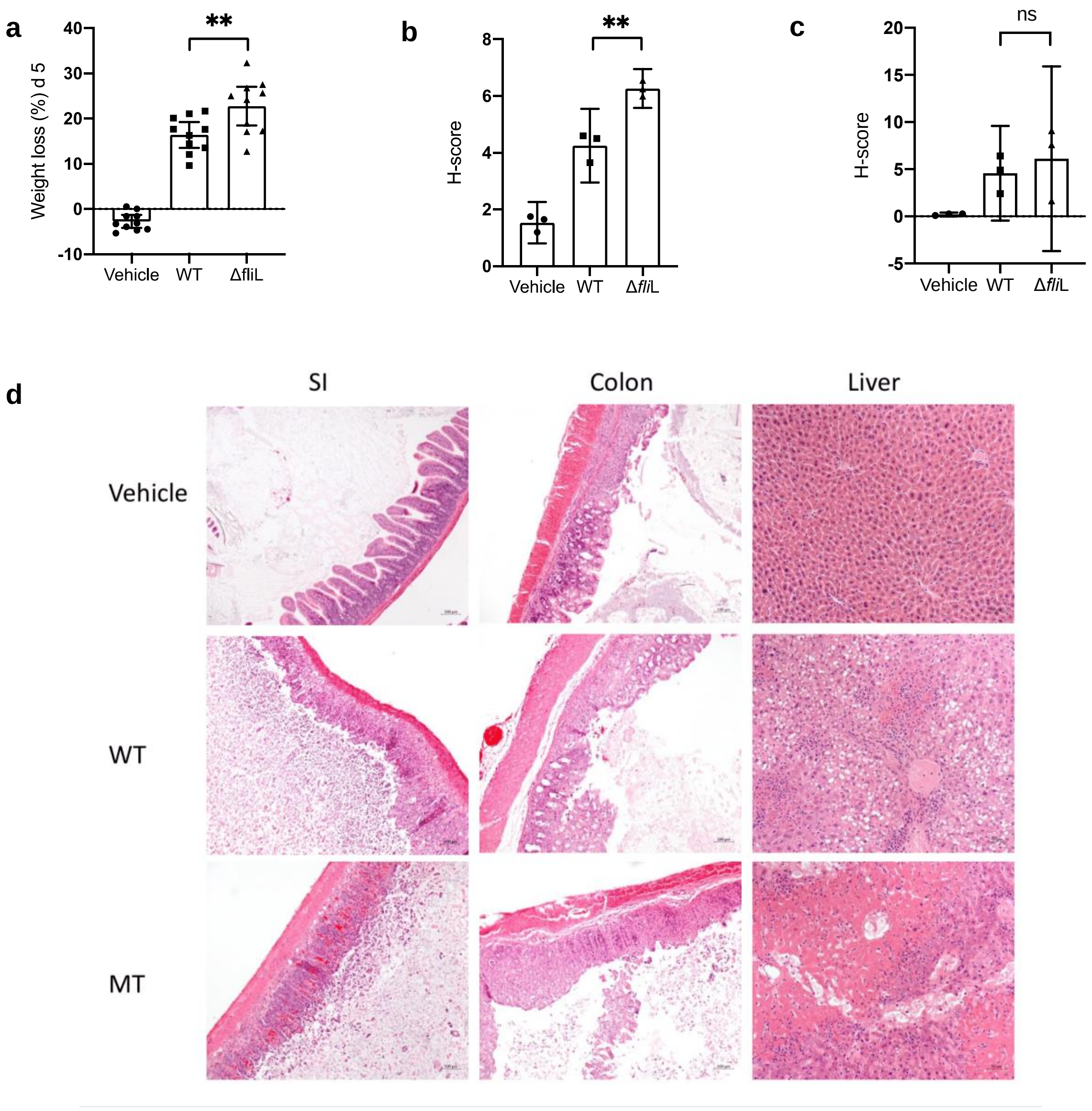


**Figure S16 │ Effects *Salmonella enterica* serovar Typhimurium on DSS-induced colitis in C57BL/6 mice.** C57BL/6 mice (8-week old) were exposed to DSS water and treated with vehicle (LB), *S. enterica* serovar Typhimurium or its Δ*fli*L mutants by oral gavage for 5 days. **a-c** indicates the weight loss (**a**) (n = 10 per treatment group), H-score for colon (**b**) and small intestine (**c**) (n = 3 per treatment group). **d,** Representative images of H&E stained different organs (100x). Data are represented as mean and 95% CI, and significance tested using one-way ANOVA followed by Tukey's post hoc test. WT, wild type *S. enterica* serovar Typhimurium; MT, mutant strain Δ*fli*L; SI, small intestine.

**Supplementary Table 1. Genomic difference between *Enterobacter sp.* SM1 and SM3**

| Mutations / SNP's | Mismatch Nucleotide position (from Start Codon) | Gene Function | Synonymous / Non-synonymous |
|---|---|---|---|
| SM1 ctgaatggcagggaaaccgttgtcgatgac<br>SM3 ctgaatggcagggaaaccgttgtcgacgac | 207 bp | Hypothetical protein | Synonymous |
| SM1 ggtgttgaggttaataacctcagcaat<br>SM3 ggcgataaggttaataaccttgtcgat | 456, 458, 460, 474, 475,476, 478 | 16S rDNA | - |
| SM1 ggttgttcccttgaggagtg<br>SM3 ggttgtgcccttgaggcgtg | 839, 849 | 16S rDNA | - |
| SM1 gcgtggtattcaggcggcaacggctgcgcttcaggggctggct<br>SM3 gcgcggtattcaggcggcaacggcggcgcttcaggggcttgcg | 3177, 3198, 3213, 3216 | Putative large exoprotein involved in heme utilization or adhesion of ShlA/HecA/FhaA family | Synonymous |
| SM1 cccgccatgagggaaaggc<br>SM3 cccgccatgaggtgaaaggc | 713 | Fructose-1,6-bisphosphatase, GlpX type (EC 3.1.3.11) | Non-synonymous Gly (SM1) → Val (SM3) |
| SM1 gggggatcttctttatc<br>SM3 ggtgggatcttctttatc | 183 | Uncharacterized MFS-type transporter | Synonymous |
| SM1 gtgatgaaggcgagcatg<br>SM3 gtgatgaaggcgcgcatg | 73 | Low-affinity gluconate/H+ symporter GntU | Non-synonymous Ser (SM1)→ Arg (SM3) |

*SNP's and mutations between SM1 and SM3 are highlighted yellow

**Supplementary Videos**

**Video S1. Time lapse video of swarming *Enterobacter* sp. SM1, SM2 and SM3.** A .mp4 file of Supplementary Video S1 | Five (5) μL of *Enterobacter* sp. SM1, SM2, and SM3 overnight culture were inoculated on 0.5% LB agar (labeled as SM1, SM2, and SM3 respectively). The plate was incubated in 37°C chamber for 10 hours and time lapse photos were taken. Image contrast was enhanced using Adobe Photoshop CC 2017 and the images were rendered to a video file at 8 frames per second (fps).

**Video S2. Representative video of *S. marcescens* race assay on normal mucosal surface.** A .mp4 file of Supplementary Video S2 | Two pieces of normal C57BL/6 mouse colon tissue were placed on 1.5% agar bordering on 0.5% LB agar. Overnight culture of *S. marcescens* Db10 (marked as WT) and its mutant JESM267 (marked as JES) were diluted $10^{12}$ times and inoculated on the far end of the tissue (marked as ticks) respectively. The plate was incubated in 37°C chamber for 10 hours and time lapse photos were taken. Images were rendered to video using ImageJ at 10 fps.

**Video S3. Representative video of *S. marcescens* race assay on colitic mucosal surface.** A .mp4 file of Supplementary Video S3 | Two pieces of colitic C57BL/6 mouse colon tissue were placed on 1.5% agar bordering on 0.5% LB agar. Overnight culture of *S. marcescens* Db10 (marked as WT) and its mutant JESM267 (marked as JES) were diluted $10^{12}$ times and inoculated on the far end of the tissue (marked as ticks) respectively. The plate was incubated in 37°C chamber for 10 hours and time lapse photos were taken. Images were rendered to video using ImageJ at 10 fps.

**Video S4. Time lapse video of Δ*mot*A *Enterobacter* sp. SM1 and LB only on normal and colitic mucosal surface.** A .mp4 file of Supplementary Video S4 | Normal and colitic mouse colon tissue were placed on 1.5% agar bordering on 0.5% LB agar. LB and Δ*mot*A SM1 overnight culture diluted $10^{12}$ were inoculated on the tissues far end, respectively. The plate was incubated in 37°C chamber for 20 hours and time lapse photos were taken. Images were rendered to video file using ImageJ at 20 fps.

#### Supplementary References


Bodour, A.A., and Miller-Maier, R.M. (1998). Application of a modified drop-collapse technique for surfactant quantitation and screening of biosurfactant-producing microorganisms. Journal of Microbiological Methods *32*, 273-280.

Darling, A.E., Mau, B., and Perna, N.T. (2010). progressiveMauve: multiple genome alignment with gene gain, loss and rearrangement. PLoS One *5*, e11147.

Dilmohamud, B.A., Seeneevassen, J., Rughooputh, S.D.D.V., and Ramasami, P. (2005). Surface tension and related thermodynamic parameters of alcohols using the Traube stalagmometer. European Journal of Physics *26*, 1079-1084.

Kearns, D.B. (2010). A field guide to bacterial swarming motility. Nat Rev Microbiol *8*, 634-644.

Kearns, D.B., Chu, F., Rudner, R., and Losick, R. (2004). Genes governing swarming in Bacillus subtilis and evidence for a phase variation mechanism controlling surface motility. Mol Microbiol *52*, 357-369.

Kearns, D.B., and Losick, R. (2003). Swarming motility in undomesticated Bacillus subtilis. Mol Microbiol *49*, 581-590.

Newsom, D.M., Bolgos, G.L., Colby, L., and Nemzek, J.A. (2004). Comparison of body surface temperature measurement and conventional methods for measuring temperature in the mouse. Contemp Top Lab Anim Sci *43*, 13-18.

Pradel, E., Zhang, Y., Pujol, N., Matsuyama, T., Bargmann, C.I., and Ewbank, J.J. (2007). Detection and avoidance of a natural product from the pathogenic bacterium Serratia marcescens by Caenorhabditis elegans. Proc Natl Acad Sci U S A *104*, 2295-2300.

Seemann, T. (2014). Prokka: rapid prokaryotic genome annotation. Bioinformatics *30*, 2068-2069.

Selvam, R., Maheswari, P., Kavitha, P., Ravichandran, M., Sas, B., and Ramchand, C.N. (2009). Effect of Bacillus subtilis PB6, a natural probiotic on colon mucosal inflammation and plasma cytokines levels in inflammatory bowel disease. Indian J Biochem Biophys *46*, 79-85.

Yoon, S.H., Ha, S.M., Lim, J., Kwon, S., and Chun, J. (2017). A large-scale evaluation of algorithms to calculate average nucleotide identity. Antonie Van Leeuwenhoek *110*, 1281-1286.
